# Linking local and large-scale salient events with oscillatory and broadband arrhythmic activities in the resting human brain

**DOI:** 10.1101/2024.02.28.582552

**Authors:** Damián Dellavale, Emahnuel Troisi Lopez, Antonella Romano, Giovanni Rabuffo, Pierpaolo Sorrentino

## Abstract

**Objective:** Narrowband oscillations (NOs) and Broadband Arrhythmic Activity (BAA) are valuable conceptualizations extensively used to interpret brain data, with NOs linked to communication and synchronization and BAA encompassing scale-free dynamics and neuronal avalanches. Although both frameworks offer critical insights into brain function, they have largely evolved in parallel, with limited integration and no unifying mechanistic account of how these dynamics interact to generate transient, Salient Events (SEs). This gap is particularly pressing given recent interest in how SEs—brief (≈ 100 ms) bursts of activity coordinated across brain regions—relate to large-scale brain function and cognition. To address this, we introduce a signal-level framework that links the Fourier spectral properties (oscillation-domain) of neural signals to the emergence of realistic SEs in the time-domain from NOs and BAA.

**Methods:** Our approach is grounded in a novel concept—Spectral Group Delay Consistency (SGDC)—along with associated measures that quantify the temporal alignment of spectral components and capture the conditions under which Nos and BAA coalesce into transient, burst-like events. Unlike traditional power- or phase-based measures, or higher-order statistical metrics such as kurtosis and cokurtosis, SGDC provides a signal-level mechanistic account of how local and large-scale SEs emerge from the spectral structure of the underlying signals. Empirical validation is provided using source-reconstructed MEG data from a large cohort and a comprehensive array of features characterizing the statistical, spatiotemporal and spectral properties of observed SEs.

**Results:** We found that the SEs identified in our empirical MEG dataset can be segregated based on their spectral signature in two main groups having different propagation patterns. Using generative models based on the SGDC mechanism we provide a theoretical framework to interpret these experimental results showing that cluster 2 events are specifically related to the long-range spread of narrowband alpha bursts across the brain network (i.e., SNEs: Salient Network Events), whereas cluster 1 events correspond to more short-lived and spatially localized fluctuations mainly promoted by the BAA (i.e., SLEs: Salient Local Events). We also provide analytical arguments and numerical simulations showing that a) high SGDC in specific narrow frequency bands, b) transient cross-regional coherent NOs and c) BAA, are all key ingredients for the emergence of realistic SNEs.

**Significance:** We combine experimental evidence supported by a signal-level analytical framework and numerical simulations based on generative models to demonstrate that transient phase-structured alpha bursts, shaped by the SGDC mechanism, contribute to long-range coordination during rest. This extends the communication-through-coherence hypothesis to the transient domain. Additionally, SGDC links to findings that NOs interact with fast microstates (≈100 - 200 ms) and may modulate long-range dependencies across timescales. While previous studies have described SEs within the framework of neuronal avalanches, they often lacked a generative, signal-level account. Here, we bridge that divide by offering a mathematically grounded and empirically validated framework that accounts for oscillatory and aperiodic bursts perspectives on brain activity.

**Highlights:**

- Salient network events propagating across the brain during spontaneous resting state activity, are highly structured in terms of their spatial, temporal and spectral properties.
- The spectral group delay consistency framework provides a signal-level mechanism that accounts for transient salient network events emerging from oscillatory components of the brain activity.
- Narrowband oscillations and broadband arrhythmic activity interact to shape the timing and spatial extent of salient events.
- Spectral group delay consistency, transient cross-regional coherent narrowband oscillations and broadband arrhythmic activity, are all key ingredients for the emergence of realistic salient network events.
- Salient network events during rest reflect large-scale spreading of synchronized alpha band activity, which may play a functional role as a long-range interaction mechanism in the human brain.

## 1. INTRODUCTION

The human brain generates complex behaviors from the coordinated interaction of neuronal populations, with evidence showing different degrees of specialization/distribution of these networks. Such coordination is accompanied (or driven) by neural activity patterns that can be measured using techniques like electroencephalography (EEG) or magnetoencephalography (MEG). In general, electromagnetic brain signals are characterized by both narrowband rhythmic (i.e., oscillations) and broadband arrhythmic (e.g., 1*/f* scaling in power spectra) components [27].

Oscillatory neural activity comprises rhythmic, periodic fluctuations around a central value in the brain’s signals, which occur across various narrow frequency bands and have been associated with specific cognitive functions [25]. For example, the alpha rhythm, typically between 8-13 Hz, emerges during eyes-closed wakefulness [10, 1, 47], while the gamma rhythm, exceeding 30 Hz, has been proposed to play a role in higher cognitive processes[12]. These oscillatory components manifest as “bumps” in the signals’ Power Spectral Density (PSD).

In contrast to brain oscillations, broadband arrhythmic neural activity exhibits a more complex and irregular nature, often associated with scale-free dynamics (i.e., no characteristic temporal scale) [27]. It generally, but not exclusively, displays a 1*/f*^*β*^ decay pattern in the PSD featuring a fractal-like distribution of power across frequencies, with *β* spanning a range of values depending on the brain condition, frequency range and recording modality (roughly 0.1 ≲ *β* ≲5). This broadband activity contributes significantly to the brain’s overall signal and is intricately linked with cognitive processes, potentially carrying valuable information [43]. Traditionally, the study of brain *Narrowband Oscillations* (NOs) and *Broadband Arrhythmic Activity* (BAA) has provided two lenses through which electrophysiological data have been examined [24]. In general, spectral (i.e., oscillation-domain) attributes like power and phase offer rich insights into brain dynamics, enabling the discrimination of brain activity during perceptual tasks and distinguishing between healthy and pathological dynamics in resting states [30]. For instance, the literature on brain connectivity has traditionally diverged into two primary streams: one emphasizing NOs—rhythmic, frequency-specific activity linked to communication and synchronization [56, 18, 15, 29, 38]—and another focused on BAA, encompassing scale-free dynamics such as 1*/f* activity and neuronal avalanches [71, 5, 48]. Although both frameworks offer critical insights into brain function, they have largely evolved in parallel, with limited integration and no unifying mechanistic account of how these dynamics interact to generate transient, *Salient Events* (SEs). This gap is particularly pressing given recent interest in how SEs—brief (≈ 100 ms) bursts of activity coordinated across brain region—relate to large-scale brain function and cognition [46, 49, 36, 70, 2, 40, 37, 35].

Besides NOs and BAA, the analysis of collective brain dynamics reveals that system-level neuronal activity is interspersed by two types of SEs: *Salient Local Events* (SLEs) and *Salient Network Events* (SNEs). During SNEs, subsets of brain regions collectively exhibit rare fluctuations above a threshold (e.g., signal amplitude *>* 3 standard deviations), igniting from specific brain sites, propagating across the brain circuitry in an avalanche-like cascade of activations, and finally decaying below the threshold. As an example, Fig. C.1 in Appendix C shows a SE observed in our MEG dataset, constituted by transient above-threshold fluctuations overlapped (i.e., disclosing time-overlap, coordinated) across 5 brain regions. Due to the fact that the SE shown in Fig. C.1 involves the activation of more than 1 brain region is named *Salient Network Event* (SNE). On the other hand, a local transient above-threshold fluctuation involving the activation of just 1 brain region is named *Salient Local Event* (SLE). SNEs occur aperiodically and are consistently observed across imaging modalities, including multielectrode array recordings [7, 8], EEG [46, 20], MEG [46, 60], SEEG [52, 73], fMRI [64], and calcium imaging [72, 13]. In particular, SNEs have drawn considerable interest due to their potential significance in information processing [59, 58], facilitating responses with a wide dynamical range [31, 34], and playing a role in achieving flexible dynamics [67, 62, 61]. A specific subtype of SNEs are known as *neuronal avalanches*. The latter were largely studied in the context of the *critical brain hypothesis*, which posits that the brain might be operating near a critical point (i.e., at the edge of a phase transition). In fact, neuronal avalanches display hallmark properties expected in systems that self-organize at a critical point, such as the power-law distribution of avalanche durations (life span) and sizes (number of regions recruited) [46, 60, 41]. However, previous studies raised concerns about the interpretation of power law statistics associated with neuronal avalanches. First, power law distributed avalanches have been found in stochastic noncritical systems (see [16] and references therein). These works highlight the fact that power-law distributions are not unique to systems near a critical point or a phase transition and can be generated by other mechanisms [44]. Second, several factors can contribute to deviations from power-law statistics such as finite size effects (size of the neuronal network or the sampling region) and thresholding procedures used for avalanche detection [69, 68]. Third and more crucially, the neuronal avalanches statistics can be influenced by heterogeneous factors like network interaction/synchronization, the concomitant presence of oscillations and/or other type of SEs (e.g. IEDs: Interictal Epileptiform Discharges) and also external interventions (e.g., antiseizure medications) [42].

The signal processing tools proposed in this work can be used to study a variety of SLEs and SNEs: sleep spindles and K-complexes observed during non-rapid-eye-movement sleep, IEDs and Spike and Wave Discharges (SWDs) associated with epileptogenicity [19] and Paroxysmal Slow-Wave Events (PSWEs) observed in epilepsy and age related neuropathology (e.g., Alzheimer’s disease) [50, 39]. In general, these SEs do not follow power law statistics, indeed, IEDs, SWDs and PSWEs have been observed in a wide range of dynamical regimes associated with clinical and subclinical brain states (see for instance Fig. 5 in [42]). Thus, for the sake of generality, we focus our analysis on the relationship among NOs, BAA, and SEs, without implying a connection to power law distributed neuronal avalanches nor the brain criticality hypothesis.

NOs, BAA, and SEs offer valuable conceptualizations to interpret brain data, however, these well-established perspectives have mainly progressed in parallel, with only limited literature linking them largely restricted to the context of neuronal avalanches [46, 49, 36, 2, 40, 37, 35]. Given the ubiquitous and concurrent presence of NOs, BAA, and SEs in the brain during rest, a fundamental question arises: Can we establish a connection between these perspectives? In other words, can we invoke a parsimonious explanation that justifies the simultaneous presence of these phenomena? To address this, we introduce a signal-level framework that links the Fourier spectral properties (oscillation-domain) of neural signals to the emergence of realistic SEs in the time-domain from rhythmic and broadband aperiodic dynamics. Our approach is grounded in a novel concept— Spectral Group Delay Consistency (SGDC)—along with associated measures that quantify the temporal alignment of spectral components and capture the conditions under which NOs and BAA coalesce into transient, burst-like events. Unlike traditional power- or phase-based measures, or higher-order statistical metrics such as kurtosis and cokurtosis, SGDC provides a signal-level mechanistic account of how local (SLEs) and large-scale (SNEs) salient events emerge from the spectral structure of the underlying signals.

While previous works primarily focused on describing the *interaction* between neuronal avalanches and NOs, in this work we adopts a bottom-up approach, using generative models based on the SGDC mechanism, aimed at elucidating how local and large-scale SEs *emerge* from the oscillatory and broadband arrhythmic components of the brain activity. The proposed data analysis tools are supported by a signal-level analytical SGDC framework designed to be applicable across a variety of (bio)physical domains, regardless of the specific details of the underlying system.

In addition, empirical validation is provided using source-reconstructed MEG data from a large cohort, demonstrating that transient phase-structured alpha bursts, shaped by the SGDC mechanism, contribute to long-range coordination during rest. This extends the communication-through-coherence (CTC) hypothesis, according to which neuronal information is transferred via phase alignment (coherence) of rhythmic activity [22, 23], to the transient domain. Additionally, SGDC links to findings that NOs interact with fast microstates (≈ 100 - 200 ms) [3, 6, 66, 48] and may modulate long-range dependencies across timescales [5]. Thus, while previous studies have described SEs within the framework of neuronal avalanches, they often lacked a generative, signal-level account. Here, we bridge that divide by offering a mathematically grounded and empirically validated framework that accounts for oscillatory and aperiodic bursts perspectives on brain activity.

## 2. METHODS

### 2.1. Participants and data

We analyzed a source-reconstructed MEG dataset previously published in [63, 62]. In short, 58 young adults (32 males/26 females, mean age *±*SD was 30.72 *±* 11.58) were recruited from the general community. All participants were right-handed and native Italian speakers. The inclusion criteria were (1) no major internal, neurological, or psychiatric illnesses; and (2) no use of drugs or medication that could interfere with MEG/MRI signals. The study complied with the Declaration of Helsinki and was approved by the local Ethics Committee. All participants gave written informed consent. The details regarding the MRI acquisition are described in Section [63]. All technical details in connection with the MEG device are reported in [54]. MEG pre-processing and source reconstruction were performed as in [63, 62]. Briefly, the MEG registration was divided into two eyes-closed segments of 3:30 min each. To identify the position of the head, four anatomical points and four position coils were digitized. Electrocardiogram (ECG) and electro-oculogram (EOG) signals were also recorded. The MEG signals, after an anti-aliasing filter, were acquired at 1024 Hz, then a fourth-order Butterworth IIR band-pass filter in the 0.5-48 Hz band was applied. Principal component analysis was used to remove environmental noise measured by reference magnetometers. Supervised independent component analysis was adopted to clean the data from physiological artefacts, such as eye blinking (if present) and heart activity (generally one component). Noisy channels were identified and removed manually by an expert rater (136 *±* 4 sensors were kept). After this pre-processing, 47 subjects were selected for this work and all further analyses were conducted on traces of 1 min in duration source-reconstructed to 84 brain Regions Of Interest (ROI) based on the Desikan-Killiany-Tourville (DKT) anatomical parcellation atlas (see brain topographies in Figs. 1 and C.3).

**Figure 1:**
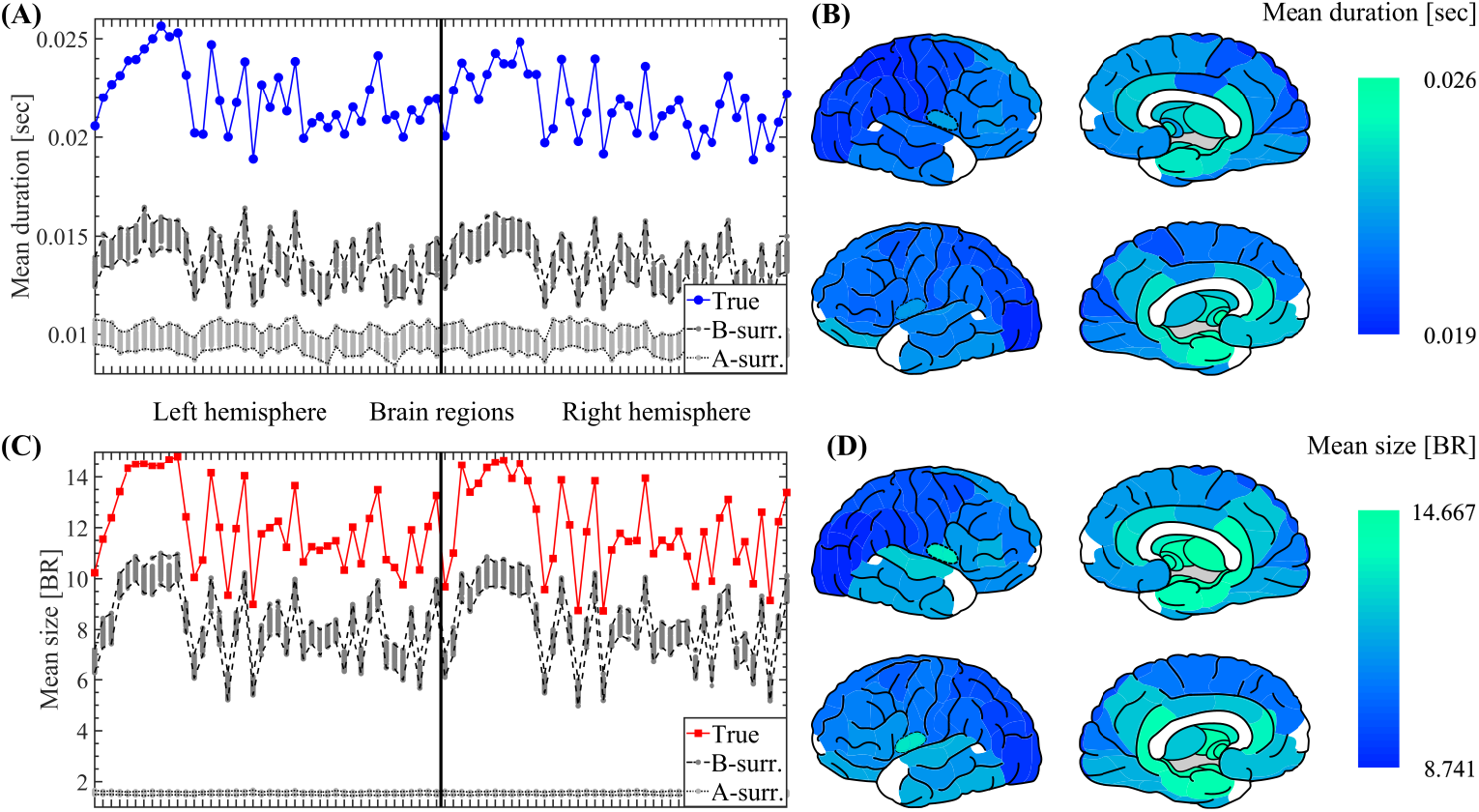
Spatiotemporal characterization of SEs. (A) Spatial profile showing the mean duration of SEs propagating through each brain region (mean value across the 47 participants, see Section 2.4 in Methods). The mean event duration is shown for the MEG data together with the 100 A- and B-surrogates (see Section 2.8 in Methods). The labels and ordering of the brain regions are the same as those shown in Fig. C.2. (B) Brain topographies for the mean duration of SEs as shown in panel A. (C) Same as in A for the size of SEs. (D) Same as in B for the size of SEs. Symbols and abbreviations: SEs, Salient Events; BR, Brain Regions.

### 2.2. Salient events detection

To estimate SEs we first detected the local above-threshold fluctuations on the pre-processed and source-reconstructed MEG time series as described in Section 2.1. In each participant, the 1-minute source-reconstructed MEG time series of each brain region were individually z-scored. Positive and negative excursions beyond a threshold were then identified. The amplitude threshold was set to |*z* |= 3, equivalent to three standard deviations (*±* 3*σ* or equivalently |*z*| = 3). The same amplitude threshold | *z*| = 3 was used in all analyzed brain regions. This procedure was applied separately to all the 47 participants included in the study. An analysis supporting the validity and robustness of using a single amplitude threshold (|*z*| = 3) consistently across all 47 participants is presented in Appendix C.1. Then the SEs duration was assessed by considering that a salient event begins when, in a sequence of contiguous time bins, at least one brain region is active (i.e., above the amplitude activation threshold: |*z*| *>* 3) and ends when all the brain regions are inactive (i.e., below the amplitude activation threshold: |*z*| *≤* 3) [7, 60]. Besides, the SEs size was defined as the total number of brain regions activated during a given event. Note that a salient event involving more than one brain region (i.e., SNE) is associated with a sequence adjacent time bins in which at least one brain region is active (|*z*| *>* 3). Thus, the detection of SNE depends on the time binning of the analyzed time series. Unless otherwise specified, in this study we used a time binning corresponding to 1 time sample per time bin (time binning = 1 ms). This procedure allowed the detection of both SLEs (i.e., SEs of size = 1 brain region) and SNEs (i.e., SEs of size *>* 1 brain region, see Fig. C.1).

### 2.3. Salient events activation and co-activation matrices

For each detected SE, we computed the activation matrix (brain regions *×* time bins) as follows. The source-reconstructed, z-scored and time binned signal were binarized, such that, at any time bin, a brain region exceeding *±*3 was set to 1 (active), and all other regions were set to 0 (inactive, see Figs. 2A,B). For each detected SE, we also computed the co-activation matrix (brain regions *×* brain regions) by assigning 1 to all the brain regions recruited in that particular event. Thus, the diagonal of the co-activation matrix contains 1s in all the brain regions active during a given SE. Besides, summation across rows (or columns) produce, in each brain region, the number of co-activated regions during a given SE (i.e., in terms of graph theory, this is known as the degree of each brain region). The mean co-activation matrix shown in the Fig. 7C was computed by first averaging the co-activation matrices corresponding to all the SEs detected in each subject, and then, averaging the resulting matrix across all the participants.

**Figure 2:**
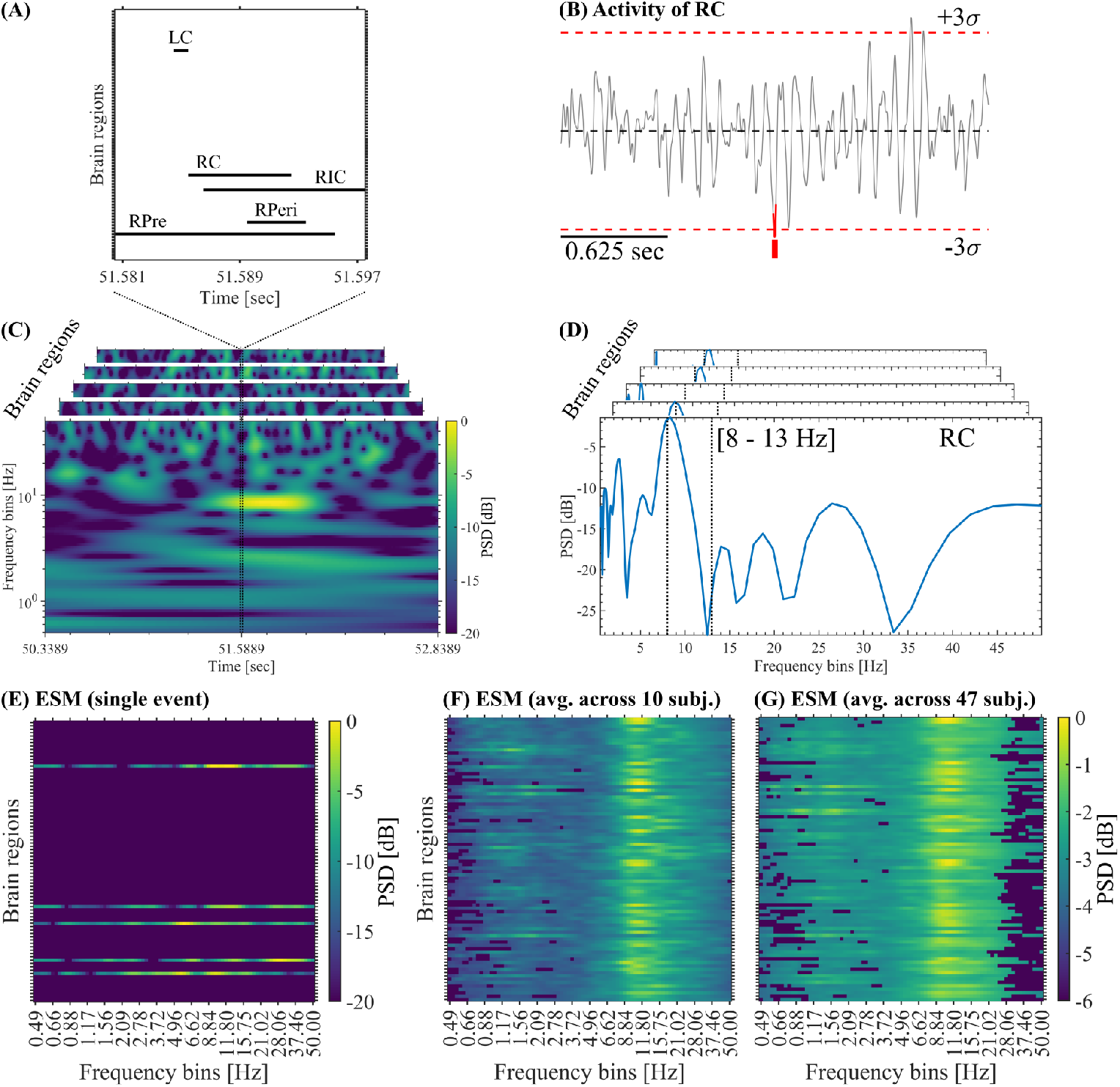
Event Spectral Matrix. (A) Activation matrix of a single SE showing the time intervals in which each brain region was active (i.e., absolute amplitude *>* 3*σ*). (B) Activity of the brain region RC disclosing the above-threshold fluctuation (highlighted in red) associated with the SE shown in panel A. (C) Whitened time-frequency maps of each brain region involved in the SE shown in panel A. (D) Whitened power spectra associated with each brain region involved in the SE shown in panel C. The vertical dotted lines indicate the alpha band. To build the ESM, we average the whitened time-frequency maps selectively across the time samples corresponding to the occurrence of each SE. As a result, we obtain a whitened power spectrum for each brain region (see Section 2.5 in Methods). (E) ESM corresponding to the SE shown in panel C. (F) Mean ESM resulting from the average across all the SEs detected in the 10 subjects, and then, Bonferroni-thresholded using the C-surrogates (see Methods). (G) Mean ESM resulting from the average across all the SEs detected in the 47 subjects, and then, Bonferroni-thresholded using the B-surrogates (see Methods). Symbols and abbreviations: SEs, Salient Events; ESM, Event Spectral Matrix; PSD, Power Spectral Density; RPre, Right Precuneus; RC, Right Cuneus; RPeri, Right Pericalcarine; RIC, Right Isthmus Cingulate; LC, Left Cuneus.

### 2.4. Salient events spatiotemporal profile

To characterize SEs spatiotemporal profile, we introduce two ROI-wise metrics: The *mean event duration* measuring the typical duration of SEs propagating through a brain region; and the *mean event size* measuring the typical size of SEs propagating through a brain region. Specifically, we assign to each brain region the mean event duration (or size) computed on all the SEs recruiting that particular region. The *mean event duration* and *mean event size* profiles shown in the Fig. 1 were computed by first considering all the SEs detected in each subject, and then, averaging the resulting profiles across all the participants.

### 2.5. Event spectral matrix

For the spectral characterization of SEs we introduce the Event Spectral Matrix (ESM). To obtain the ESM we first compute the whitened time-frequency representation on the whole time series of each brain region (see Fig. 2C). Then, the time-frequency maps were selectively averaged across the time points corresponding to the occurrence of the SE of interest. As a result, we obtain a whitened power spectrum corresponding to each brain region recruited by that particular SE (see Fig. 2D). Finally, these power spectra are arranged in a single matrix conforming the ESM (Brain regions *×* Frequency bins, see Fig. 2F). The time-frequency maps were computed as scalograms using Morlet wavelets of duration 2 *g width/*(2*πf*) sec., where *g* = 3 (std. dev.), *width* = 7 (cycles) and *f* ∈ [0.5, 50] Hz. Spectral whitening, via *Z*_*H*0_-score normalization of each frequency bin across time samples as described in [19], was included in the computation of the time-frequency maps to facilitate the visualization of the high-frequency components in the resulting ESM. The ESM can be defined at the single event level (see Fig. 2F), by averaging all the SEs in each subject (data not shown) and by averaging the mean ESM of each subject across participants (see Figs. 2F,G and 3A,B). Of note, the ESM does not represent the frequency content of SEs since the latter are very short-duration transient events, instead, the ESM reveals the spectral signature associated with the oscillatory activity co-occurring with each SE. That is, the ESM reveals the co-occurrence (or coupling) between the oscillatory activity and SEs across brain regions. To assess the statistical significance of the spectral signatures associated with the SEs, we compute pixel-level thresholding on the mean ESM with Bonferroni correction for multiple comparisons. More specifically, we computed the mean ESM on each one of the 100 B- or C-surrogate datasets (see Section 2.8). Then these 100 surrogate mean ESMs were used to compute pixel-level thresholding on the true mean ESM using a Bonferroni-adjusted two-tailed statistical threshold = 0.05*/*(Brain regions *×* Frequency bins). Note that this Bonferroni correction for multiple comparisons assuming independence between adjacent spatial/frequency bins of the mean ESM is a quite conservative test, yet, the observed spectral signature in the alpha band is evident even after this stringent thresholding process (see Fig. 2F,G).

### 2.6. Salient events waveform shape

To characterize the waveform shape of SEs we follow a ROI-wise approach. First, in each brain region we computed the average across the 200 ms signal epochs (absolute value) centered around the start time of the SEs of interest recruiting that particular region (see gray lines in Figs. 3C,D and D.1C,D). Then, we obtained the mean SEs waveform shape by computing the average of the resulting time series across the brain regions (see the red and blue lines in Figs. 3C,D and D.1C,D).

**Figure 3:**
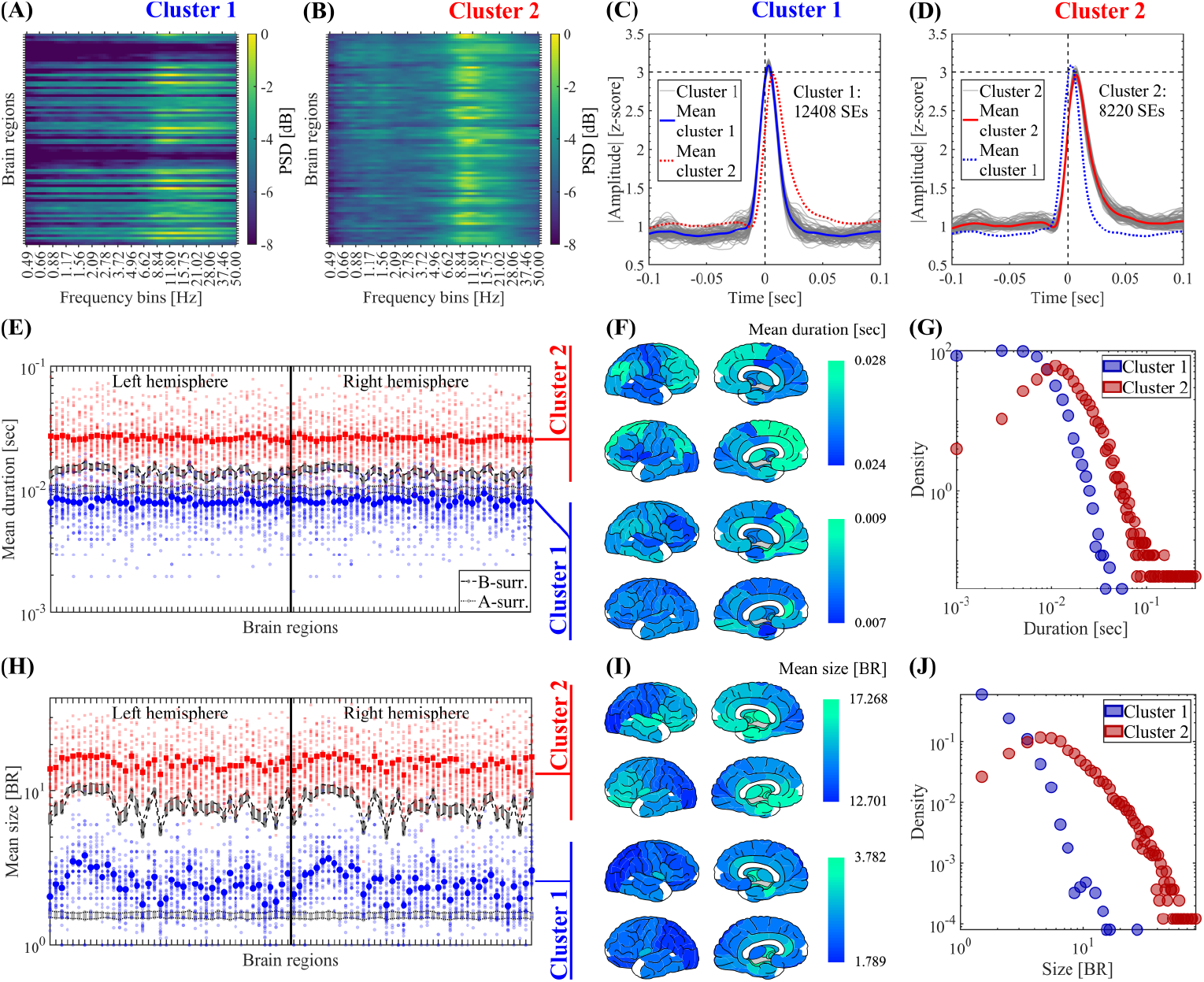
Clustering of SEs according to their spectral signature. The SEs obtained from 41 subjects were clustered using the Louvain algorithm (resolution parameter *γ* = 1, see Methods). (A, B) Mean ESM of the two SE clusters identified by the Louvain algorithm computed on the SEs detected in the 41 participants. (C, D) Waveform shapes of the SEs pertaining to the two SE clusters identified by the Louvain algorithm. Thin gray lines correspond to the average waveform shape in each brain region. Thick blue and red lines correspond to the resulting waveform shape averaged across the brain regions for cluster 1 and 2 SEs, respectively. (E) Spatial profile showing the mean duration of SEs pertaining to cluster 1 (in blue) and cluster 2 (in red). For the true data, the small and big markers correspond to the mean spatial profile in each patient and the average across the 41 participants, respectively (see Methods). The labels and ordering of the brain regions are the same as those shown in Fig. C.2. To test the significance of the difference of the mean SEs duration between cluster 1 and cluster 2, in each brain region we computed a non-parametric permutation test (random sampling without replacement, 1*×* 10^4^ permutations). All the brain regions disclosed a statistically significant difference of the mean SEs duration between cluster 1 and 2 (the Bonferroni-adjusted two-tailed P values result *P <* 0.001 in all the brain regions). (F) Brain topographies for the mean duration of SEs averaged across the 41 participants as shown in panel E. (G) Distribution of the duration of SEs pertaining to the cluster 1 and cluster 2 observed in the 41 participants. (H) Same as in E for the size of SEs. To test the significance of the difference of the mean SEs size between cluster 1 and cluster 2, in each brain region we computed a non-parametric permutation test (random sampling without replacement, 1 *×* 10^4^ permutations). All the brain regions disclosed a statistically significant difference of the mean SEs size between cluster 1 and 2 (the Bonferroni-adjusted two-tailed P values result *P <* 0.001 in all the brain regions). (I) Same as in F for the size of SEs. (J) Same as in G for the size of SEs. Symbols and abbreviations: SEs, Salient Events; ESM, Event Spectral Matrix; BR, Brain Regions.

### 2.7. Salient events propagation modes

To assess the SEs starting modes we assign to each brain region the number of events igniting in that particular site (e.g., see the RPre brain region in the activation matrix shown in Fig. 2A). Similarly, for the SEs ending modes we assign to each brain region the number of events extinguishing in that particular site (e.g., see the RIC brain region in the activation matrix shown in Fig. 2A).

For the SEs maximum recruitment modes we assign to each brain region the number of events involving that particular site during the event maximum size (e.g., see the 4 brain regions active at *Time ≈* 51.591 sec in the activation matrix shown in Fig. 2A). Last, by dividing the event count obtained in each brain region by the total number of processed SEs, we obtained the mean spatial profiles for the starting, maximum recruitment and ending SEs modes as shown in the Figs. C.5, C.6, D.2 and D.3.

### 2.8. Surrogate datasets

We generated phase-randomized A-surrogate datasets, that preserve the PSD in each brain region, while disrupting the phase relationships of the spectral components (both within and between brain regions) [51]. For this, in each brain region we implemented a frequency-domain randomization procedure, which involves taking the Discrete Fourier Transform (DFT) of the time series, adding a random phase-shift in the range [*−π, π*] on each spectral component of the DFT (preserving the odd phase symmetry associated with real signals [14]), and then taking the inverse DFT to obtain the surrogate signal back in the time-domain [17]. The 100 phase-randomized A-surrogate datasets were obtained by applying this procedure 100 times on each brain region independently. In addition, we also generated B-surrogate datasets that randomize the phases similarly to the A-surrogate, but in this case preserving both the regional PSDs and the cross-spectra. For this, we follow a similar procedure as described above with the difference that the same random phase-shift was applied to all the brain regions. This implies that the phase difference between any pair of brain regions in *homologous frequency components* is preserved (i.e., preservation of cross-spectra). This implies to preserve the Pearson’s correlations between brain regions (see Appendix A.1). Note that the B-surrogates destroy the phase relationships only between *non-homologous frequency components*. Finally, we generated 100 C-surrogate sets of SEs by randomizing the starting time of each observed salient event and keeping unaltered all the other properties like the event duration and brain regions recruited in each event.

### 2.9. Clustering of salient events

SEs were clustered according to their spectral signature by using the Louvain method for community detection based on modularity maximization [11, 55]. First, the Matrix of Paired Distance (MPD) was obtained by computing the Euclidean distance between the vectorized ESMs corresponding to the SEs of interest taken in pairs. The resulting MPD (Events *×* Events) was normalized to be in the range [0, 1], and the Adjacency Matrix (AM) was computed as AM = 1 *−* MPD. Then, the Louvain algorithm was repeated 100 times on the AM for resolution parameter values in the range 0.5 ≤ *γ ≤* 2 [11, 57]. Optimization of modularity quality function, based on the maximization of the similarity measure (z-scored Rand index) [57], was achieved for resolution parameter values within the range 0.9 ≲ *γ* ≲ 1.1. Finally, a consensus partition was found from the 100 partitions [33, 4, 21]. For the events detected in our source-reconstructed MEG dataset, the Louvain algorithm consistently identified two SE clusters with significant differences in terms of mean event duration, size and spectral signature in their mean ESM (see Figs. 3, C.11 and D.1).

### 2.10. Spectral group delay consistency measures

In this study, we introduce the pairwise complex baseband representation of band-limited signals (Eqs. A.7 - A.10 and A.13 - A.16) to provide analytical arguments showing that the link between local above-threshold fluctuations and oscillations can be understood in terms of the group delay consistency across the spectral components (i.e., Fourier oscillatory constituents) of the neuronal activity. Specifically, in Appendix A.2 we show that the time-domain representation of any finite-length time series *x*(*t*) (inverse DFT, Eq. A.5) can be re-arranged by grouping the Fourier spectral components *X*(*k*) in non-overlapping adjacent pairs, leading to the pairwise complex baseband representation (Eq. A.7). In this new representation (Eq. A.7), the signal *x*(*t*) is decomposed into a linear superposition of amplitude modulated components, each synthesized from an adjacent pair of spectral components (*X*(2*k*), *X*(2*k* + 1) in Eq. A.7). Crucially, the Eq. A.7 explicitly shows that the group delay is the key spectral feature determining the transient synchronization of the Fourier oscillatory constituents of the signal *x*(*t*) leading to the emergence of SEs (see Eq. A.17 and Figs. 4, A.2, A.3 and A.4). More precisely, the group delay determines the time alignment of the amplitude modulated ocillatory constituents of the signal *x*(*t*) in the pairwise complex baseband representation. Such time alignment promotes transient large-amplitude excursions of the signal (i.e., above-threshold fluctuations). Thus, the Eq. A.7 constitutes a group delay-domain representation of the signal *x*(*t*), which lies in-between and links the time-domain and frequency-domain representations:

**Figure 4:**
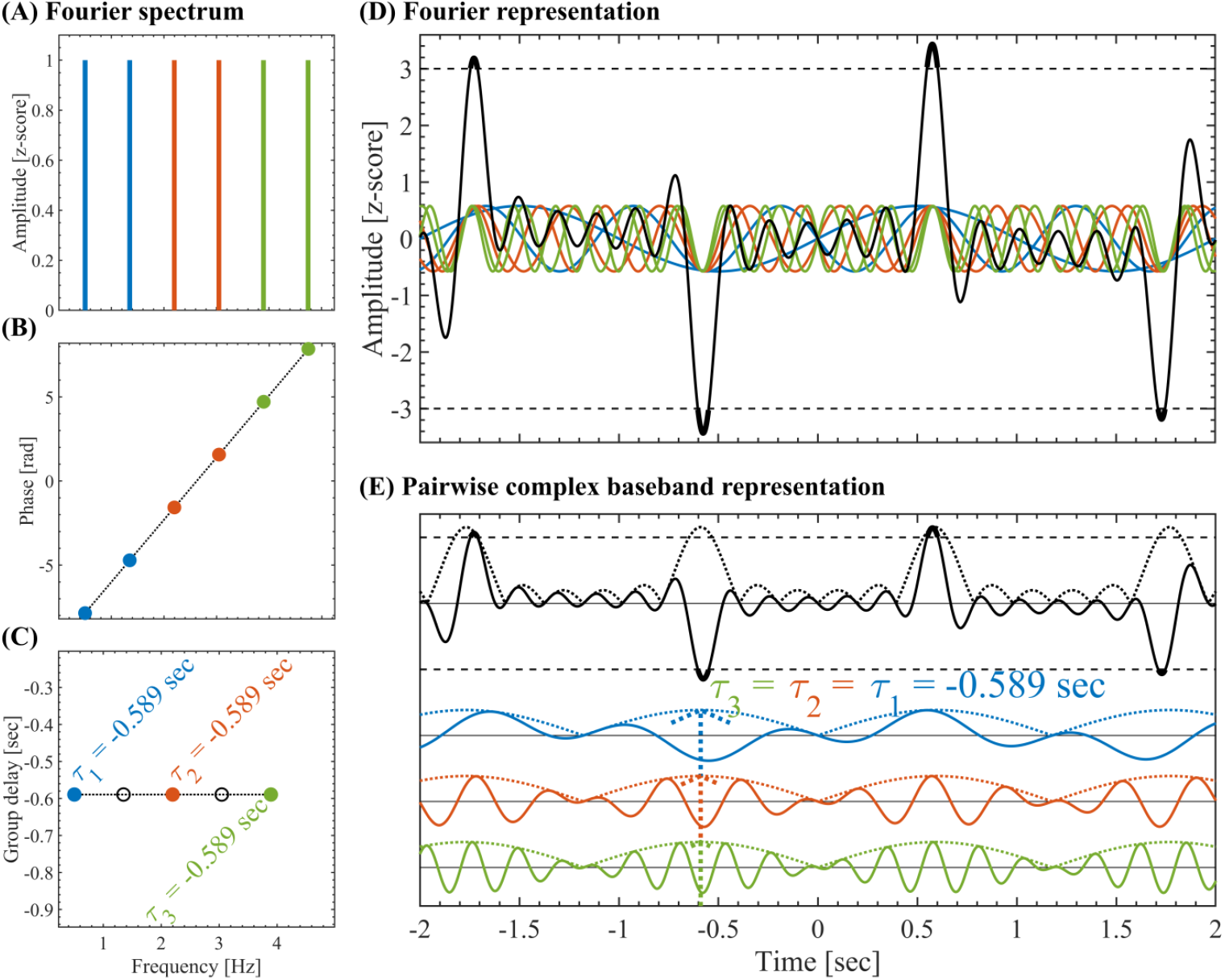
Pairwise complex baseband representation. (A) Set of constant-amplitude *A*(*k*) = 1 oscillatory components uniformly spaced 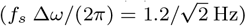 and having non-harmonic frequencies *f*_*s*_ *ω*(*k*)*/*(2*π*) = 0.5 + *k f*_*s*_ Δ*ω/*(2*π*) ∈ [0.5 *−* 5] Hz, where *f*_*s*_ = 1024 Hz is the sampling rate. The pairwise complex baseband representation (Eq. A.13) was obtained by grouping the oscillatory components in adjacent non-overlapping pairs color-coded in blue, red and green. (B) Phases *ϕ*(*k*) having a linear dependence as a function of the frequency within the range *ϕ*(*k*) ∈ 2.5 [−*π, π*]. (C) Group delay *τ* (*k*)*/f*_*s*_ = −Δ*ϕ*(*k*)*/*(*f*_*s*_ Δ*ω*) for the pairs of adjacent oscillatory components. The color-coded filled markers correspond to the *τ* (2*k*)*/f*_*s*_ values, and the black empty markers correspond to *τ* (2*k* + 1)*/f*_*s*_ values (see Eq. A.13). (D) Z-scored signals associated with the Fourier representation. The solid color-coded lines represent the individual oscillatory components, the solid black line is the resulting signal *x*(*t*), the horizontal dashed black lines indicate the threshold at |*z* |= 3. (E) Pairwise complex baseband representation. The solid color-coded lines represent the individual amplitude modulated signals (pairs of adjacent oscillatory components), the solid black line is the resulting signal *x*(*t*), the color-coded and black doted lines are the corresponding amplitude envelopes. For an in depth mathematical description of the pairwise complex baseband representation see Appendix A.2.

**Figure 5:**
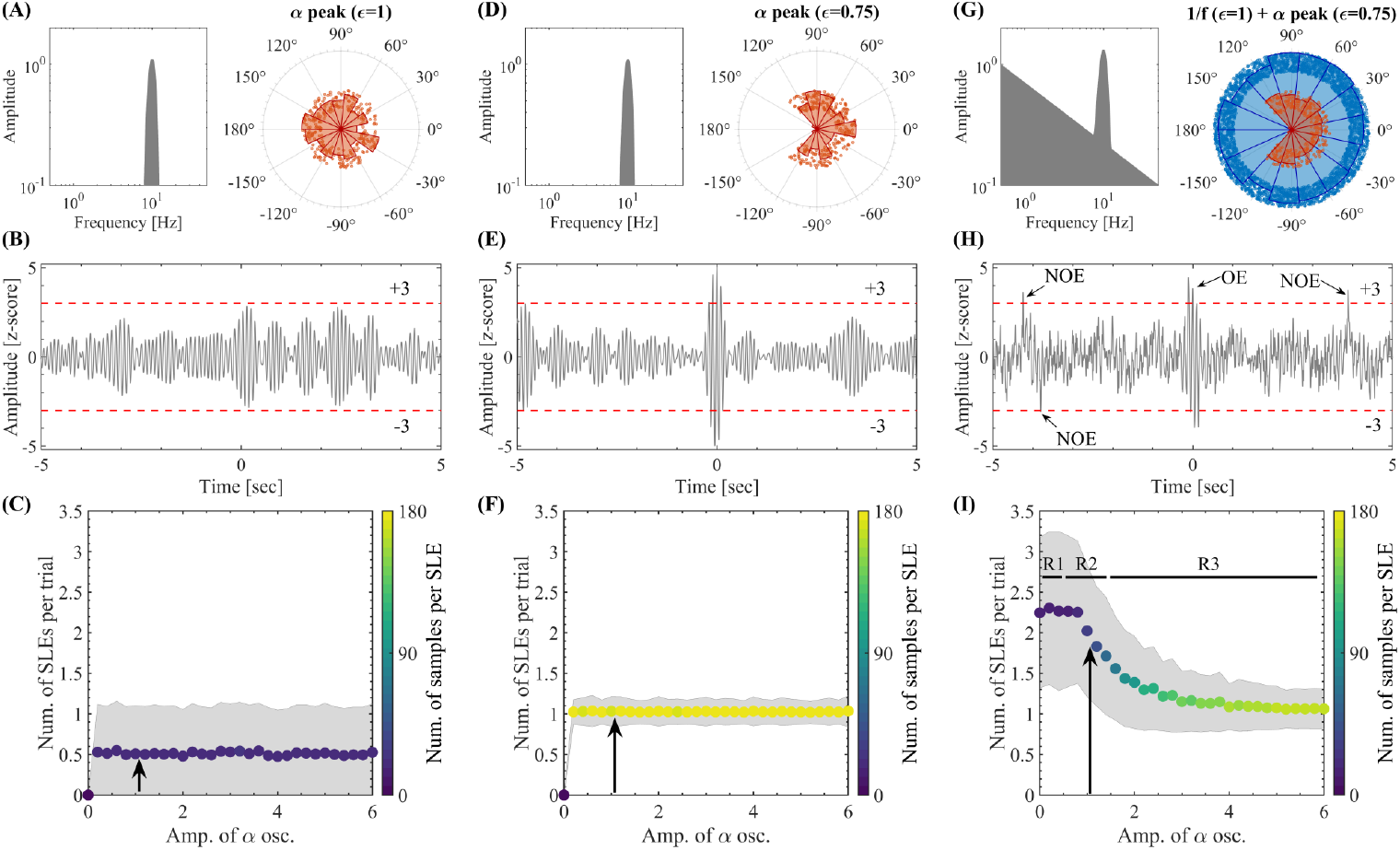
Model for local above-threshold fluctuations. (A) Amplitude spectrum (left) and distribution of the phase values assigned to the spectral components (right) for the oscillatory activity in the alpha band (Hann window with null-to-null bandwidth = 8-13 Hz, frequency resolution *df* = 1*/*60sec ≈ 0.017 Hz). Random phases were assigned to all the spectral components within the range [*− ϵπ, ϵπ*] with a phase factor *ϵ* = 1. (B) 10 sec epoch extracted from the synthetic time series produced by the amplitude spectrum and phase distribution shown in panel A (sampling rate of *f*_*s*_ = 1024 Hz). The horizontal dashed lines in red indicate the 3 standard deviations (*±*3*σ*) thresholds used to compute the SLEs as above-threshold amplitude fluctuations. (C) Number of SLEs per trial as a function of the maximum amplitude of the oscillatory activity in the alpha band. For each maximum amplitude value, we counted the number of SLEs across 1000 time series of 10 sec in duration (trials) synthesized as the one shown in panel B. In each trial, we recomputed the random phases of the spectral components within the range [*− ϵπ, ϵπ*] with *ϵ* = 1. The colored markers indicate the mean number of SLEs per trial across the 1000 trials. The shaded error bars in gray correspond to the standard deviation around the mean value. The pseudocolor scale represents the mean value for the number of above-threshold samples per SLE. The black arrow indicates the maximum amplitude of the alpha oscillations used in panels A and B. (D-F) Same as in A-C for spectral components with random phases constrained within the range [*−ϵπ, ϵπ*] with *ϵ* = 0.75 (see the distribution of the phase values in panel D right). (G) Amplitude spectral profile (left) resulting from the linear superposition of 1) a narrowband amplitude spectrum around the alpha band (Hann window with null-to-null bandwidth = 8-13 Hz), and 2) a set of spectral components with power *A*^2^(*f*) ∝ 1*/f* (frequency resolution *df* = 1*/*60sec ≈ 0.017 Hz). The right side of panel G shows the distribution of phase values assigned to the spectral components. Random phases within the range [*− ϵπ, ϵπ*] with *ϵ* = 1 where assigned to the spectral components constituting the 1*/f* background (blue circles) and *ϵ* = 0.75 where assigned to the spectral components associated with the alpha bump (red circles). (H) Same as in B and E for the spectrum shown in panel G. In this case, it is possible to distinguish Oscillatory (OEs) and Non-Oscillatory (NOEs) Salient Local Events. R1, R2 and R3 indicate regions characterized by *Amp. of alpha oscillations* less than, approx. equal to and greater than the *Amp. of* 1*/f activity*, respectively. Symbols and abbreviations: SLEs, Salient Local Events; OEs, Oscillatory Salient Local Events; NOEs, Non-Oscillatory Salient Local Events.

- *Time-domain representation*: Waveform shape of the *x*(*t*) (inverse DFT, Eq. A.5).
- *Group delay-domain representation*: Amplitude-modulated ocillatory constituents of *x*(*t*) defined by the adjacent pairs *X*(2*k*), *X*(2*k* + 1) in Eq. A.7.
- *Frequency-domain representation*: Constant-amplitude oscillatory constituents of *x*(*t*) defined by the spectral components *X*(*k*) in the DFT (Eq. A.4).

We used the group delay-domain representation to analytically show that the emergence of SEs (i.e., above-threshold fluctuations) in the time-domain, is associated with a high group delay consistency across the oscillatory components in the frequency-domain representation (i.e., approx. constant group delay disclosed by the Fourier constituents of the signal, see Appendix A.2). This mathematical fact, conceptually illustrated in Fig. 4, constitutes an essential signal-level feature inherent to the frequency-domain representation of time series and holds true regardless of both the *x*(*t*) waveform shapes and the underlying biological mechanisms associated with the analyzed SEs.

The group delay is defined as the rate of change of the phase with respect to the frequency, then, a constant group delay across the Fourier frequencies implies a constant incremental phase across frequencies (provided that Δ*ω* = const.). Thus, highly structured Fourier phase values, that is, incremental phase values disclosing low variability across frequencies, promote the time alignment of the amplitude modulated components of the signal (see Fig. 4), and therefore, the emergence of transient above-threshold fluctuations. To quantitatively assess this effect, we introduce the SGDC measures as described below.

The spectral group delay associated with the activity of the brain region *r*, is defined as the rate of change of the phase *ϕ*_*r*_(*ω*) with the frequency *ω* computed on the Fourier spectrum of the brain activity (i.e., the DFT): *τ*_*r*_(*ω*) = Δ*ϕ*_*r*_(*ω*)*/*Δ*ω*(*ω*). The incremental phase Δ*ϕ*_*r*_(*ω*) is defined as the phase difference between spectral components (adjacent in frequency *ω*) constituting the neural activity of the brain region *r*. The theoretical analysis presented in Appendix A.2 shows that the spectral group delay consistency (SGDC) is an important feature linking the oscillatory properties of a signal to the above-threshold fluctuations associated with SEs. For an in-depth mathematical description of the oscillatory mechanisms eliciting above-threshold fluctuations in the brain signals and the measures quantifying the SGDC, the reader is referred to Appendix A.2 and Appendix A.3. Here, we briefly introduce the SGDC measures designed to efficiently quantify this feature in the experimental data,

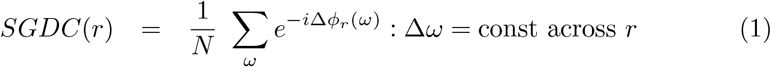

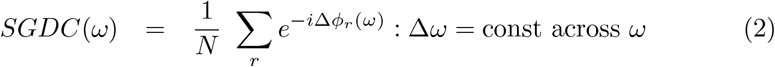

Eqs. 1 and 2 define the SGDC measures as the Euler transformed incremental phase values Δ*ϕ*_*r*_(*ω*) averaged across the spectral components or brain regions, respectively, with *N* being the number of either frequency values or brain regions as appropriate. Importantly, the *SGDC*(*r*) measure (Eq. 1) assesses the emergence of local above-threshold fluctuations from the spectral components constituting the activity of the brain region *r*, whereas the *SGDC*(*ω*) measure (Eq. 2) quantifies the synchronization of the above-threshold fluctuations at the frequency *ω* across brain regions. We also define the pairwise spectral group delay consistency (pSGDC) to quantify the burstiness and cross-regional bursts synchronization in a single measure.

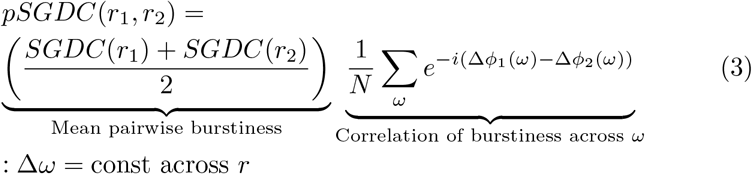

Eq. 3 shows that *pSGDC*(*r*_1_, *r*_2_) is a linear measure conformed by two factors: a factor quantifying the cross-regional correlation between the group delays across the frequency values, weighted by a coefficient quantifying the burstiness of the two involved brain regions (*r*_1_, *r*_2_).

The SGDC measures (Eqs. 1, 2 and 3) were computed using both non-time-resolved and time-resolved approaches. In the non-time-resolved case, the SGDC measures (Eqs. 1, 2 and 3) were computed on the whole time series of each brain region. That is, we first obtain the phase values corresponding to the Fourier spectral components by computing the DFT (via the Fast Fourier Transform algorithm) on the whole time series of each brain region. Then, SGDC measures (Eqs. 1, 2 and 3) were computed on the incremental phase Δ*ϕ*_*r*_(*ω*) obtained as the phase difference between the Fourier spectral components taken in non-overlapping adjacent pairs across the frequency *ω*. In particular, this non-time-resolved approach was used to produce the results shown in the Figs. 7 and A.7A,B. On the other hand, in Figs. C.11 and A.7C,D we follow a time-resolved approach. That is, the *SGDC*(*r*) and *SGDC*(*ω*) measures (Eqs. 1 and 2) were computed on each detected SE by considering the brain regions and time interval associated with each particular event. In the case of the Fig. C.11, the *SGDC*(*r*) and *SGDC*(*ω*) arrays were averaged selectively across the SEs segregated in the two clusters produced by the Louvain algorithm (see Section 2.9).

## 3. RESULTS

### 3.1. Statistical, spatiotemporal, and spectral characterization of salient events

We identified SEs in our dataset and studied their characteristic signatures. In particular, we introduced a comprehensive array of features describing the statistical, spatiotemporal and spectral properties of SEs. The proposed tools allowed for the characterization of SNEs by the way they spread across the brain network. Indeed, we found the role that each brain region plays in the propagation of these SNEs is not homogeneous. To characterize the spatiotemporal profiles of SEs, we defined two ROI-wise metrics (see Methods, Section 2.4): The *mean event duration* measures the typical duration of SEs propagating through a brain region (Figs. 1A,B); and the *mean event size* measures the typical size of SNEs propagating through a brain region (Figs. 1C,D). The brain plots in Figs. 1B,D reveal a characteristic topography, demonstrating the heterogeneous role that each brain region plays in the propagation of SNEs. In particular, SEs with bigger size and longer duration seems to be more associated with the temporal and deep brain regions.

Regarding the statistical characterization, we found that the SEs detected in our MEG data obtained from 47 subjects (1 min MEG time series source-reconstructed to 84 brain regions), disclose exponential-like distributions of the events size and duration with steep slope exponents (⪅ *−*3, see Fig. C.3), which do not follow the power law statistics putatively associated with the dynamical regime around a critical point or phase transition (see Introduction). Importantly, the exponential-like distributions of the events size and duration shown in the Fig. C.3 do not modify significantly when the time binning value is varied from 1 to 5 samples per time bin (time binning ranging from 1 ms to 5 ms, data not shown).

Next, we introduce a tool to characterize the spectral signature of SEs, by first transforming the regional signals into a time-frequency representation and then averaging the time-frequency maps selectively across the time points corresponding to the occurrence of each SE (Figs. 2A-D). This way, we defined the spectral fingerprint of each SE, which we named Event Spectral Matrix (ESM, see Methods and Figs. 2E-G). Of note, the ESM does not represent the frequency content of SEs since the latter are very short-duration transient events, instead, the ESM reveals the spectral signature associated with the oscillatory activity co-occurring with each SE. That is, the ESM reveals the co-occurrence (or coupling) between the oscillatory activity and SEs across brain regions. Fig. 2 displays the ESM for a single event (panel E), the average ESM across 10 subjects (panel F) and the ESM averaged across all the 47 subjects (panel G). Figs. 2F,G show that the oscillatory activity of most brain regions peaks in the alpha band (8-13 Hz) during SEs. In other words, during SEs, brain regions fluctuate predominantly in alpha. This is also observed away from the occipital regions, suggesting that synchronization in the alpha band might spread on a large-scale during SNEs. Note that this result provides a relevant insight regarding the connection between SEs and NOs and, it is non-trivial since SEs are rare phenomena, occupying only a small fraction of the total recording (in space and time).

### 3.2. Salient events and phase coherence: surrogate data analysis

The spectral signature in the alpha band disclosed by the averaged ESMs shown in Figs. 2F,G suggest that a significant fraction of the SEs observed in our MEG data co-occur with (or are coupled to) alpha oscillations. To test this hypothesis, we generated 100 C-surrogate sets of SEs (see Methods, Section 2.8) that randomize the starting time of each observed SE and keep unaltered all the other properties like the time width and brain regions recruited during each event. Importantly, as shown in the Fig. 2F, the average ESM of the true SEs thresholded with the average ESM of the C-surrogate SEs (see Methods, Section 2.5) discloses a prominent spectral signature in the alpha band. This result reveals a significant (i.e., above chance level) coupling between the true SEs and alpha oscillations, supporting our hypothesis that the large-scale spreading of transient alpha bursts is associated with SNEs. Taking together these results suggest that during SNEs, the brain activity display large deviations from the baseline, which are coordinated across regions, giving rise to complex activation patterns with well-defined statistical, spatiotemporal, and spectral features.

To investigate the statistical properties of the signals associated with the emergence of realistic SEs, we first tested whether the observed SEs require additional features beyond the autocorrelation (PSD) of each MEG trace, which could include cross-correlation, non-stationarity, or non-Gaussianity. A common way to test the necessary and/or sufficient conditions underlying a phenomenon (here, SEs) is the use of surrogate data analysis [51]. This approach involves creating surrogate datasets that remove or alter a specific property (e.g., phase relationships) while preserving other statistical characteristics, allowing one to determine if the absence or modification of the property affects the observed feature of interest. Following this line of reasoning, we generated 100 phase-randomized A-surrogate datasets (see Methods). Each A-surrogate preserves the PSD (and thus the autocorrelation) of each brain region but disrupts the phase relationships of spectral components. When phases are randomized independently across regions, this procedure also disrupts inter-regional phase relationships and therefore removes cross-correlation structure that depends on those phases. Hence, A-surrogates implement the null hypothesis that the observed SEs can be explained solely by the preserved PSDs (i.e., by stationary, approximately Gaussian signals with inter-regional phase relationships removed). Despite the A-surrogates having the same spectral content as the original data, they disclose distributions with significantly less SEs with large size and duration values when compared to those observed in the true data (see A-surrogates in Figs. C.3A,B). Besides, A-surrogates do not reproduce realistic spatiotemporal patterns of propagation (see A-surrogates in Figs. 1A,C) and ESMs (data not shown).

We then tested whether the observed SEs require additional structure beyond the auto- and cross-correlation of the MEG trace, which could include non-stationarity, or non-Gaussianity. To test this hypothesis, we generated 100 phase-randomized B-surrogate datasets (see Methods) that randomize the phases similarly to the A-surrogate, but in this case preserving both the regional PSDs and the cross-spectra. The preservation of cross-spectra implies that the phase difference between any pair of brain regions in *homologous frequency components* is preserved. This implies to preserve Pearson’s correlations between brain regions (see Appendix A.1 and Fig. A.1). However, the B-surrogates destroy the phase relationships between *non-homologous frequency components*.

B-surrogates therefore implement the null hypothesis that the observed SEs can be explained by the preserved auto- and cross-correlation (i.e., by stationary, approximately Gaussian signals with inter-regional phase relationships preserved). The observed mean spatiotemporal properties (see B-surrogates in Figs. 1A,C), the alpha signature disclosed by the ESM (see the average ESM thresholded using the B-surrogates shown in Fig. 2G), and the distributions of SEs duration and size (see B-surrogates in Figs. C.3A,B) are not explained by the B-surrogates. Notice that these results are non-trivial, since in both the original and the B-surrogate datasets the number of SEs is almost identical, and large events are also observed in the surrogate data (see B-surrogates in Figs. C.3A,B).

To summarize, despite retaining the same power spectra and cross-spectra, the loss of synchronization across spectral components (given by the phase randomization), impairs large-scale coordinated SNEs, significantly disrupting the statistics and features of SEs.

### 3.3. Clustering of salient events

The ESM can be defined at the single event level (Fig. 2E). Thus, we asked if SEs with different spectral signatures propagate differently. In particular, we hypothesized a relationship between the event spectral signature (as measured by the ESM) and the event duration, size and propagation topographies (see Methods). To test this relationship, we clustered SEs according to their ESM using the Louvain algorithm (see Methods). We found that SEs cluster into two main groups based on their spectral signature (Figs. 3A,B). The SEs belonging to cluster 1 (Fig. 3A) display less marked and widespread alpha peak in the ESM as compared to cluster 2 (Fig. 3B). Importantly, we found a statistically significant differences in the mean event duration and size between cluster 1 and cluster 2 (see Figs. 3E,H). To assess this, in each brain region we computed a non-parametric permutation test (random sampling without replacement, 1 *×* 10^4^ permutations). All the brain regions disclosed a statistically significant difference of the mean event duration and size between cluster 1 and 2 (the Bonferroni-adjusted two-tailed P values result *P <* 0.001 in all the brain regions). Consistently, the two clusters are also well distinguished by their different waveform shapes, with cluster 1 showing shorter temporal profiles of above-threshold fluctuations. Figs. 3C,D show the average waveform shapes of SEs, obtained by averaging in each brain region (BR) the absolute value of the time series associated with each event (see Methods).

These results suggest that cluster 2 events are specifically related to the long-range spread of narrowband alpha bursts across the brain network (i.e., SNEs), whereas cluster 1 events correspond to more short-lived and spatially localized fluctuations mainly promoted by the BAA (i.e., SLEs. See Figs. 3A-D,E,H). Consistently, the two identified clusters are also characterized by different event duration and size, which supports our hypothesis. In particular, cluster 1 events are generally small and short-lived when compared to cluster 2 events, although both clusters display event size and duration distributions spanning across a few orders of magnitude (see Figs. 3G,J). Interestingly, the event duration and size distributions are different between the two clusters, which could have implications for the study of the spectral background statistics.

We also found that SEs propagate in a cluster-specific manner (see Figs. C.5 and C.6 in Appendix C). In cluster 1, the spatial profiles associated with the events start, maximum recruitment and end are highly correlated (see Fig. C.5A, pairwise Pearson’s correlations *r >* 0.978, *P <* 0.001 two-tailed Student’s t-test), pointing out that cluster 1 events do not propagate to brain regions distant from those igniting the events. This result strongly supports the evidence presented above regarding the spatially localized nature of cluster 1 events. On the other hand, the spatial profiles associated with the events start and end are also highly correlated in cluster 2 SEs (see Fig. C.5B, Pearson’s correlation *r* = 0.895, *P <* 0.001 two-tailed Student’s t-test), suggesting that the brain regions involved in the ignition of a particular cluster 2 event tend to remain active until the event extinction. However, the maximum recruitment profile of cluster 2 events disclose a weak negative correlation with respect to the start spatial profile (Pearson’s correlation *r* = −0.298, *P <* 0.01 two-tailed Student’s t-test), supporting our hypothesis that cluster 2 events spread in the form of narrowband alpha bursts across the brain network. Intriguingly, the spatial profiles associated with the events start and end are highly correlated between the two clusters (see Figs. C.6A-C and Figs. C.6G-I), whereas a different scenario is observed in terms of how the brain regions are recruited by the two event clusters. Specifically, brain regions that are recruited by the longer events of cluster 1, will be recruited by the shorter events of cluster 2, and vice versa (see Fig. C.6D). Within cluster 1, the longest SEs occupy the frontal and occipital regions (see Fig. C.6E), whereas in cluster 2, associated with the spectral signature in the alpha band, the longest SEs are in the parietal and temporal regions (see Fig. C.6F). The opposite trend is observed for the shortest SEs. In fact, performing Pearson’s correlations between the spatial profiles of cluster 1 and cluster 2 corresponding to the maximal size of recruitment across brain regions, we obtain a strong negative correlation (*r* = *−*0.841, *P <* 0.001 two-tailed Student’s t-test, see Fig. C.6D). Note that the specificity of cluster 2 events, associated with transient above-threshold alpha bursts, in recruiting parietal and temporal brain regions can not be trivially explained by the presence of elevated (steady) alpha oscillatory power, which is commonly observed in occipital brain regions during the eyes-closed resting state (see Figs. 2B and C.4).

In summary, in this section we have introduced a comprehensive array of SE features, showing that rare, short-lived SEs propagating across the brain during spontaneous resting state activity are highly structured in terms of their spatial, temporal, and spectral properties. In particular, the spectral characterization using the ESM provided relevant insights regarding the connection between the observed SNEs and NOs in the alpha band.

### 3.4. Analytical framework: Spectral group delay consistency

We next explored the mechanism mediating the reduction of local and cross-regional burstiness observed in our surrogate data computed via phase randomization (see Section 3.2). Notice that this is a relevant question since surrogate data analysis based on phase randomization is extensively used in many (bio)physical domains including Neuroscience. Importantly, being the phase an intrinsic property of NOs, it is not obvious how the modification (e.g., randomization) of this oscillation-domain parameter affects the emergence of SEs (compare the true data with the A- and B-surrogates in Figs. 1 and C.3). This question becomes apparent by taking into account that despite preserving both the power spectrum (PSD) in each brain region and the cross-spectra (i.e., functional connectivity) B-surrogates fail to account for the SEs observed in our MEG dataset. To address this question, we developed a signal-level analytical framework, named Spectral Group Delay Consistency (SGDC), designed to provide an analytical rationale supporting the emergence of SEs from the oscillatory constituents of the brain activity.

Let us focus on a single brain activity time series. We first compute the DFT to decompose the time series as a linear superposition of its Fourier oscillatory components (see Figs. 4A,B,D and Eq. A.4). Then, we group the Fourier components in (non-overlapping) pairs adjacent in frequency (see color-paired Fourier components in Fig. 4A). This lead to the pairwise complex baseband representation of the time series. In this representation, the time series of interest is decomposed as a linear superposition of amplitude modulated components (see the color coded amplitude modulated signals in Fig. 4E and Eqs. A.7 and A.17). Importantly, the time offset of each amplitude modulated component is determined by the spectral group delay *τ* (*ω*) ≈ *−*Δ*ϕ*(*ω*)*/*Δ*ω*. Where *τ* (*ω*) is computed on Fourier spectrum of the brain activity (i.e., the DFT), as the rate of change of the phase *ϕ*(*ω*) with the frequency *ω*. Essentially, when all the Fourier components are added together to synthesize the signal in the time-domain (i.e., the inverse DFT), the spectral group delay determines the time alignment of the envelope of the amplitude modulated components associated with each pair of adjacent spectral components. Such time alignment promotes transient large-amplitude excursions of the signal (i.e., above-threshold fluctuations). In the case of adjacent spectral components with phase values depending linearly with *ω*, we obtain approximately constant spectral group delay values for all the pairs of adjacent spectral components (see Fig. 4C). In such a case, the signal has a high spectral group delay consistency (SGDC) which promotes the time alignment of the amplitude modulated components (see the color coded amplitude modulated signals in Fig. 4E), hence, supporting the occurrence of above-threshold fluctuations (see the large-amplitude excursions of the black time series in Figs. 4D,E). On the other hand, for adjacent frequency components having phase values disclosing a nonlinear dependence with the frequency *ω* (e.g., a quadratic dependence as shown in Fig. A.2G), the resulting spectral group delay depends on *ω* (see Figs. A.2H). The latter disrupts the time alignment of the amplitude modulated components (see the color coded amplitude modulated signals in Fig. A.2J). In this case, we say that the signal has low SGDC which reduces the occurrence of above-threshold fluctuations (see the sub-threshold fluctuations of the black time series in Figs. A.2I,J).

The results discussed above constitutes strong analytical arguments pointing out that the reduction of the local burstiness observed in our surrogate data computed via phase randomization, can be understood in terms of the group delay consistency across the spectral components of the neuronal activity (i.e., SGDC). Specifically, the phase randomization process produces phase values having a nonlinear (random) dependence with the frequency of the Fourier components, hence, reducing the SGDC of the resulting surrogate time series. We confirmed this theoretical results using analytically tractable model of synthetic time series (see the discussion about Figs. A.2 and A.3 in Appendix A.2), numerical simulations (see Section 3.5) and empirical MEG data (see the discussion about Figs. A.7A,C in Appendix A.4). In particular, in Appendix A.4 we analytically show that, despite preserving the regional power spectrum (PSD), the phase randomization associated with both A- and B-surrogates significantly reduces the burstiness of each brain region as assessed by the *SGDC*(*r*) measure (Eq. 1). Importantly, the reduction of the regional SGDC, as quantified by the *SGDC*(*r*) measure, offers an analytical rationale supporting the evidence showing that B-surrogates failed to reproduce the SEs observed in our MEG dataset despite preserving both the regional PSDs and the cross-spectra. As a conclusion, the SGDC constitutes a signal-level analytical model linking the emergence of SEs from the oscillatory components of the brain activity and underpining the evidence showing that our A- and B-surrogates computed via phase randomization failed to reproduce realistic SEs (see Section 3.2).

For an in-depth mathematical description of the SGDC framework and measures, the reader is referred to Appendix A.2.

### 3.5. Numerical models: SEs, NOs and BAA

We built a numerical signal model to elucidate the relation between SEs, NOs, and BAA. We model the activity of single brain regions as the linear superposition of Fourier components oscillating in a narrow frequency band. As a result, the corresponding spectral representation discloses a “bump” of (null-to-null) bandwidth in the alpha band (8-13 Hz, Figs. 5A,D,G). The BAA was modeled by imposing a 1*/f* trend in the PSD of each signal (Fig. 5G). This 1*/f* spectral background was chosen to mimic the 10*dB/dec* log-log decay rate observed in the PSDs associated with our MEG dataset (see Fig. C.4). To model different degrees of phase coherence, we assign random phase values to the spectral components within a range [*− ϵπ, ϵπ*] with *ϵ* ∈ [0, 1] (see the polar plots in Figs. 5A,D,G). On the one hand, for *ϵ* ≃ 1, the spectral components of the signal were desynchronized (i.e., independent oscillatory components, Fig. 5A). On the other hand, for *ϵ* ≃0 the spectral components were highly synchronized (i.e., high cross-frequency coherence). We first focused on a *single brain signal* and measured the number of SLEs (i.e., transient amplitude excursions above a fixed threshold of 3 standard deviations: *±*3*σ*) across 1000 realizations (i.e., trials), depending on the presence or absence of coherent NOs and 1*/f* activity (see Figs. 5C,F,I). In the absence of 1*/f* activity and for uniformly distributed random phases assigned to the spectral components in the alpha band (*ϵ* = 1, Figs. 5A), the model displays very few above-threshold fluctuations across trials (Figs. 5B,C). Increasing the coherence of the spectral components in the alpha band (*ϵ* = 0.75, Figs. 5D), despite the absence of 1*/f* activity, the number of above-threshold fluctuations increased, producing a salient burst in most of the trials (Figs. 5E,F). Importantly, Fig. B.1 shows that the results discussed above, in connection with the emergence of local above-threshold fluctuations from the Fourier oscillatory constituents of the brain activity (i.e., SLEs), can be understood in terms of the SGDC as quantified by the *SGDC*(*r*) measure (Eq. 1). Specifically, Fig. B.1 shows that the increase of the signal burstiness, as quantified by the kurtosis of the signal’s amplitude values, associated with more constrained random phase values (i.e., low phase factor *ϵ* values) correlates with the increase in the SGDC as quantified by the *SGDC*(*r*) measure. In Fig. B.1, the time series were synthesized by adding pure sinusoidal signals. The *SGDC*(*r*) was then computed directly from the synthetic phases of these sinusoidal components. Because the phases were taken from the exact analytical components, no spectral leakage was present in this case. In contrast, in Fig. B.2, 1 min in duration time series were synthesized following the same procedure as in Fig. B.1, but this time the *SGDC*(*r*) was computed using the alpha band phases obtained from the DFT applied to the synthesized time series. This procedure inherently introduces spectral leakage due to the time-domain tapering (rectangular window), which affects the phase values involved in the computation of the *SGDC*(*r*) measure and is visible in the corresponding power spectra. Fig. B.2 shows that the increase of the salience of transient fluctuations in a signal, as quantified by the kurtosis of the signal’s amplitude values, is reproduced by the *SGDC*(*r*) measure. Importantly, these results highlight that the *SGDC*(*r*) measure is not primarily driven by the spectral leakage. Instead, it reflects the relationship between the salience of transient fluctuations and the consistency (spread or variability) of the group delay across the Fourier frequencies, independently of the spectral leakage. In addition, we re-compute the signal model for the same set of phase factor values used in Fig. B.1, this time using spectral phase values disclosing not a random but a linear dependence with the frequency (i.e., a time-shift in the time-domain). The results obtained with this configuration are shown in Fig. B.3. As predicted by the SGDC mechanism (see Figs. 4A-E), we obtained *SGDC*(*r*) 1 independently of the phase factor value (*ϵ* ∈ [0, 1]), and the time series produced by the signal model disclosed (time-shifted) above-threshold fluctuations in all the cases (see Fig. B.3). These numerical results constitute further evidence showing that the SGDC effectively underlies the emergence of local above-threshold fluctuations from NOs, as in the case shown in Figs. 5D,E,F. Then, we introduced the broadband 1*/f* activity into the model through a linear superposition (addition) with the oscillatory activity in the alpha band. As a result, the presence of the broadband 1*/f* activity with *ϵ* = 1 and coherent spectral components in the alpha band with *ϵ* = 0.75 (Fig. 5G) further increased the number of salient events in a single brain signal (Figs. 5H,I). Importantly, the 1*/f* activity also influences the rhythmicity of above-threshold fluctuations, which occur aperiodically. More specifically, if we synthesize a long time series by concatenating trials constructed without the 1*/f* activity (as in Figs. 5E), the concatenated time series will disclose a periodic series of above-threshold alpha bursts (i.e., one salient alpha burst per trial). Instead, in the presence of 1*/f* activity we obtain above-threshold fluctuations occurring aperiodically in each trial besides the salient alpha burst, hence, the time series resulting from concatenating trials (as in Fig. 5H) will disclose an aperiodic series of above-threshold fluctuations, elicited by the interaction between the 1*/f* and oscillatory activities. Furthermore, the regime R2 in Fig. 5I points out a plausible range for the relative amplitude between NOs and the BAA in order to obtain realistic aperiodically occurring above-threshold fluctuations. That is, in the regime R1 only Non-Oscillatory Salient Events (NOEs) are observed, in the regime R3 only Oscillatory Salient Events (OEs) are observed. In contrast, the regime R2 is characterized by a stochastic-resonance-like effect in which the resulting local activity exhibits both NOEs and OEs mirroring the two SE clusters observed in our MEG dataset. In Appendix C.2 we discuss additional empirical evidence supporting the theoretical findings described in Sections 3.4 and 3.5.

In summary, these results suggest that the mere presence of oscillations associated with an increase of power around a narrow frequency band does not guarantee the stable occurrence of above-threshold fluctuations (Figs. 5A-C). However, if the phases of the spectral components are coherent producing high |*SGDC*(*r*)| values, then high-amplitude fluctuations are consistently observed in the signal (Figs. 5D-F).

Next, we extended the above setup to model *whole-brain activity* and SNEs. For each simulated brain signal, we set the amplitude of the alpha peak (with alpha amplitude ∈ [0, 1]) proportionally to the mean alpha amplitude (average across the 47 participants) observed in the empirical MEG recordings, thus modeling the non-homogeneous presence of alpha activity across brain regions. In addition, in each region, we bounded the random phases assigned to the spectral components in the alpha band within a range [*− ϵπ, ϵπ*], whose width *ϵ* ∈ [0.75, 1] was inversely proportional to the empirical alpha power (i.e., the higher the alpha peak, the higher the phase coherence among the spectral components). This choice was motivated by the fact that high PSD bumps are generally interpreted as stronger narrowband synchronization within local neuronal populations [22] (see Discussion). Using this setup, we measured synthetic SEs and tested their dependence on the 1*/f* activity. When only alpha oscillations were present, and no broadband 1*/f* activity (Fig. 6A), the resulting ESM was not realistic compared to the empiric one (compare Figs. 6B and 2G), and the distributions of SEs duration and size were not approximating the exponential-like distributions observed in our MEG dataset (compare Figs. B.4A,B and C.3). Instead, when only broadband 1*/f* activity was present, and no oscillatory activity in the alpha band nor coherent phase values were used (i.e., *ϵ* = 1; Fig. 6C), the ESM did not show the spectral signature associated with the alpha component (Fig. 6D). Also, the distribution of SEs duration was similar to the empirical data, while the distribution of SEs size was shrunk, as the model did not display SEs involving large populations (Figs. B.4C,D). Finally, when both broadband 1*/f* activity and alpha oscillations were simultaneously present (Fig. 6E), the emerging SEs displayed a realistic ESM (compare Figs. 6F and 2G) as well as exponential-like distributions of SEs duration and size (Figs. B.4E,F); although the SEs size decayed in a markedly more rapid fashion than in the empirical data (compare Figs. B.4E,F and C.3). The Pearson’s correlation between the vectorized versions of the empirical (Fig. 2G) and simulated (Figs. 6B,D,F) ESMs are as follows: Empirical (non-thresholded version of the ESM shown in Fig. 2G) vs. Large scale model including only alpha oscillations (ESM shown in Fig. 6A): *r* = 0.594, *P <* 0.001. Empirical (non-thresholded version of the ESM shown in Fig. 2G) vs. Large scale model including only broadband arrhythmic activity (ESM shown in Fig. 6D): *r* = 0.167, *P <* 0.001. Empirical (non-thresholded version of the ESM shown in Fig. 2G) vs. Large scale model including both alpha oscillations and broadband arrhythmic activity (ESM shown in Fig. 6F): *r* = 0.611, *P <* 0.001. The statistical significance of these linear correlations was assessed by using the Student’s t distributions of the two-tailed hypothesis test under the null hypothesis that the correlation is zero.

**Figure 6:**
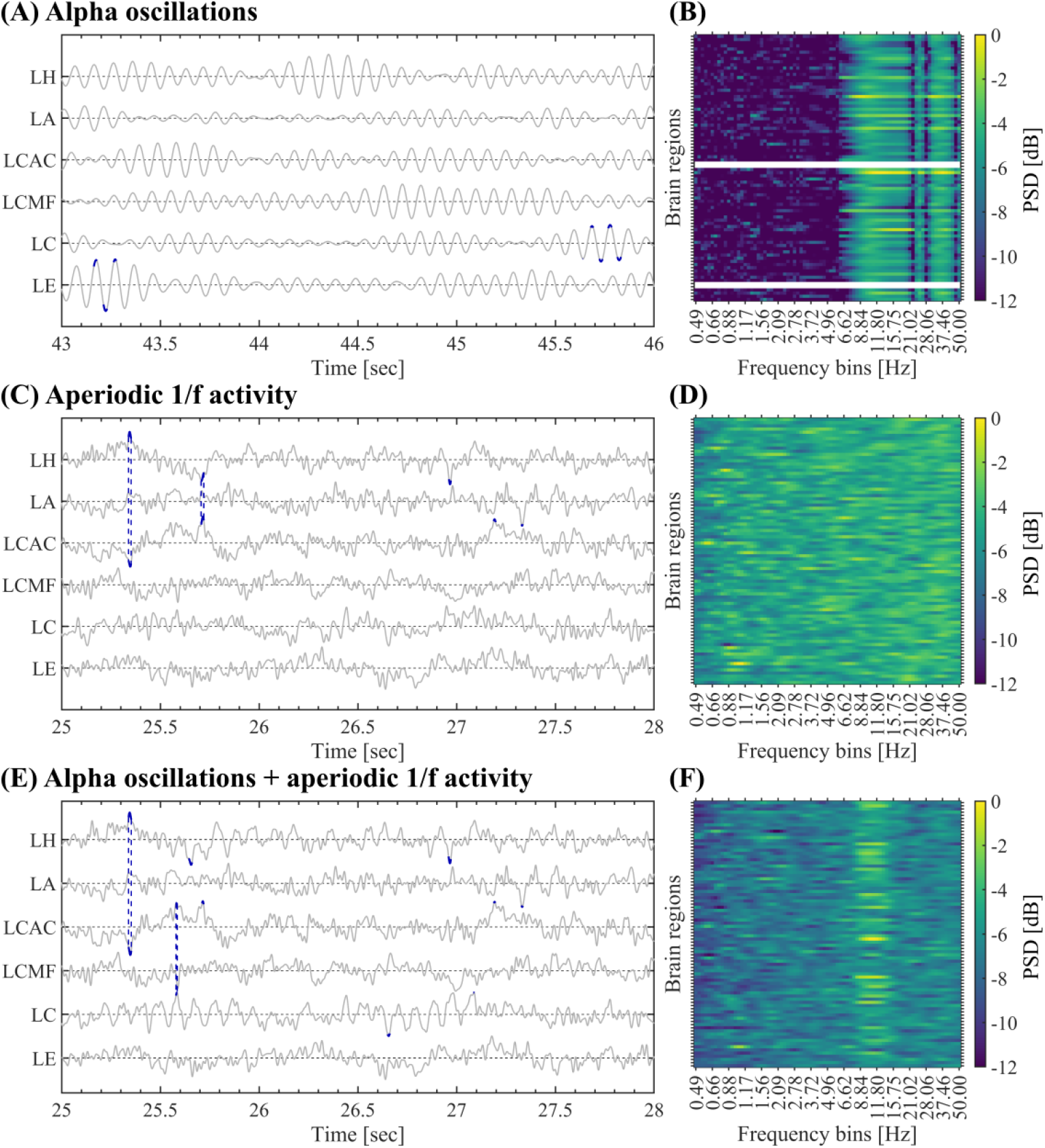
Large-scale signal model for SEs. (A-B) Large-scale model for SEs including only alpha oscillations (random phase values in the alpha band constrained to the range [*ϵπ, ϵπ*] with *ϵ* ∈ [0.75, 1]). Panel A shows a subset of synthetic activities. In each time series, the above-threshold fluctuations (*±*3*σ*) are highlighted in dark blue. Vertical dashed lines connect the activations associated with SEs completely contained in the subset of signals shown. Panel B shows the resulting ESM averaged across the SEs. Panels A-B were computed on all the SEs detected in a simulated time series of 1-minute duration. (C-D) Same as in A-B for the large-scale model including only broadband 1*/f* activity, and no oscillatory activity in the alpha band nor phase consistency values were present (*ϵ* = 1). (E-F) Same as in A-B for the large-scale model including both broadband 1*/f* activity with non-constrained random phases (*ϵ* = 1) and alpha oscillations with random phases constrained proportionally to the observed alpha power in the range (*ϵ* ∈ [0.75, 1]). Symbols and abbreviations: SEs, Salient Events; LH, Left Hippocampus; LA, Left Amygdala; LCAC, Left Caudal Anterior Cingulate; LCMF, Left Caudal Middle Frontal; LC, Left Cuneus; LE, Left Entorhinal.

**Figure 7:**
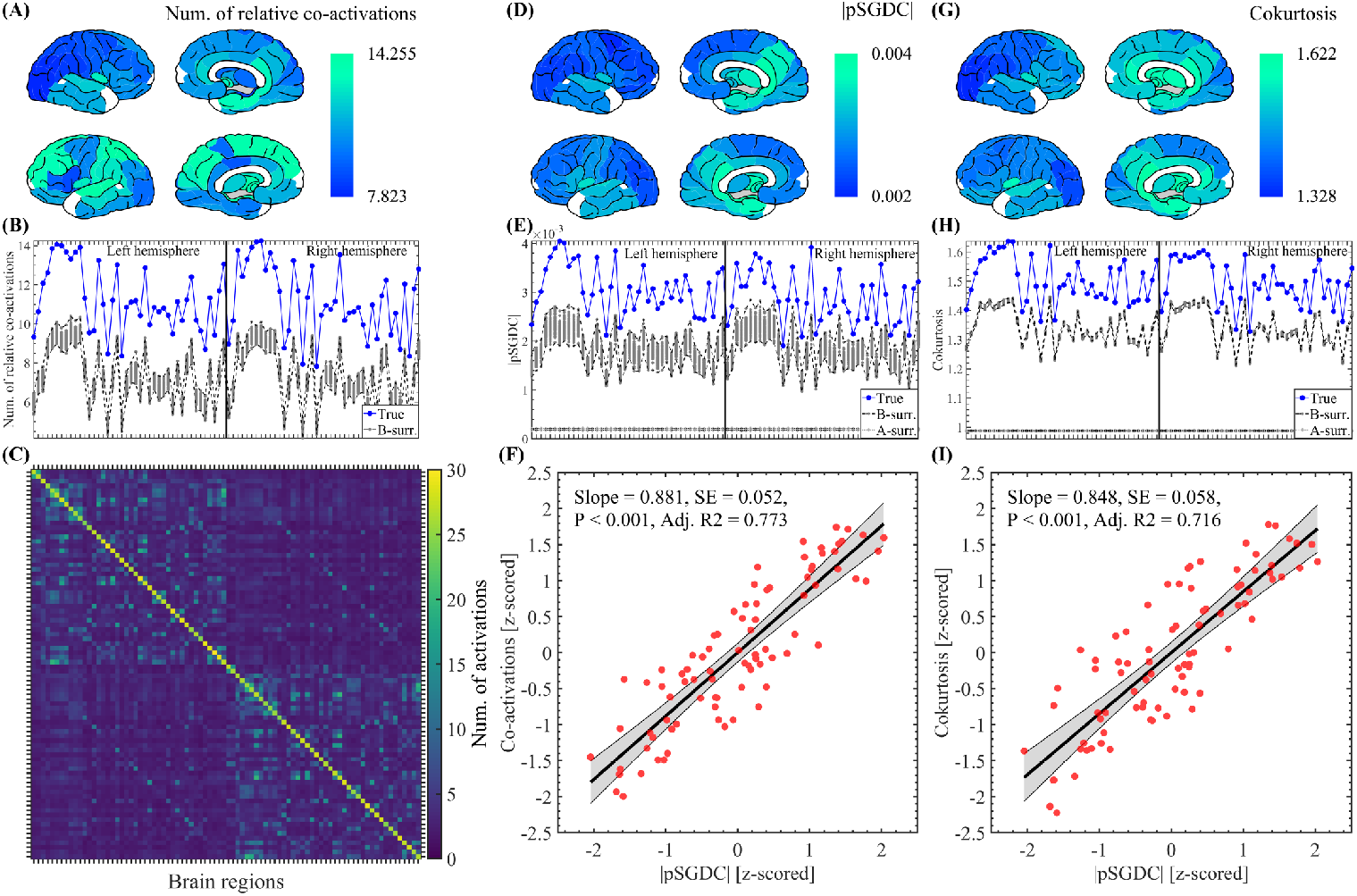
Co-activation pattern of SEs compared against pSGDC and cokurstosis measures computed on whole time series of the brain regions taken in pairs. (A) Brain topographies corresponding to the co-activation profile shown in panel B (blue markers). (B) Spatial profile showing the number of relative co-activations (mean value across the 47 participants), i.e., the accumulated number of activations in each row of the co-activation matrix relative to the total number of activations in each brain region (diagonal of the co-activation matrix). Note that the spatial profiles corresponding to the 100 B-surrogates (dark gray markers) fail to reproduce the spatial profile associated with the true MEG data (blue markers). (C) Co-activation matrix averaged across the SEs observed in the 47 participants (see Section 2.3 in Methods). (D-E) Same as in A-B for the pSGDC measure. (F) Scatter plot showing the correlation between the co-activation and pSGDC spatial profiles shown in panels B and E, respectively. Number of samples (red circles) = Number of brain regions = 84. The thick black line and black shaded error bars represent the linear regression and the 95% confidence interval, respectively. The reported P value for the statistical significance of the linear regression was assessed using Student’s t distributions of the two-tailed hypothesis test under the null hypothesis that the correlation is zero. (G-H) Same as in D-E for the cokurtosis measure. (I) Same as in F for the correlation between the cokurtosis and pSGDC spatial profiles. In panels B, E and H, the labels and ordering of the brain regions are the same as those shown in Fig. C.2. Symbols and abbreviations: SEs, Salient Events; pSGDC, pairwise Spectral Group Delay Consistency.

These results suggest that both NOs and broadband 1*/f* spectral background contribute to the signal deviations from baseline activity and realistic SEs, provided that the narrowband spectral components display appropriate levels of SGDC.

### 3.6. Mechanisms of long-range interactions

Whereas the *SGDC*(*r*) assesses the emergence of local above-threshold fluctuations from the Fourier oscillatory constituents of the activity in a single brain region (i.e., SLEs), it does not account for cross-regional effects associated with SNEs. To quantitatively study the cross-regional effects of SGDC on our data we introduce the *SGDC*(*ω*) measure (Eq. 2). The magnitude of *SGDC*(*ω*) is bounded in the range [0, 1] and quantifies how much the group delay at a given frequency *ω* varies across brain regions. By using synthetic time series, in Appendix A.3 we show that the *SGDC*(*ω*) measure assesses the contribution of each frequency component in the co-activation (synchronization in time) of above-threshold fluctuations across brain regions (see Figs. A.5 and A.6). Of note, Figs. A.5 and A.6 show that the *SGDC*(*ω*) measure effectively resolves the cross-regional synchronization of SEs across frequency bands, whereas phase coherence measures (e.g., PLV: Phase Locking Value) are completely blind to this effect (see detailed description in Appendix A.3). In Appendix C.2 we present additional empirical evidence supporting the connection between the *SGDC*(*r*) and *SGDC*(*ω*) measures and the emergence of local and large-scale salient events. In particular, Fig. C.11C shows that only cluster 2 SEs, associated with the spectral signature in the alpha band, disclose |*SGDC*(*r*)| values higher than those disclosed by the C-surrogate SEs. Importantly, Fig. C.11D shows the increase of transient cross-regional coherence around the alpha band, as quantified by the *SGDC*(*ω*) measure, associated with the SEs disclosing the alpha spectral signature in the average ESM (i.e., cluster 2 SEs). Notably, Fig. C.11E shows that the transient cross-regional coherence around the alpha band associated with the cluster 2 SEs is also captured by the large-scale model presented in Section 3.5.

Synchrony is thought to play a role in coordinating information processing across different brain regions. However, correlation structures such as hemodynamic functional connectivity are better explained in terms of power amplitude correlations of electrophysiological signals (e.g., MEG), rather than phase-synchrony. In a recent work, it was demonstrated that power correlation between two signals can be analytically decomposed into signal coherence (a measure of phase synchronization), cokurtosis (a measure of the probability of simultaneous large fluctuations), and conjugate-coherence [28]. In particular, it was proposed that the cokurtosis between two signals provides a measure of co-bursting that offers a robust neurophysiological correlate for hemodynamic resting-state networks [28]. Here we show that the SGDC conceptualization provides a coherent account of both the co-burstiness and the cokurtosis in terms of the group delay consistency of the signals’ spectral content, therefore, advancing our understanding of the signal-level mechanisms of long-range communication. For this, we counted the co-participation of pairs of brain regions across SEs (see Methods, Section 2.3). Fig. 7C shows the co-activation matrix indicating the number of co-activations between each pair of brain regions. Fig. 7B shows the number of relative co-activations, i.e., the accumulated number of activations in each row of the co-activation matrix relative to the total number of activations in each brain region (diagonal of the co-activation matrix). Fig. 7A displays the brain plots corresponding to the number of relative co-activations shown in Fig. 7B. Importantly, the topography of co-activations shown in the Figs. 7A-C can not be trivially explained by the chance co-occurrence of rare above-threshold fluctuations in the brain activity. Note that the B-surrogates shown in Fig. 7B fail to reproduce the the topography of co-activations despite preserving both the power spectrum (PSD) in each brain region and the cross-correlations (i.e., functional connectivity). Moreover, we found that the the kurtosis and *SGDC*(*r*), two measures related to the occurrence of local above-threshold fluctuations (i.e., SLEs), when computed in a non-time-resolved manner in each brain region fail to reproduce the topography of co-activations associated with the observed SEs. In the case of the *SGDC*(*r*) measure, compare the spatial profiles shown in Figs. A.7A and 7B. To account for both the burstiness and cross-regional bursts synchronization in a non-time-resolved manner we used the pairwise SGDC measure (pSGDC). The *pSGDC*(*r*_1_, *r*_2_) is defined as the product of two fators: a factor quantifying the cross-regional correlation between the group delays across the frequency components, weighted by the average *SGDC*(*r*) of each pair of signals *r*_1_ and *r*_2_ (see Eq. 3 in Methods and Eqs.

A.23 and A.24 in Appendix A.3). In [28], it was analytically shown that power correlation depends on signal coherence, cokurtosis, and conjugate-coherence. In particular, co-occurring bursts in neuronal activity, statistically measured by cokurtosis, are relevant for our discussion of SNEs. We computed the pSGDC and cokurtosis (Eq. A.26) measures on our MEG dataset by using the whole time series of the brain regions taken in pairs (i.e., non-time-resolved approach). As a result, we found that the pSGDC measure and the cokurtosis disclose a similar correlation degree with the observed co-activations topography (compare Figs. 7F and 7I) and generates statistics that are lost in the A- and B-surrogates (see Figs. 7E and 7H). Linear correlations between topographies: Co-activations vs pSGDC, *r* = 0.881, *P <* 0.001 (Fig. 7F). Cokurtosis vs pSGDC, *r* = 0.848, *P <* 0.001 (Fig. 7I). Co-activations vs Cokurtosis, *r* = 0.937, *P <* 0.001 (Figs. 7B,H). Co-activations vs Pairwise Pearson’s correlation, *r* = 0.612, *P <* 0.001 (Figs. 7B and A.1B). The statistical significance of these linear correlations was assessed by using the Student’s t distributions of the two-tailed hypothesis test under the null hypothesis that the correlation is zero.

The pSGDC measure quantifies the co-occurrence of above-threshold bursts mainly associated with SGDC in the alpha band, whereas cokurtosis assesses the presence of both oscillatory and non-oscillatory co-burstiness across brain regions. Importantly, the analytical framework proposed in this work based on the *SGDC*(*r*), *SGDC*(*ω*) and *pSGDC*(*r*_1_, *r*_2_) measures, admits relevant signal-level mechanistic interpretations linking the Fourier oscillatory constituents of the brain activity and SEs. Note that the latter is less evident when considering measures based on higher-order statistical moments like the kurtosis and cokurtosis. Specifically, using the group delay-domain representation, one can quantify the group delay consistency of the spectral (Fourier) constituents of the signals of interest (via the SGDC measures) to predict the emergence of SEs (without doing any explicit computation in the time-domain). This prediction linking the oscillation and time-domains can not be done by higher-order statistical moments like the kurtosis and cokurtosis, mainly because they operate exclusively in the time-domain. Therefore, the SGDC framework provides a deeper understanding of the link between the oscillation-domain (Fourier representation) and the emergence of transient, salient fluctuations in the time-domain. Thus, the SEs co-activation pattern reproduced by the pSGDC measure (see Figs. 7A-F) can be mechanistically segregated in two components: 1) the results associated with the *SGDC*(*r*) measure (Fig. C.11C) supporting the emergence of local above-threshold fluctuations via SGDC mainly in the alpha band, and 2) the results associated with the *SGDC*(*ω*) measure (Fig. C.11D) supporting the co-occurrence of above-threshold alpha bursts across brain regions (i.e., transient cross-regional coherence around the alpha band). We speculate that component 1 can be interpreted as an entrainment mechanism that produces transient synchronization of the oscillatory activity of neuronal populations around specific frequency bands (local cross-frequency synchronization), whereas component 2 can be associated with long-range interaction mediated by transient cross-regional coherence in NOs.

In summary, these results suggest that a) spectral group delay consistency in specific narrow frequency bands (as assessed by the *SGDC*(*r*) measure), b) transient cross-regional coherent NOs (intra-frequency coherence across brain regions assessed by the *SGDC*(*ω*) measure) and c) BAA, are all key ingredients for the emergence of realistic SEs. In particular, the (pairwise) long-range interactions mediated by oscillatory SNEs can be effectively quantified using the *pSGDC*(*r*_1_, *r*_2_) measure.

## 4. DISCUSSION

Frequency-domain representation of signals, via Fourier transforms (e.g., DFT), have been extensively used for decades in many neuroscience fields to analyze neuronal and brain activities across several spatiotemporal scales. Regardless of the functional significance of neural oscillations, if any, the Fourier basis functions provide an arguably good characterization of the rhythmic components observed in the brain activity. In this study, we used the complex baseband representation of signals, based on the Fourier theory, to analytically define the spectral group delay consistency (SGDC) as a novel conceptualization linking SEs with the signals’ spectral content. Importantly, the signal-level analytical framework associated with the SGDC concept allowed us to provide a unifying rationale for the emergence of salient local and large-scale events from the Fourier oscillatory constituents of the brain activity. First, the analytical arguments described in the Sections 3.4, 3.5 and Appendix A.2 point out that in order to observe realistic local above-threshold fluctuations, the spectral components constituting the brain signals must disclose a certain degree of cross-frequency coherence as assessed by the *SGDC*(*r*) measure. Second, in Sections 3.4 and Appendix A.4 we analytically showed that A- and B-surrogates failed to reproduce realistic SEs mainly because the phase randomization reduces the SGDC across frequency bands in each brain region, which impairs the burstiness of each signal (occurrence of local above-threshold fluctuations). In the case of the A-surrogates the phase randomization also reduces the SGDC across brain regions in each frequency band, which impairs the synchronization of above-threshold fluctuations across brain regions. Third, in Section 3.1 we showed that the spectral signature in the alpha band disclosed by the averaged ESM of cluster 2 SEs constitutes relevant evidence linking the observed SEs with NOs. Importantly, in Sections 3.6 and Appendix C.2, we demonstrated that the synchronization of above-threshold alpha bursts across brain regions can be described at the signal-level by the SGDC mechanism. Specifically, we showed that the SNEs disclosing the alpha spectral signature in the average ESM (see cluster 2 in Fig. C.11B) also disclose an increase of transient cross-regional coherence around the alpha band, as quantified by the *SGDC*(*ω*) measure (see cluster 2 in Fig. C.11D). Of note, the *SGDC*(*ω*) measure effectively captures transient, cross-regional coherent NOs associated with SNEs, a phenomenon that traditional coherence metrics, such as the Phase Locking Value (PLV), fail to detect (see Figs. A.5 and A.6). Thus, we combine analytical arguments, based on the SGDC framework, with experimental evidence obtained using novel tools like the ESM and SGDC measures, to provide a more direct and generative link for NOs (e.g., alpha oscillations) role in the coordination of SNEs observed in spontaneous MEG activity. This moves beyond mere correlation or characterization to offer a plausible generative model for SNEs as spatiotemporal cascades of above-threshold fluctuations associated with phase-structured NOs. Fourth, the SGDC conceptualization allowed us, via the *pSGDC*(*r*_1_, *r*_2_) measure, to account for both the co-activation pattern of brain avalanches and cokurtosis in terms of the coherence of the signals’ spectral content, therefore, advancing our understanding of the signal-level mechanisms of long-range communication. The empiric, modeling and analytical results presented in this work guided us to identify the essential building blocks underlying the emergence of realistic SEs as observed in our MEG dataset, which can be summarized as follows:

1. Spectral group delay consistency. This feature provides a signal-level mechanism for the emergence, in a single brain region (i.e., locally), of transient above-threshold fluctuations associated with an specific frequency band (e.g., alpha bursts). We speculate that the SGDC (e.g., bounded phase differences across spectral components within a narrowband) may be associated with the presence of mesoscopic neural oscillators that are not tightly tuned. We hypothesize that different brain regions may host mesoscopic oscillators disclosing rhythmic (likely non-sinusoidal) dynamics whose fundamental frequencies span a quasi-continuum within a given frequency band (e.g., alpha band), rather than clustering around a single sharply defined value. Thus, the linear superposition of these rhythmic components with slightly different frequencies within a narrowband (e.g., alpha range) could support the emergence of SEs via the SGDC signal-level mechanism.
2. Transient cross-regional coherent alpha oscillations. This feature is associated with the transient synchronization of the above-threshold alpha bursts across brain regions, giving rise to the SNEs producing the alpha spectral signature in the ESM (i.e., cluster 2 SEs). This type of SEs may be associated with a long-range interaction mechanism mediated by specific NOs taking place in a transient manner (i.e., transient CTC).
3. BAA. This feature is associated with the emergence of non-oscillatory above-threshold fluctuations occurring in an aperiodic manner, mainly related to the short-lived SEs with no characteristic spectral signature in the ESM (i.e., cluster 1 SEs). We hypothesize that the close relationship between cluster 1 SEs and arrhythmic broadband spectral features implies that cluster 1 SEs may play a more local role, linked either to local excitation-inhibition balance or to critical dynamics [43].

Linking the presence of SEs to the group delay consistency across the Fourier oscillatory components of the brain activity is a relevant result of this study implying that SEs might mediate interactions across both frequency bands and brain regions as discussed above. In this regard, the CTC hypothesis posits that neural communication is facilitated by the presence of synchronized (steady) oscillations across brain regions. Our results extend the CTC hypothesis by showing that long-range interaction through specific NOs may take place in a transient manner via SNEs (i.e., transient CTC). Indeed, our results suggest that the large-scale spreading of transient alpha bursts is associated with SNEs. As a conclusion, this evidence suggests that transient cross-regional coherence associated with the occurrence of SEs disclosing the spectral signature in the alpha band (i.e., cluster 2 SEs), may play a functional role as a long-range interaction mechanism in the resting human brain.

One of the main limitations of this study is related to the uncertain capability of our dataset to accurately identify deep brain sources along the cortical surface, mainly due to the ill-posed nature of the source-reconstructed MEG data. In order to address this issue, we re-computed the analysis of SEs presented above, but this time excluding the deep sources. It was found that all the conclusions and, in particular, all the characteristics of the observed SEs remain essentially unaltered when the deep sources are excluded from the SE analysis (see Appendix D). Specific analyses demonstrating that volume conduction alone is unlikely to account for the cascade of above-threshold fluctuations (i.e., SNEs) observed in our empirical MEG dataset have been presented and discussed in a previous publication [63]. The spatial leakage analyses and the full discussion can be accessed via this link: https://elifesciences.org/articles/67400/peer-reviews#content

## 5. CONCLUSION

In this work we provided a detailed analytical description of the mechanisms underlying the emergence of SEs from NOs and BAA co-existing in the human brain. The proposed analytical arguments were tested and confirmed using local and large-scale numerical models together with experimental MEG recordings obtained in healthy subjects during eyes-closed resting state. While previous studies have described SEs within the framework of neuronal avalanches, they often lacked a generative, signal-level account. Here, we bridge that divide by offering a mathematically grounded and empirically validated framework that accounts for oscillatory and aperiodic bursts perspectives on brain activity. We combine experimental evidence supported by a signal-level analytical framework and numerical simulations based on generative models to demonstrate that transient phase-structured alpha bursts, shaped by the SGDC mechanism, contribute to long-range coordination during rest. This extends the communication-through-coherence hypothesis into the transient domain. In summary, our multi-pronged approach, grounded in experimental evidence supported by analytical arguments and extensive model-based validation, enhances the robustness and interpretive depth of our results, offering a more comprehensive picture of how SEs arise from NOs and BAA as fundamental components of MEG activity during resting-state.

## ETHICAL PUBLICATION STATEMENT

We confirm that we have read the Journal’s position on issues involved in ethical publication and affirm that this report is consistent with those guidelines.

## DATA AND CODE AVAILABILITY

The MEG data are available upon request to the corresponding author (Pier-paolo Sorrentino), conditional on appropriate ethics approval at the local site. The availability of the data was not previously included in the ethical approval, and therefore data cannot be shared directly. In case data are requested, the corresponding author will request an amendment to the local ethical committee. Conditional to approval, the data will be made available. The code and simulated data that support the findings of this study are available from the corresponding author (Damián Dellavale), upon reasonable request. We are willing to provide technical support to investigators who express an interest in implementing the SGDC tools in other programming languages, integrate it in open-source software toolboxes, or use it for non-profit research activities.

## AUTHOR CONTRIBUTIONS

DD, GR and PS contributed to the conceptualization, methodology, formal analysis, writing the original draft and figures preparation. ETL, AR and PS contributed to the data acquisition, curation of the dataset and visual analysis of the recordings, review and editing the manuscript. PS contributed to the funding acquisition.

## FUNDING

### DECLARATION OF COMPETING INTERESTS

None of the authors has any conflict of interest to disclose.

## Appendix A. Supplementary analytical results

### Appendix A.1. Preservation of the Pearson’s cross-correlation in the B-surrogates

Let us start by considering the circular cross-correlation *R*_*xy*_(*t*) between the time series *x*(*t*) and *y*(*t*) representing the activities of two brain regions [45, pp. 571, 746],

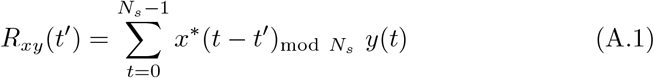

where (*x*(*t*), *y*(*t*)) ∈ ℝ are finite-length discrete time series having *N*_*s*_ time samples satisfying (*x*(*t*) = 0, *y*(*t*) = 0) ∀ 0 *> t > N*_*s*_ *−* 1, being *t* ∈ ℤ the discrete time index. By applying the Discrete Fourier Transform (DFT) 𝔉 {.} on both sides of Eq. A.1 we obtain [45, pp. 575, 746],

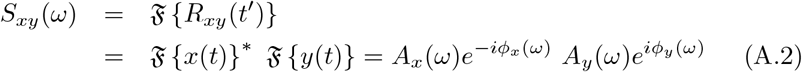

where *A*_*x*_(*ω*), *ϕ*_*x*_(*ω*) and *A*_*y*_(*ω*), *ϕ*_*y*_(*ω*) are the magnitude and phase angle of the DFT spectrum corresponding to the signals *x*(*t*) and *y*(*t*), respectively. The computation of surrogate time series involves the addition of random phases *θ*(*ω*) to the corresponding DFT spectra as follows,

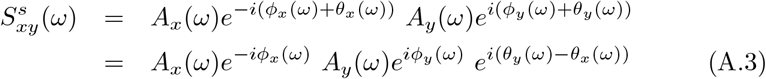

In the A.3, 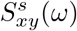 is the DFT of the circular cross-correlation associated with the surrogated time series 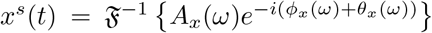 and 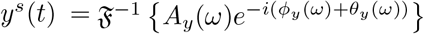., where 𝔉 ^−1^ {.} stands for the inverse DFT. In the particular case of the B-surrogates (see Section 2.8 in Methods) we add the same random phase-shift in all the brain regions, that is, *θ*_*x*_(*ω*) = *θ*_*y*_(*ω*) producing 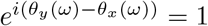 in the Eq. A.3. Under this condition, the Eqs. A.2 and A.3 becomes equivalent which in turn implies the equivalence between the circular cross-correlations associated with the true data and the B-surrogate,

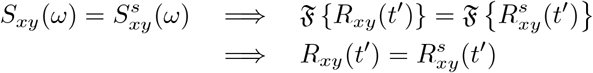

We confirmed this analytical results by computing the time-averaged functional connectivity as quantified by the pairwise Pearson’s correlation on our empirical MEG dataset and the corresponding A- and B-surrogates (seeSection 2.8 in Methods). Fig. A.1C shows the matrix resulting from computing the Pearson’s correlation on whole time series of the brain regions taken in pairs. Fig. A.1B shows the spatial profile obtained by averaging the Pearson’s correlation matrix across rows. Fig. A.1A displays the brain plots corresponding to the spatial profile of the Pearson’s correlation shown in Fig. A.1B. Importantly, Fig. A.1B shows that only B-surrogates reproduce the spatial profile of the Pearson’s correlation computed on the MEG data, hence, confirming that the pairwise Pearson’s correlation is preserved in the B-surrogates, and not in the case of A-surrogates.

**Figure A.1:**
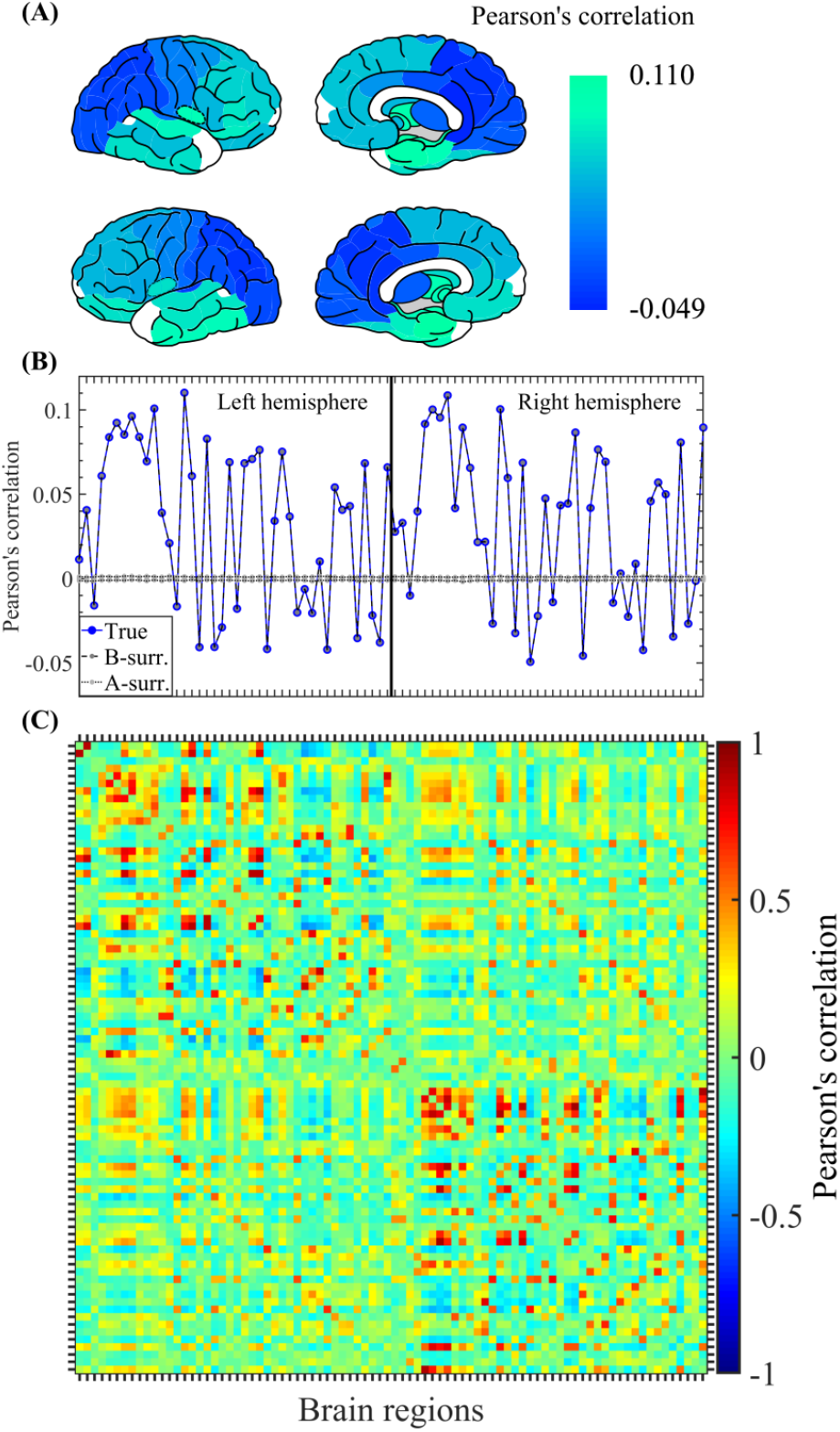
Pearson’s correlation pattern computed on whole time series of the brain regions taken in pairs. (A) Brain topographies corresponding to the Pearson’s correlation profile shown in panel B (blue markers). (B) Spatial profile showing the Pearson’s correlation (mean value across the 47 participants), i.e., the mean value computed on in each row of the Pearson’s correlation matrix. Note that the spatial profiles corresponding to the 100 B-surrogates (dark gray markers) overlap with the spatial profile associated with the true MEG data (blue markers). (C) Pearson’s correlation matrix (average across the 47 participants) obtained by computing the Pearson’s correlation on the whole time series of the brain regions taken in pairs. In panel B, the labels and ordering of the brain regions are the same as those shown in Fig. C.2.

### Appendix A.2. Oscillatory mechanisms underlying the emergence of local above-threshold fluctuations

In this section we provide a detailed description of the mechanism underlying the emergence of local above-threshold fluctuations from the Fourier oscillatory constituents of the brain activity. Our analysis start by projecting the brain signal of interest *x*(*t*) onto the Fourier basis functions using the Discrete Fourier Transform (DFT) equations [45, Chapters 8 and 10]. In doing so we are assuming that *x*(*t*) satisfies certain conditions so the resulting spectral estimates exist and are meaningful. Specifically, by considering finite-length time series constituted by *N*_*s*_ time samples, the existence of the DFT representation requires that *x*(*t*) is bounded (| *x*(*t*) < *M* ∈ ℝ ∀ 0 *> t > N*_*s*_ *−*1). Besides, the analyzed brain activity are in general nonstationary, that is, the time series *x*(*t*) can be represented as a sum of sinusoidal components with time-varying amplitudes, frequencies, or phases. In this regard, we consider a small enough number of time samples *N*_*s*_ such that the spectral characteristics of the signal *x*(*t*) can be assumed stationary during the analyzed time window. Thus, by considering *x*(*t*) ∈ ℝ being a finite-length discrete time series having an even number of time samples *N*_*s*_ and *x*(*t*) = 0 ∀ 0 *> t > N*_*s*_ *−*1, where *t* ∈ ℤ is the discrete time index. The analysis equation corresponding to the Discrete Fourier Transform (DFT) of *x*(*t*) can be written as follows [45, p. 561, Eq. (8.67)],

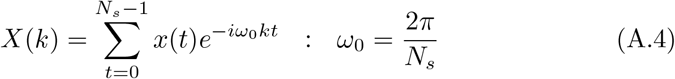

where *k* ∈ ℤ is the discrete frequency index, in general producing complex Fourier coefficients *X*(*k*) ∈ ℂ and *X*(*k*) = 0 ∀ 0 *> k > N*_*s*_ *−*1. Then, the synthesis equation associated with the inverse DFT (iDFT) is [45, p. 561, Eq. (8.68)],

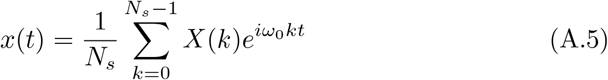

Taking into account that *X*(*k*) = |*X*(*k*)| *e*^*iϕ*(*k*)^ ∈ ℂ, the Eq. A.5 can be rewritten as,

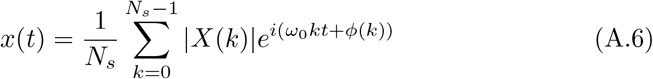

The core of the proposed conceptualization is to note that the Eq. A.6 can be expressed as a sum of (non-overlapping) pairwise adjacent spectral components as follows,

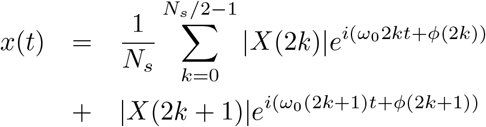

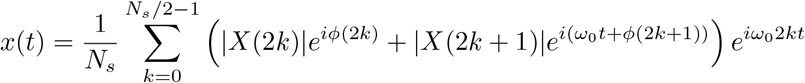

By defining the forward phase difference as Δ*ϕ*(2*k*) = *ϕ*(2*k* + 1) *− ϕ*(2*k*), and substituting *ϕ*(2*k* + 1) = *ϕ*(2*k*) + Δ*ϕ*(2*k*) in the previous equation we have,

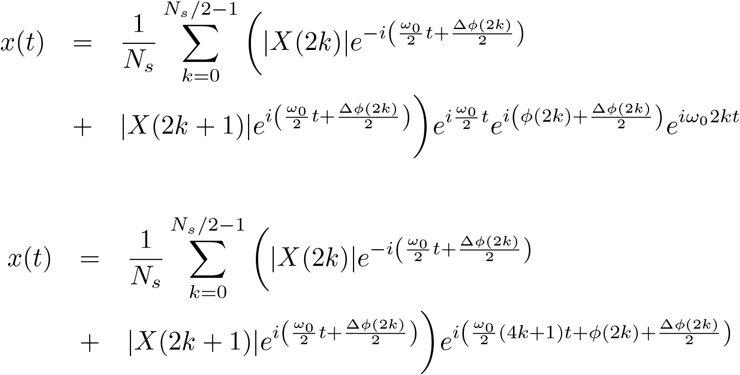

Then, by introducing in the previous equation the forward frequency difference Δ*ω* = *ω*_0_ (*k* + 1) − *ω*_0_ *k* = *ω*_0_, it results,

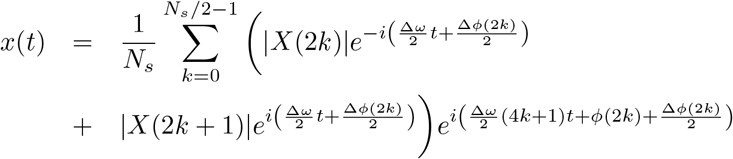

Taking out Δ*ω/*2 as a common factor we have,

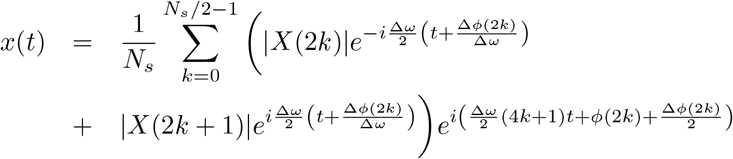

The rate of change of the phase with the frequency is associated with the group delay defined as *τ* (*k*) = *−*Δ*ϕ*(*k*)*/*Δ*ω*. Using this definition, the previous equation can be written as,

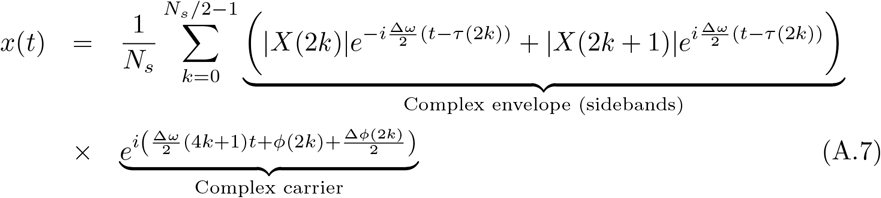

It is essential to note that in Eq. A.7, each (non-overlapping) pair of adjacent spectral components *X*(2*k*), *X*(2*k* + 1) can be interpreted as the sidebands of an amplitude modulated carrier at (4*k* + 1)Δ*ω/*2. Importantly, the frequency of the carrier (4*k* + 1)Δ*ω/*2 is a function of the frequency index *k*, that is, it depends on the particular pair of spectral components under consideration (*X*(2*k*), *X*(2*k* + 1)). However, the frequency of the modulating component is the same for all the pair of spectral components involved in Eq. A.7, i.e., it is independent of the frequency index *k* and only determined by the frequency resolution of the DFT as Δ*ω/*2 = *ω*_0_*/*2 (i.e., half the separation between the two sidebands). Another important characteristic of the representation given by the Eq. A.7 is that the frequencies associated with the complex envelopes (Δ*ω/*2) and with the complex carrier ((4*k* +1)Δ*ω/*2) satisfy the condition Δ*ω/*2 ≤ (4*k* + 1)Δ*ω/*2. In the telecom theory, a spectral profile satisfying these characteristics is known as the complex baseband representation of a band-limited signal (e.g., amplitude modulated signal) [45, Chapter 11.4.2, p. 796; 53, Chapter 4.1, p. 152; 26, Chapter A2.4, p. 725]. Accordingly, we refer to the Eq. A.7 as the inverse DFT based on the pairwise complex baseband representation of *x*(*t*). In line with this, the Eq. A.7 can be rewritten as a summation of amplitude modulated signals corresponding to each pair of adjacent spectral components as follows,

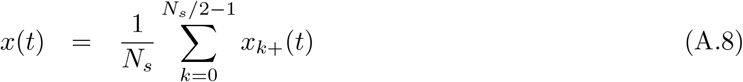

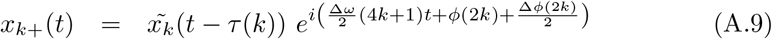

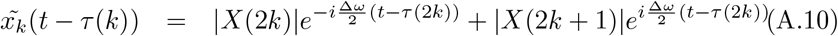

In the Eq. A.8, *x*_*k*+_(*t*) is the discrete time analytic signal (a.k.a., pre-envelope) corresponding to each amplitude modulated component constituting the original signal *x*(*t*), and it is defined in Eq. A.9. In the Eq. A.9, 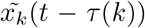 is the complex envelope of each amplitude modulated component constituting the original signal *x*(*t*), and it is defined in terms of the spectral components *X*(*k*) in the Eq. A.10. It is important to note that the alignment in time of the complex envelopes 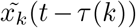 synthesizing the original signal *x*(*t*), via the Eq. A.8, is determined by the group delay *τ* (*k*).

The Eqs. A.7 - A.10 constitute a useful conceptualization linking the DFT and the complex baseband representation to account for the emergence of salient events from the Fourier oscillatory constituents of a band-limited signal. Due to the fact that the analysis proposed above is based on the DFT, in the case of *x*(*t*) ∈ ℝ the result of the summation in Eqs. A.7 and A.8 is guaranteed to be real valued. At the same time, this also restrict the validity of the analysis to harmonic spectral components *ω*_0_ *k* associated with the fundamental frequency *ω*_0_ = 2*π/N*_*s*_. Now we will present the general equations valid for all the cases, that is, harmonic (Δ*ω*(*k*) = cte, *ω*(*k* + 1)*/ω*(*k*) ∈ ℚ), non-harmonic (Δ*ω*(*k*) = cte, *ω*(*k* + 1)*/ω*(*k*) ∈ ℝ\ ℚ) and non-uniformly spaced (Δ*ω*(*k*) ;= cte) Fourier oscillatory components. Let us consider a real valued signal *x*(*t*) ∈ ℝ resulting from the linear superposition of an even number *N*_*s*_ of oscillatory components of arbitrary amplitude *A*(*k*), frequency *ω*(*k*) and phase *ϕ*(*k*).

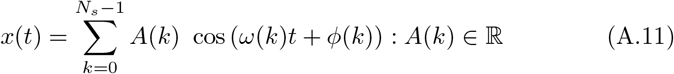

Since the Eq. A.11 is linear we can introduce the complex notation via the Euler’s formula as follows,

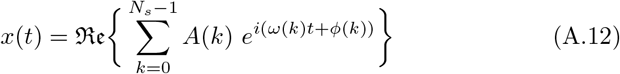

In the Eq. A.12, the operator *ℜ𝔢* . stands for “the real part of”. By following a similar procedure applied above on the Eq. A.6, the Eq. A.12 can be rewritten as follows,

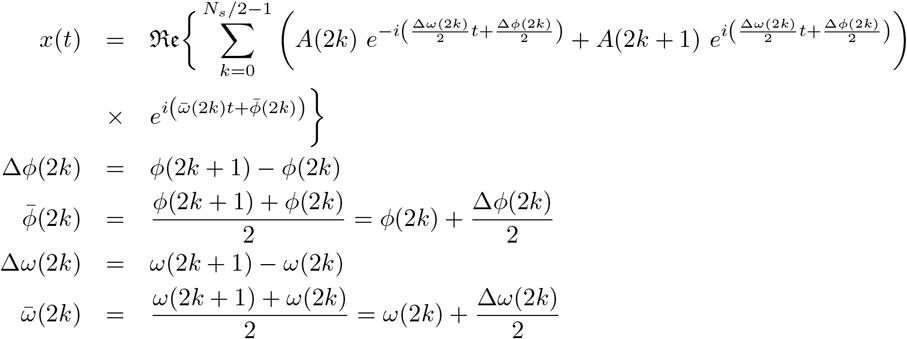

In this case the group delay is defined as 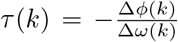, thus, the previous equation results,

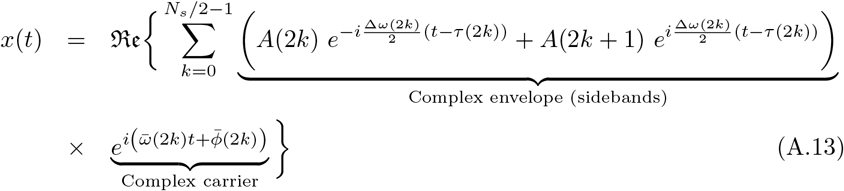

The Eq. A.13 is the pairwise complex baseband representation of the signal *x*(*t*). Provided that the frequencies associated with the complex envelopes (Δ*ω*(2*k*)*/*2) and the complex carrier 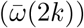 satisfy the condition 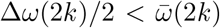, the Eq. A.13 can also be written as a summation of discrete time analytic signals *x*_*k*+_(*t*) associated with amplitude modulated signals corresponding to each pair of adjacent oscillatory components as follows,

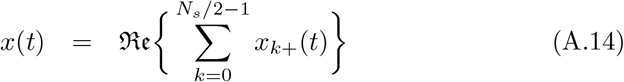

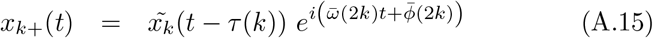

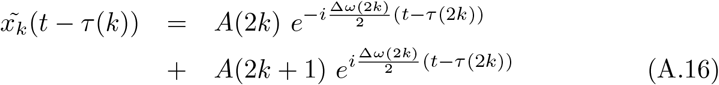

Similarly to the previous case the time alignment of the complex envelopes 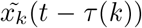 synthesizing the original signal *x*(*t*), via the Eq. A.14, is determined by the group delay *τ* (*k*).

In what follows we will use the Eq. A.13 to illustrate the role of the group delay in the emergence of above-threshold fluctuations from the oscillatory constituents of the synthetic signal *x*(*t*). As a first example, let us consider a spectral profile given by a set of constant-amplitude *A*(*k*) = *A* = 1 oscillatory components uniformly spaced 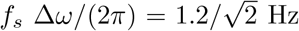 Hz and having non-harmonic frequencies *f*_*s*_ *ω*(*k*)*/*(2*π*) = 0.5 + *k f*_*s*_ Δ*ω/*(2*π*) ∈ [0.5 *−*5] Hz, where *f*_*s*_ = 1024 Hz is the sampling rate (see Figs. A.2A and A.2F). Accordingly, the Eq. A.13 becomes,

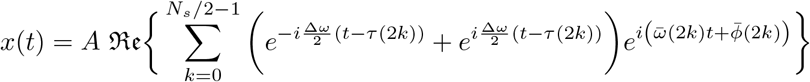

By using the Euler’s formula to rearrange the modulating factor, the previous equation results,

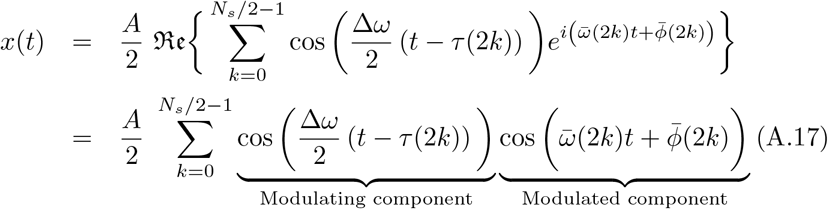

The Eq. A.17 explicitly shows that any pair of adjacent oscillatory components associated with the signal *x*(*t*) can be interpreted as an amplitude modulated signal with the same modulating function cos 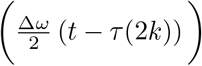 . The key concept here is to note that, when all the oscillatory components in Eq. A.17 are added together to synthesize the signal *x*(*t*) in the time-domain, the group delay *τ* will determine the time alignment of the modulating functions associated with each pair of adjacent oscillatory components. As a consequence, in the case of all the spectral components *A*(*k*) *e*^*i*(*ω*(*k*)*t*+*ϕ*(*k*))^ in Eq. A.12 having constant phase produces Δ*ϕ* = 0 ⇒ *τ* = −Δ*ϕ/*Δ*ω* = 0, hence, all the modulating functions cos 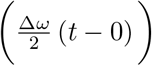 in Eq. A.17 will be aligned in time (at *t* = 0) giving rise to a sinc-like function representing the maximum amplitude excursion (i.e., a salient event) that can be elicited by the set of Fourier oscillatory components constituting the Eq. A.12. In the case of all the spectral components in Eq. A.12 having a phase proportional to the discrete frequency index *ϕ*(*k*) = −*τ*_0_ Δ*ω k* =⇒ Δ*ϕ*(*k*) = −*τ*_0_ Δ*ω*, results in a group delay which does not dependent on the frequency *τ* (*k*) = −Δ*ϕ*(*k*)*/*Δ*ω* = *τ*_0_, thus, in Eq. A.12 we obtain a modulating component cos 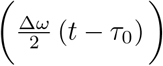 . That is, all the modulating functions will again be aligned in time producing the same salient event given by the sinc-like function as in the previous case but this time centered at *t* = *τ*_0_ (i.e., a time-shift, see Figs. A.2A-E). On the other hand, in the case of the phases associated with the spectral components in Eq. A.12 having a non-linear dependence with the discrete frequency index, e.g., *ϕ*(*k*) = −*τ*_0_ Δ*ω k*^2^ =⇒ Δ*ϕ*(*k*) = −*τ*_0_ Δ*ω*(2*k* + 1), the group delay results a function of the frequency *τ* (*k*) = *τ*_0_(2*k* + 1), hence, preventing the alignment in time of the modulating functions associated with each pair of adjacent spectral components cos 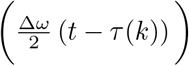 . In this case, the signal *x*(*t*) discloses sub-threshold excursions of amplitude (see Figs. A.2F-J). It is worth mentioning that in deriving the pairwise complex baseband representation of *x*(*t*) given by the Eqs. A.7 and A.13, we grouped the original spectral components (Eqs. A.5 and A.11) in subsets of (non-overlapping) pairs adjacent in frequency. The strategy of grouping the spectral components in subsets is necessary to obtain a representation based on a sum of complex envelopes modulating the complex carriers. Representations similar to those presented in the Eqs. A.7 and A.13 can be obtained by defining subsets containing more than 2 non-overlapping spectral components (not necessarily adjacent in frequency). However, our approach based on grouping adjacent spectral components in non-overlapping pairs discloses the following relevant features:

1. By defining subsets of 2 spectral components, we obtain the simplest complex envelopes characterized by a cos- or sin-like waveform shape (see the modulating component in the Eq. A.17 and the colored solid lines in Figs.A.2E and A.3E).
2. By defining pairs of spectral components adjacent in frequency, we maximize the waveform shape similarity among the resulting complex envelopes. In the case of uniformly spaced spectral components (Δ*ω* = cte), we obtain complex envelopes having the same time period 2*/*Δ*ω* (see the colored doted lines in Figs.A.2E and A.3E).
3. By defining pairs of spectral components adjacent in frequency, we also maximize the similarity among the resulting complex carriers (see the colored solid lines in Figs.A.2E and A.3E).

Taking together, these features are of particular importance to support the link between the spectral group delay consistency (SGDC) defining the time alignment of the modulating components (complex envelopes) with the constructive interference of the modulated components (complex carriers), which in turn lead to the occurrence of salient events. As a conclusion, the results described above in connection with the Eqs. A.7, A.13, show that the emergence of above-threshold fluctuations in the signal *x*(*t*) is related to the consistency of the group delay *τ* (*k*) across the discrete frequency values *k*. That is, the occurrence of salient events is supported by a slowly varying group delay as a function of the frequency, and this hold true for harmonic, non-harmonic and also for non-uniformly spaced Fourier oscillatory constituents of the signal under analysis.

**Figure A.2:**
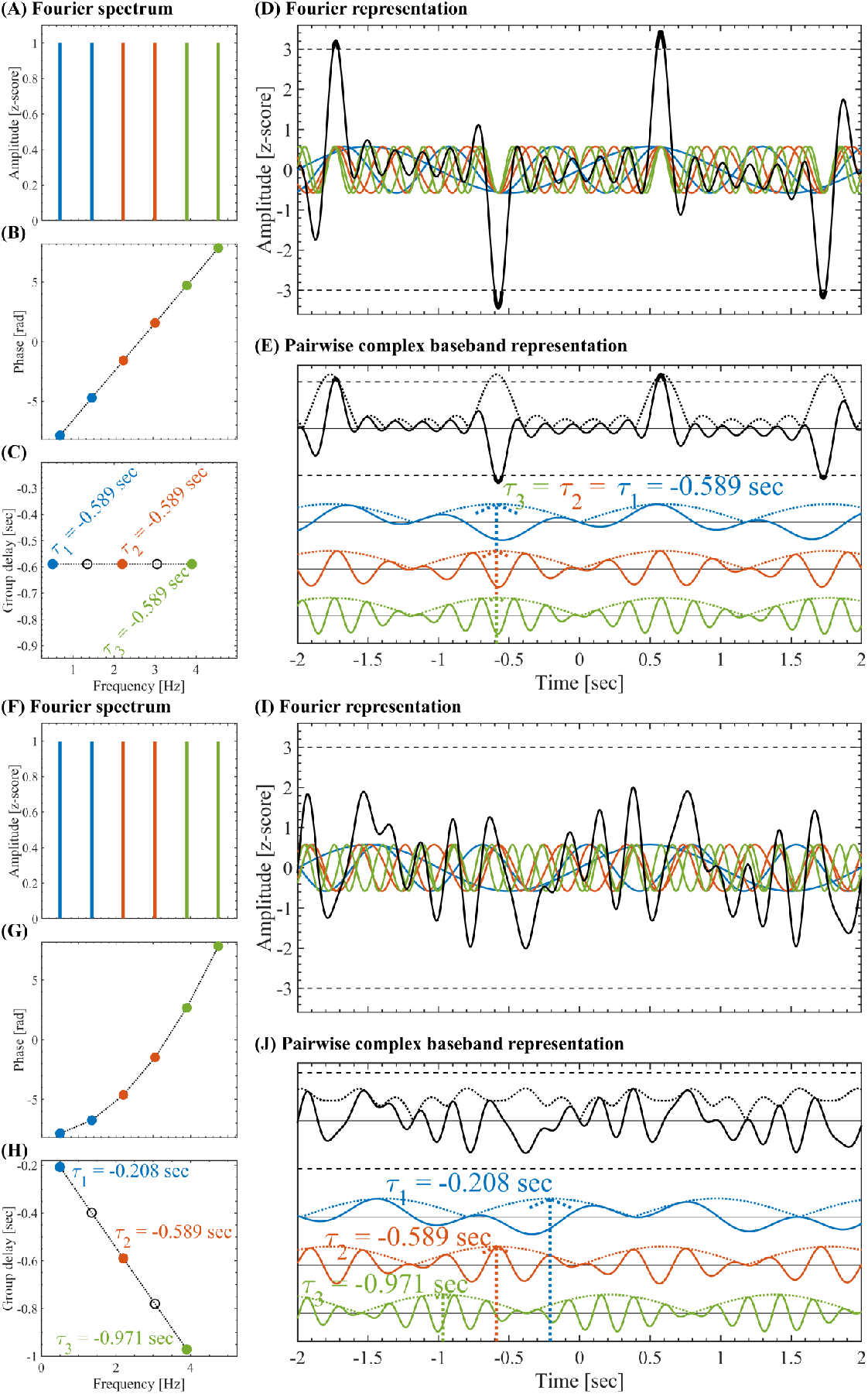
Pairwise complex baseband representation for a set of oscillatory components with *A*_*k*_ = cte. (A) Set of constant-amplitude *A*(*k*) = 1 oscillatory components uniformly spaced 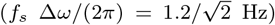 and having non-harmonic frequencies *f*_*s*_ *ω*(*k*)*/*(2*π*) = 0.5 + *k f*_*s*_ Δ*ω/*(2*π*) ∈ [0.5 *−*5] Hz, where *f*_*s*_ = 1024 Hz is the sampling rate. The pairwise complex baseband representation (Eq. A.13) was obtained by grouping the oscillatory components in adjacent non-overlapping pairs color-coded in blue, red and green. (B) Phases *ϕ*(*k*) having a linear dependence as a function of the frequency within the range *ϕ*(*k*) ∈ 2.5 [−*π, π*]. (C) Group delay *τ* (*k*)*/f*_*s*_ = −Δ*ϕ*(*k*)*/*(*f*_*s*_ Δ*ω*) for the pairs of adjacent oscillatory components. The color-coded filled markers correspond to the *τ* (2*k*)*/f*_*s*_ values, and the black empty markers correspond to *τ* (2*k* + 1)*/f*_*s*_ values (see Eq. A.13). (D) Z-scored signals. The solid color-coded lines represent the individual oscillatory components, the solid black line is the resulting signal *x*(*t*), the horizontal dashed black lines indicate the threshold at |*z*| = 3. (E) Pairwise complex baseband representation. The solid color-coded lines represent the individual amplitude modulated signals (pairs of adjacent oscillatory components), the solid black line is the resulting signal *x*(*t*), the color-coded and black doted lines are the corresponding amplitude envelopes. (F - J) Same as panels (A - E), this time with phases *ϕ*(*k*) having a quadratic dependence as a function of the frequency within the range *ϕ*(*k*) ∈ 2.5 [*−π, π*] (see panel G).

The group delay is defined in terms of the rate of change of the phase with the frequency, being independent on the amplitude of the spectral components. As a consequence, the consistency of the spectral group delay as a mechanism supporting the emergence of salient events is also valid for spectral profiles other than the constant-amplitude spectrum shown in the Fig. A.2. The Fig. A.3 shows the results for a spectral profile given by a set of (uniformly spaced) non-harmonic oscillatory components with amplitudes 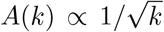, that is, the power of the spectral components *A*^2^(*k*) is proportional to 1*/k* (see Figs. A.3A and A.3F). Figs. A.3A-E show the case in which the phases *ϕ*(*k*) of the spectral components *A*(*k*) *e*^*i*(*ω*(*k*)*t*+*ϕ*(*k*))^ in Eq. A.12 are randomly distributed in a very small range around zero (*ϕ*(*k*) ∈ [*−π/*10, *π/*10]). Under this condition, the pairwise complex baseband representation (Eq. A.13) shown in the Fig. A.3E is constituted by amplitude modulated signals highly aligned in time. As a consequence, prominent salient events can be distinguished in the resulting signal (see solid black line in panels D and E of Fig. A.3). On the other hand, Figs. A.3A-E show the case in which the phase values *ϕ*(*k*) are randomly distributed in a wider range *ϕ*(*k*) ∈ [*−π, π*]. Under this condition, the pairwise complex baseband representation (Eq. A.13) shown in the Fig. A.3J is constituted by amplitude modulated signals non-aligned in time. As a consequence, the resulting signal *x*(*t*) only discloses sub-threshold excursions of amplitude (see solid black line in panels I and J of Fig. A.3).

**Figure A.3:**
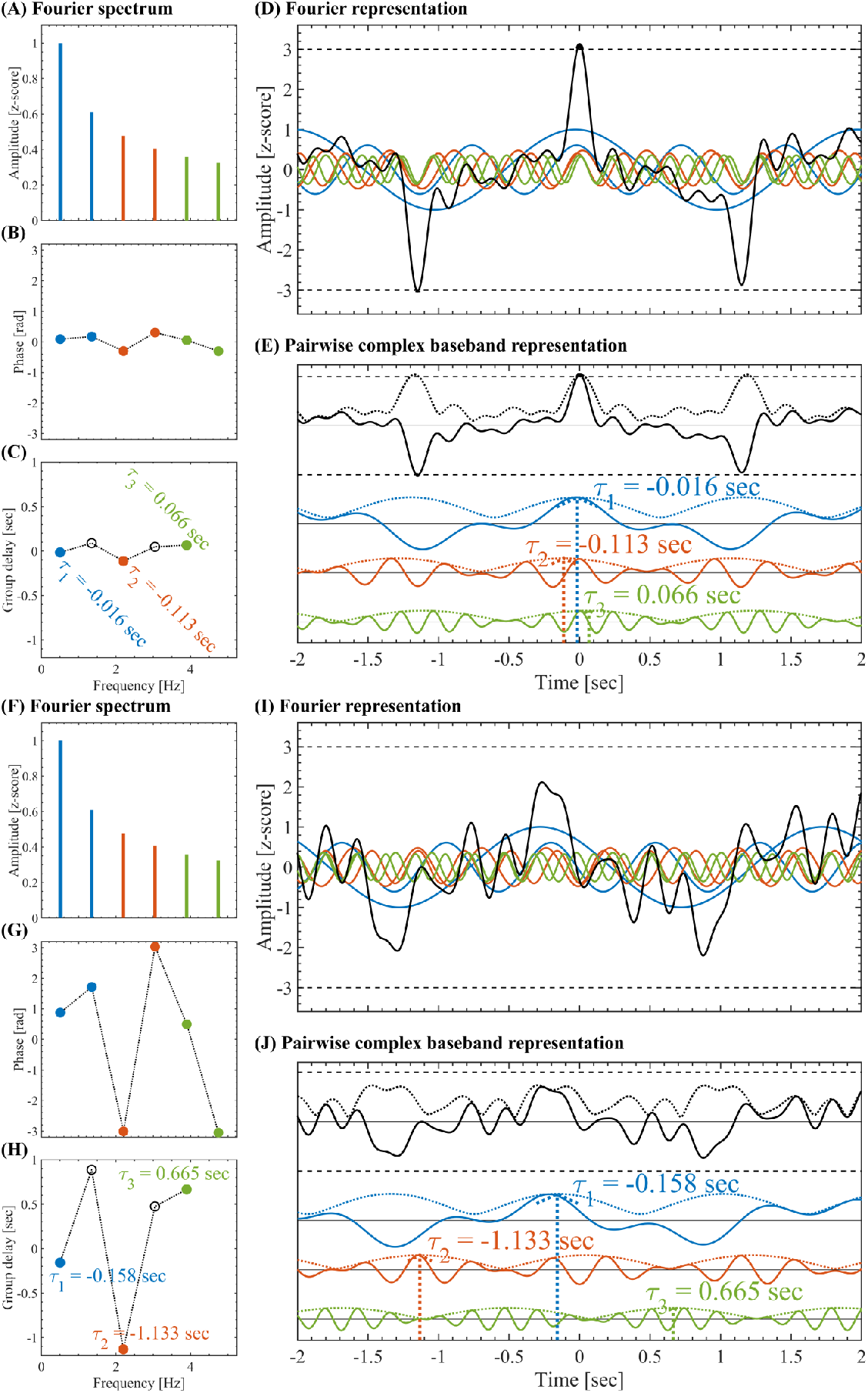
Pairwise complex baseband representation for a set of oscillatory components with 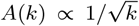. (A) Set of non-constant amplitude 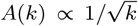 oscillatory compo-nents uniformly spaced 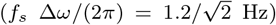 and having non-harmonic frequencies *f*_*s*_ *ω*(*k*)*/*(2*π*) = 0.5 + *k f*_*s*_ Δ*ω/*(2*π*) ∈ [0.5 *−* 5] Hz, where *f*_*s*_ = 1024 Hz is the sampling rate. The pairwise complex baseband representation (Eq. A.13) was obtained by grouping the oscillatory components in adjacent non-overlapping pairs color-coded in blue, red and green. (B) Phases *ϕ*(*k*) randomly distributed within a very small range around zero (*ϕ*(*k*) ∈ [−*π/*10, *π/*10]). (C) Group delay *τ* (*k*)*/f*_*s*_ = −Δ*ϕ*(*k*)*/*(*f*_*s*_ Δ*ω*) for the pairs of adjacent oscillatory components. The color-coded filled markers correspond to the *τ* (2*k*)*/f*_*s*_ values, and the black empty markers correspond to *τ* (2*k* + 1)*/f*_*s*_ values (see Eq. A.13). (D) Z-scored signals. The solid color-coded lines represent the individual oscillatory components, the solid black line is the resulting signal *x*(*t*), the horizontal dashed black lines indicate the threshold at |*z*| = 3. (E) Pairwise complex baseband representation. The solid color-coded lines represent the individual amplitude modulated signals (pairs of adjacent oscillatory components), the solid black line is the resulting signal *x*(*t*), the color-coded and black doted lines are the corresponding amplitude envelopes. (F - J) Same as panels (A - E), this time the phases *ϕ*(*k*) are randomly distributed within the range *ϕ*(*k*) ∈ [−*π, π*] (see panel G).

In summary, the analytical arguments presented above, condensed in the Eqs. A.7 - A.10 and A.13 - A.16, allowed us to identify the consistency of the group delay across the spectral components as a mechanism accounting for the emergence of above-threshold fluctuations from the Fourier oscillatory constituents of the activity associated with a single brain region. In the next section we describe the signal processing tools proposed to quantify the SGDC in empirical data.

### Appendix A.3. Measures to assess the spectral group delay consistency

The analytical arguments presented in the Appendix A.2 have profound consequences regarding the interpretation of the experimental results in connection with the emergence of salient events from NOs and broadband 1*/f* activity. Specifically, the pairwise complex baseband representation of band-limited signals (Eqs. A.7 - A.10 and A.13 - A.16), explicitly shows that the mechanism underlying the emergence of above-threshold fluctuations in a signal *x*(*t*) can be understood in terms of the consistency of the group delay across the Fourier oscillatory constituents of the signal (see the complex envelopes 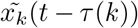 in Eqs. A.10 and A.16). By considering a multi-regional approach, the pairwise complex baseband representation can be applied on the activity *x*_*r*_(*t*) of each brain region *r*, to obtain complex envelopes of the form 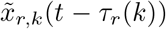. Here we recall that 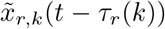 determine the envelopes of the individual amplitude modulated signals constituting the signal *x*_*r*_(*t*) (see the solid and doted color-coded curves in the Figs. A.2E,J and A.3E,J). Hence, the consistency of the spectral group delay *τ*_*r*_(*k*) determines the synchronization of the complex envelopes 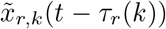 across both frequency values *ω*(*k*) and brain regions *r*. In what follows we describe the proposed measures designed to quantify the spectral group delay consistency (SGDC) in experimental data across either frequency values and/or brain regions. In order to simplify the notation, in the rest of this section we will use *ω* instead of the discrete frequency index *k*, implicitly assuming that *ω* = *ω*(*k*). In the most general case, the spectral group delay can be estimated as *τ*_*r*_(*ω*) = *−*Δ*ϕ*_*r*_(*ω*)*/*Δ*ω*(*ω*), where Δ*ϕ*_*r*_(*ω*) and Δ*ω*(*ω*) are the incremental phase and incremental frequency between adjacent spectral components associated with the activity *x*_*r*_(*t*) of the brain region *r*, respectively. Let us consider first the particular case of Δ*ω*(*ω*) = *−*Δ*ω* = const, in which the group delay results *τ*_*r*_(*ω*) ∝ − Δ*ϕ*_*r*_(*ω*). Therefore, the SGDC can be simply assessed via the Euler’s transform of the incremental phase as follows,

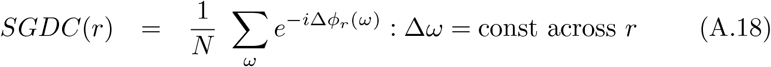

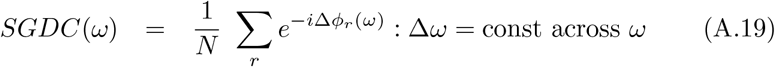

The modulus of Eqs. A.18 and A.19 satisfies,

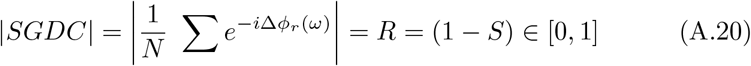

In the Eqs. A.18, A.19 and A.20, *N* is the number of either frequency values or brain regions as appropriate, *R* is the resultant vector length and *S* is the circular variance [9]. The Eq. A.20 explicitly shows that the SGDC is assessed as one minus the circular variance of the incremental phase. The definition of the SGDC measures given in the Eqs. A.18, A.19 and A.20 should not be confused with the traditional measure for quantifying coherence known as Phase Locking Value (PLV) [65, 32]. Specifically, the SGDC measures as defined in the Eqs. A.18, A.19 and A.20 assess the consistency of the incremental phase Δ*ϕ*_*r*_(*ω*) across the frequency values *ω*. In contrast, the PLV assesses the consistency of phase difference across the time samples, where the phase difference is computed between two phase time series corresponding to two specific frequency bands in the same or different brain regions [65, 32]. As stated in the Eq. A.18, the *SGDC*(*r*) is a bounded measure in the range [0, 1] and quantifies how much the group delay varies across the spectral components conforming the activity of interest *x*_*r*_(*t*). On the one hand, constant group delay values *τ*_*r*_(*ω*) ∝ *−* Δ*ϕ*_*r*_(*ω*) across the spectral components produce |*SGDC*(*r*)| ≈ 1 indicating a high SGDC, which is associated with high burstiness of the signal *x*_*r*_(*t*) (see Figs. A.2A-E and A.3A-E). On the other hand, in the case of group delay values varying randomly (or non-linearly) across the spectral components produces |*SGDC*(*r*)| ≈ 0 indicating low SGDC associated with low burstiness of the signal *x*_*r*_(*t*) (see Figs. A.2F-J and A.3F-J). Similarly, the *SGDC*(*ω*) defined in the Eq. A.19 is a bounded measure in the range [0, 1] and quantifies how much the spectral group delay at a given frequency *ω*, varies across the brain regions *r*. On the one hand, constant group delay values *τ*_*r*_(*ω*) ∝ *−* Δ*ϕ*_*r*_(*ω*) across the brain regions produce |*SGDC*(*ω*)| ≈ 1 indicating a high group delay consistency, which is associated with high cross-regional synchronization of the bursts at a given frequency *ω*. On the other hand, in the case of group delay values varying randomly (or non-linearly) across the brain regions produces |*SGDC*(*ω*)| ≈ 0 indicating low group delay consistency associated with low cross-regional synchronization of the bursts at a given frequency *ω*. Now we will consider the more general case in which Δ*ω*(*k*) ≠ cte. In line with the previous analysis, the SGDC measures can be defined in terms of the linear variance of the group delay Var(*τ*) as follows,

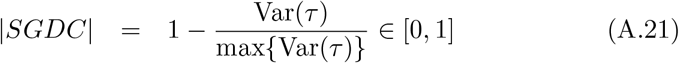

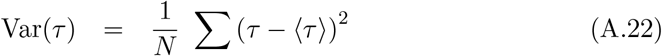

In the Eq. A.22, the mean group delay value ⟨*τ* ⟩ and the the sum associated with the linear variance Var(*τ*) are computed across the *N* frequency values *ω* or brain regions *r* in which case the Eq. A.21 produces |*SGDC*(*r*)| or |*SGDC*(*ω*)|, respectively. Importantly, the Eqs. A.18, A.19 and A.21 constitute an specialized framework to quantify the emergence of large-scale bursts (i.e., salient network events) from the brain activity. That is, the *SGDC*(*r*) assesses the emergence of local above-threshold fluctuations from the spectral components constituting the activity of a single brain region, whereas the *SGDC*(*ω*) measure quantifies the synchronization of the above-threshold bursts across brain regions. In line with this, we introduce the pairwise spectral group delay consistency (pSGDC) to quantify the burstiness and cross-regional bursts synchronization in a single measure. In the case of Δ*ω*(*ω*) = Δ*ω* = const, the pSGDC is defined as follows,

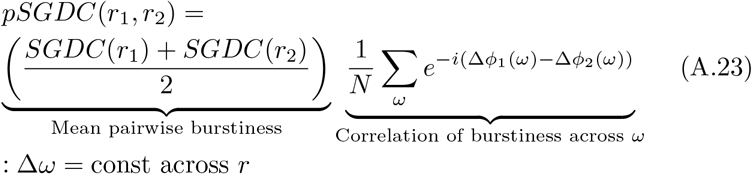

In the Eq. A.23, the quantities *SGDC*(*r*_1_) and *SGDC*(*r*_2_) are computed using the Eq. A.18. In the case of Δ*ω*(*ω*) ≠ cte the pSGDC is defined as follows,

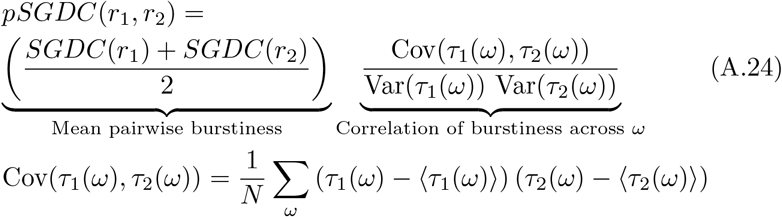

In the Eq. A.24, the quantities *SGDC*(*r*_1_) and *SGDC*(*r*_2_) are computed using the Eqs. A.21 and A.22. Besides, the quantities Var(*τ*_1_(*ω*)) and Var(*τ*_2_(*ω*)) are computed using the Eq. A.22. In both cases the sum associated with the Eq. A.22 is computed over the frequency values *ω*. The Eqs. A.23 and A.24 show that the *pSGDC*(*r*_1_, *r*_2_) is a linear measure conformed by a factor quantifying the cross-regional correlation between the group delays across the frequency values, weighted by a coefficient quantifying the burstiness of the two involved brain regions (*r*_1_, *r*_2_). Importantly, we found that the pSGDC performs similarly to the cokurtosis (fourth standardized cross central moment) [28] in reproducing the observed salient events topographies and co-activation patterns (see Fig. 7 in Section 3.6 of the main text). This is particularly interesting taking into account that these two non-time-resolved measures (i.e., computed on the whole time series) effectively reproduce the salient events topographies through two different approaches. That is, the cokurtosis is a non-linear time-domain measure, whereas the pSGDC is a linear measure entirely based on the frequency-domain. Moreover, the pSGDC and cokurtosis disclose a better performance to reproduce the observed salient events topographies and co-activation patterns when compared to the kurtosis (scaled version of the fourth central moment) and the Pearson’s linear correlation (see discussion in Section 3.6 of the main text). These results are consistent with the fact that kurtosis measures the presence of outliers (tails of the distribution of amplitude values) and the Pearson’s correlation coefficient the linear correlations between the two time series. On the other hand, pSGDC and cokurtosis measures quantify these two features simultaneously. In this work the kurtosis (K) and the cokurtosis (CK) were assessed via the following standard unbiased estimators,

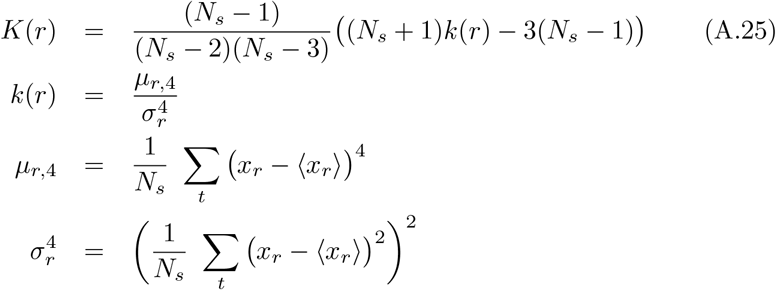

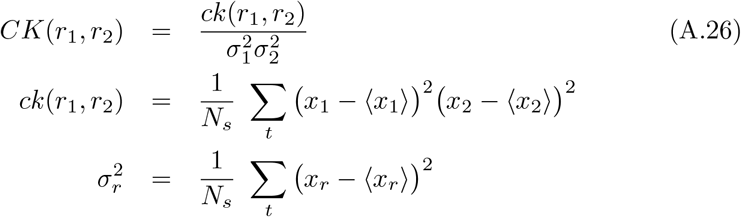

In the Eqs. A.25 and A.26, *N*_*s*_ is the number of time samples and ⟨. ⟩ stands for mean value across the time samples.

In the rest of this section, we present illustrative examples using the Eqs. A.18 and A.19 on synthetic multi-channel bursts emerging from narrowband oscillatory activity. Fig. A.4 shows the |*SGDC*(*r*) | computed using the Eq. A.18 for three time series synthesized using the Eq. A.11. In each channel, the signal was synthesized by the linear superposition of 10 sinusoidal tones with uniformly spaced frequencies (Δ*ω* = const) in the range *f*_*s*_ *ω/*(2*π*) ∈ [0.5 *−*3] Hz. In channels 1 and 2, the phase of the tones were set as a quadratic function of the frequency within the range *ϕ*_1_(*ω*) ∝ 2*πω*^2^ ∈ [*−*2*π*, 2*π*] and *ϕ*_2_(*ω*) ∝ *πω*^2^ ∈ [*− π, π*], respectively. In channel 3, the phase of the tones were set as a linear function of the frequency within the range *ϕ*_3_(*ω*) ∝ *πω* ∈ [*−π, π*]. Fig. A.4B shows that the higher the burstiness (i.e., amplitude of the transient fluctuations) disclosed by the resulting signal (see solid black line in the Fig. A.4A), the higher the |*SGDC*(*r*)| value. The channel 3, corresponding to the tones having a linear phase dependence with the frequency, discloses the maximum |*SGDC*(*r*)| ≈ 1.

**Figure A.4:**
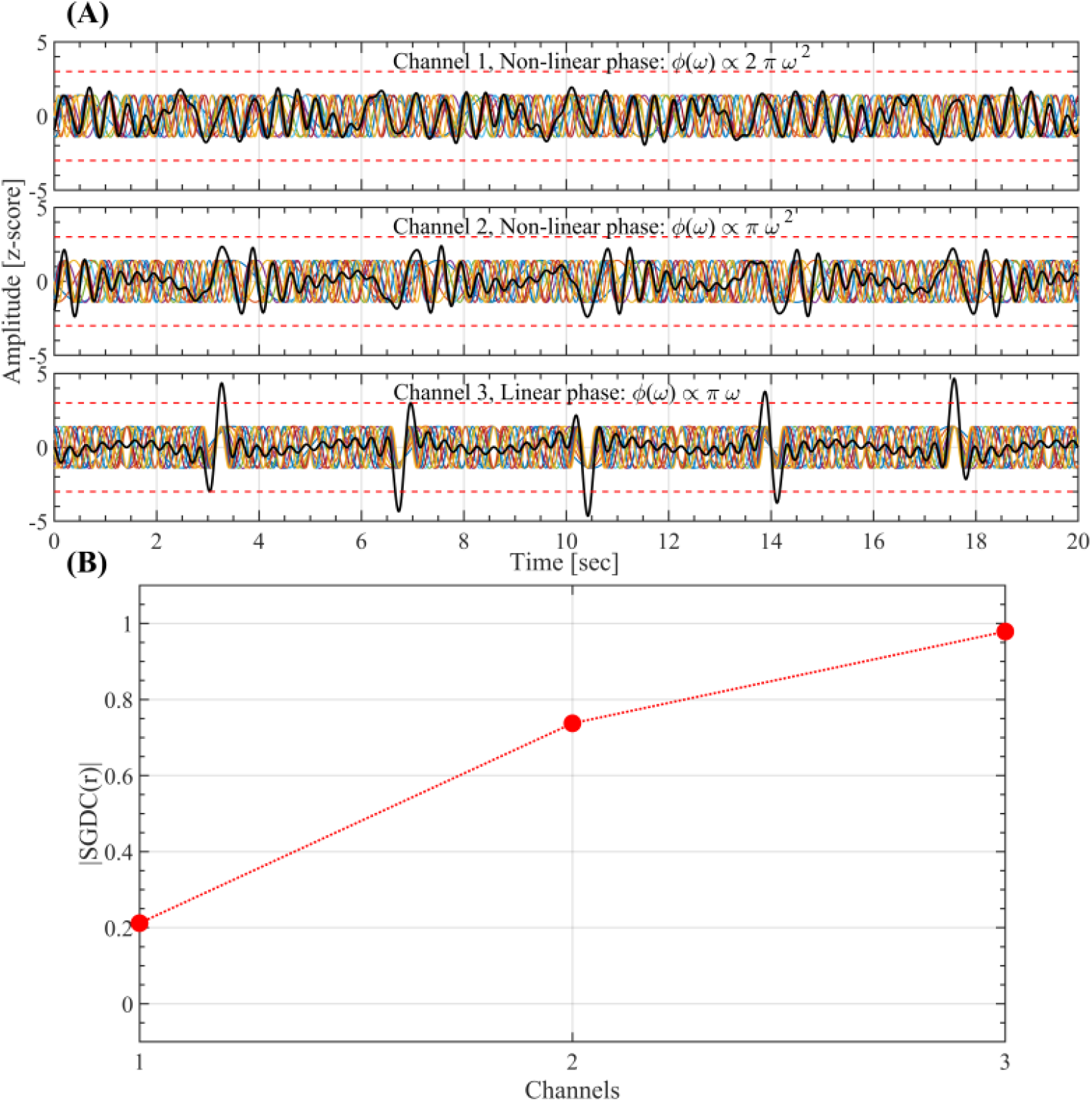
*SGDC*(*r*) computed using the Eq. A.18 on a multi-channel configuration. (A) Three time series *x*_*r*_ (*t*) (black solid lines) synthesized using the Eq. A.11. In each channel, the signal *x*_*r*_ (*t*) was synthesized by the linear superposition of 10 sinusoidal tones (colored solid lines) with unitary amplitude and uniformly spaced frequencies (*f*_*s*_ Δ*ω/*(2*π*) = 0.278 Hz) within the range *f*_*s*_ *ω*(*k*)*/*(2*π*) = 0.5 + *k f*_*s*_ Δ*ω/*(2*π*) ∈ [0.5 *−*3] Hz. In the channels 1 and 2, the phase of the tones were set as a quadratic function of the frequency within the range *ϕ*_1_(*ω*) ∝ 2*πω*^2^ ∈ [*−* 2*π*, 2*π*] and *ϕ*_2_(*ω*) ∝ *πω*^2^ ∈ [*−π, π*], respectively. In the channel 3, the phase of the tones were set as a linear function of the frequency within the range *ϕ*_3_(*ω*) ∝ *πω* ∈ [*−π, π*]. (B) Modulus of the *SGDC*(*r*) for each channel. Note that the higher the burstiness (i.e., amplitude of the transient fluctuations) disclosed by the resulting signal (see solid black line in the panel A), the higher the *SGDC*(*r*) value. As expected, the channel 3 corresponding to the tones having a linear phase dependence with the frequency discloses the maximum |*SGDC*(*r*)| ≈ 1.

Figs. A.5 and A.6 show the *SGDC*(*ω*) computed using the Eq. A.19 compared against the Phase Locking Value (PLV) assessed using the following expression [65, 32],

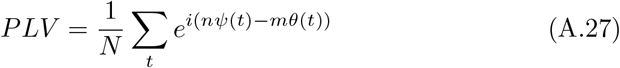

In Eq. A.27, *ψ*(*t*) and *θ*(*t*) are the phase time series of interest and the integers *n, m* ∈ ℕ are required to allow the comparison of phase time series pertaining to different frequency bands. Of note, the *SGDC*(*ω*) quantifies, at each frequency value, the bursts synchronization across the brain regions (channels), whereas the *PLV* quantifies either local or cross-regional phase coherence between two frequency bands and it is not related to the signal burstiness, i.e., the PLV is not sensitive to the emergence of above-threshold fluctuations neither to the cross-regional synchronization of salient events. Fig. A.5A shows three channels in which the resulting time series (solid black line) have been synthesized as the linear superposition of 10 sinusoidal tones with uniformly spaced frequencies (Δ*ω* = const) in the range *f*_*s*_ *ω/*(2*π*) ∈ [0.5 − 3] Hz (see Eq. A.11). In each channel, the phase of all the oscillatory components was set to zero (*ϕ*_*r*_(*ω*) = 0 ∀ *ω*). The local and cross-regional effects of this setup can be summarized as follows,

- In each channel (local effect), we obtain the maximum group delay consistency across frequency values accounting for the emergence of above-threshold fluctuations. That is, *ϕ*_*r*_(*ω*) = 0 =⇒ Δ*ϕ*_*r*_(*ω*) = 0 =⇒ *τ*_*r*_(*ω*) = −Δ*ϕ*_*r*_*/*Δ*ω* = 0 = cte =⇒ *SGDC*(*r*) = 1 : *r* = 1, 2, 3 (data not shown).
- At each frequency, we obtain the maximum group delay consistency across channels (cross-regional effect) accounting for the synchronization of the salient events across the channels. That is, *ϕ*_*r*_(*ω*) = 0 =⇒ Δ*ϕ*_*r*_(*ω*) = 0 =⇒ *τ*_*r*_(*ω*) = −Δ*ϕ*_*r*_*/*Δ*ω* = 0 = cte =⇒ *SGDC*(*ω*) = 1 ∀ *ω*. The resulting |*SGDC*(*ω*)| is shown in Fig. A.5B.
- At each frequency, we obtain the maximum phase coherence across channels (cross-regional effect). That is, *ψ*_*r,ω*_(*t*) − *θ*_*r*_*′* _,*ω*_ (*t*) = 0 =⇒ |*PLV* | = 1 ∀ *ω*, where the phase time series *ψ*_*r,ω*_(*t*) and *θ*_*r*_*′* _,*ω*_ (*t*) were extracted from different channels ((*r, r*) ∈ {1, 2, 3} : *r* ;= *r*) and evaluated at the same frequency *ω*. In other words, *ψ*_*r,ω*_(*t*) and *θ*_*r*_*′* _,*ω*_ (*t*) are the phase time series associated with two tones homologous in frequency and pertaining to different channels. The resulting |*PLV* | is shown in the Fig. A.5B.

Fig. A.5C shows three time series constituted by the same 10 tones used in Fig. A.5A, with the difference that in this case the phase of the tones were set as *ϕ*_1_(*ω*) = 0, *ϕ*_2_(*ω*) ∝ *−* 3*πω* and *ϕ*_3_(*ω*) ∝ +3*πω* for the channel 1, 2 and 3, respectively. The linear phase dependence with the frequency associated with the channels 2 and 3 produces a time-shift in the resulting signals. As a consequence, in this multi-channel configuration the resulting above-threshold fluctuations are not synchronized across channels (see the solid black lines in the Fig. A.5C). In this case, the SGDC and PLV measures result,

- In each channel (local effect), we obtain the maximum group delay consistency across frequency values accounting for the emergence of above-threshold fluctuations. That is, Δ*ϕ*_*r*_(*ω*) = const ⇒ *τ*_*r*_(*ω*) = *−*Δ*ϕ*_*r*_*/*Δ*ω* = cte ⇒ *SGDC*(*r*) = 1 : *r* = 1, 2, 3. Note that this result is similar to what we obtained for a constant group delay (i.e., not a function of the frequency) associated with the channel 3 shown in Fig. A.4.
- At each frequency, we obtain a low group delay consistency across channels (cross-regional effect) accounting for the lack synchronization of the salient events across the channels. That is, Δ*ϕ*_1_(*ω*) = 0, Δ*ϕ*_2_(*ω*) *<* 0, Δ*ϕ*_3_(*ω*) *>* 0 ⇒ *τ*_1_(*ω*) = 0, *τ*_2_(*ω*) *>* 0, *τ*_3_(*ω*) *<* 0 ⇒ *SGDC*(*ω*) ≈ 0 ∀ *ω*. The resulting |*SGDC*(*ω*)| is shown in the Fig. A.5D.
- At each frequency, we obtain the maximum phase coherence across channels (cross-regional effect). That is, *ψ*_*r,ω*_(*t*) − *θ*_*r*_*′* _,*ω*_ (*t*) = const =⇒ |*PLV* | = 1 ∀ *ω*, where the phase time series *ψ*_*r,ω*_(*t*) and *θ*_*r*_*′* _,*ω*_ (*t*) were extracted from different channels ((*r, r*) ∈ {1, 2, 3} : *r* ;= *r*) and evaluated at the same frequency *ω*. In other words, *ψ*_*r,ω*_(*t*) and *θ*_*r*_*′* _,*ω*_ (*t*) are the phase time series associated with two tones homologous in frequency and pertaining to different channels. The resulting |*PLV*| is shown in the Fig. A.5D.

It is essential to note that, the *SGDC*(*ω*) measure is highly sensitive to the cross-regional synchronization of the salient events, whereas the *PLV* measure is completely blind to this effect (compare Figs. A.5B and A.5D).

**Figure A.5:**
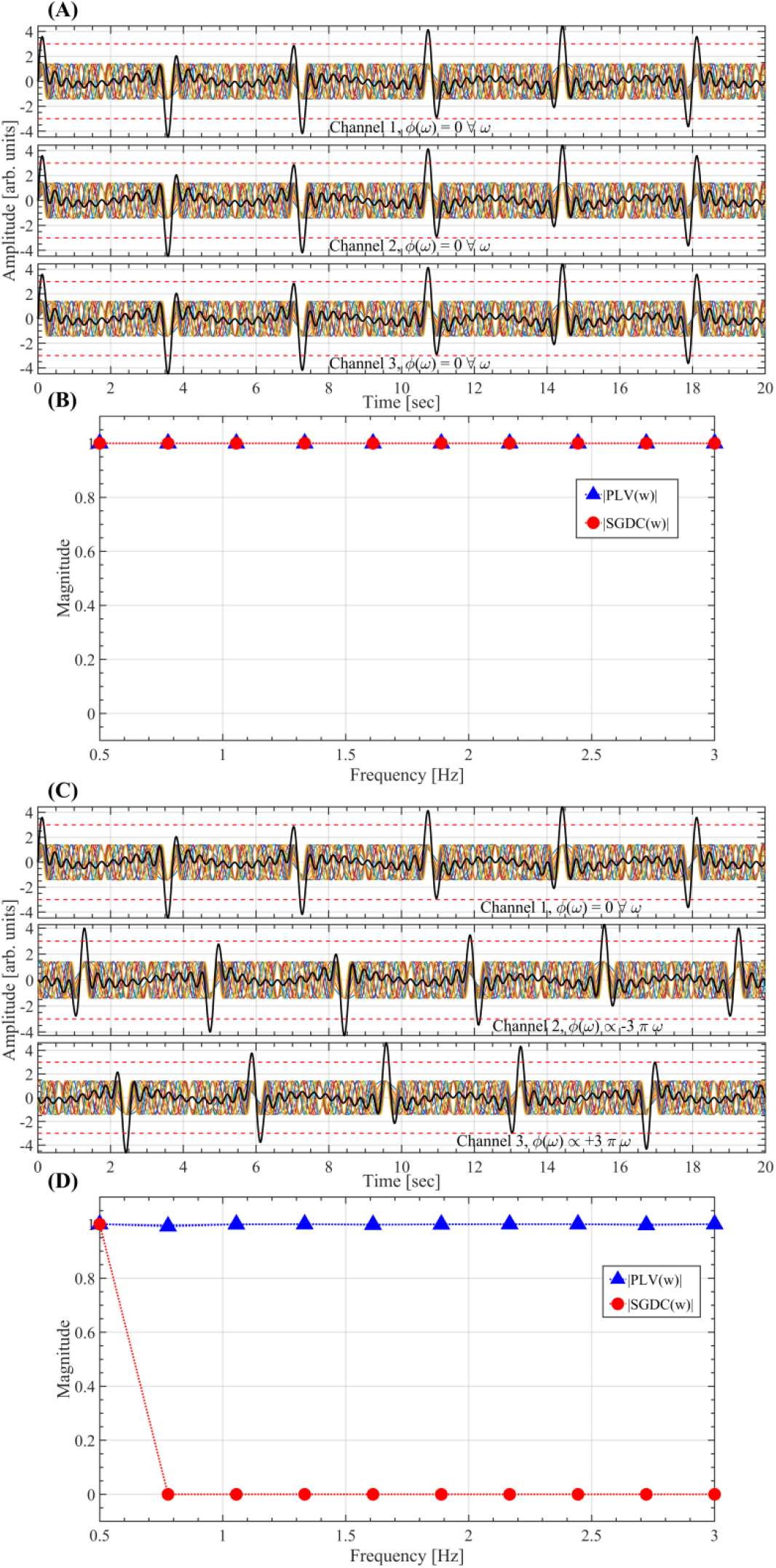
*SGDC*(*ω*) computed using the Eq. A.19 on a multi-channel configuration. (A) Three time series *x*_*r*_ (*t*) (black solid lines) synthesized using the Eq. A.11. In each channel, the signal *x*_*r*_ (*t*) was synthesized by the linear superposition of 10 sinusoidal tones (colored solid lines) with unitary amplitude and uniformly spaced frequencies (*f*_*s*_ Δ*ω/*(2*π*) = 0.278 Hz) within the range *f*_*s*_ *ω*(*k*)*/*(2*π*) = 0.5 + *k* 2*f*_*s*_0Δ*ω/*(2*π*) ∈ [0.5 − 3] Hz. In each channel, the phase of all the oscillatory components was set to zero (*ϕ*_*r*_ (*ω*) = 0 ∀ *ω*). (B) *SGDC*(*ω*) and *PLV* measures computed using the Eqs. A.19 and A.27, respectively, for the multi-channel configuration shown in panel A. (C) Same as in A, but in this case the phase of the tones were set as *ϕ*_1_(*ω*) = 0, *ϕ*_2_(*ω*) ∝ *−* 3*πω* and *ϕ*_3_(*ω*) ∝+3*πω* for the channel 1, 2 and 3, respectively. Same as in B for the multi-channel configuration shown in panel C.

Fig. A.6A shows three time series constituted by the same 10 tones used in Figs. A.5A and A.5C with the difference that in this case the phase of the tones were set as follows,

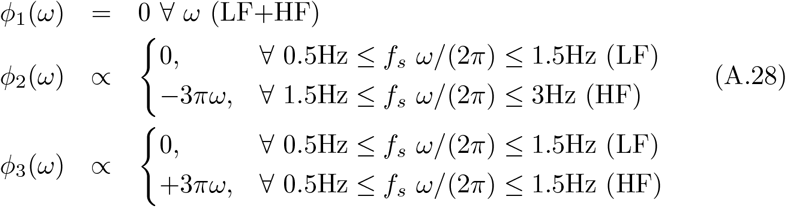

This phase configuration produce LF transient fluctuations co-occurring across the channels, while the resulting HF transient fluctuations are not synchronized across the channels (see Fig. A.6A). Importantly, the *SGDC*(*ω*) effectively discriminate the cross-regional synchronization of the transient fluctuations across the frequency values, whereas the *PLV* measure is again completely blind to this effect (see Fig. A.6B). Fig. A.6C shows three time series constituted by the same 10 tones used in Fig. A.6A (see Eq. A.11) with the difference that in this case the phase of the tones were set as follows,

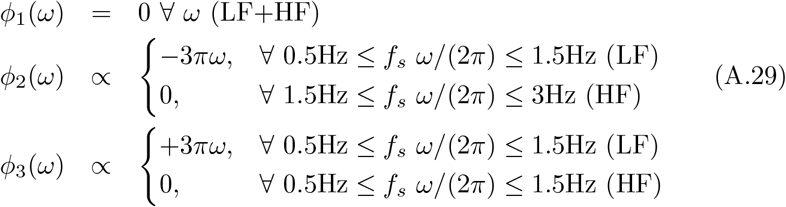

Similarly to the previous case, the *SGDC*(*ω*) effectively discriminate the cross-regional synchronization of the transient fluctuations across the frequency values, whereas the *PLV* measure is again completely blind to this effect (see Fig. A.6D). It is worth mentioning that Δ*ϕ*_*r*_(*ω*) in the Eq. A.19 is the incremental phase between adjacent spectral components associated with the activity *x*_*r*_(*t*) of the brain region *r*. Thus, for *N* spectral components we obtain *N* − 1 incremental phase values Δ*ϕ*_*r*_(*ω*). As a convention, we add an extra value Δ*ϕ*_*r*_(*ω*) = 0 as the first element (i.e., lowest frequency) of the list of incremental phase values. Hence, for *N* spectral components the Eqs. A.18 and A.19 produce *N* values of SGDC. In particular, the first value (i.e., lowest frequency) of *SGDC*(*ω*), associated with the artificially added Δ*ϕ*_*r*_(*ω*) = 0, is always equal to 1 (this becomes evident in the Figs. A.5D and A.6D).

**Figure A.6:**
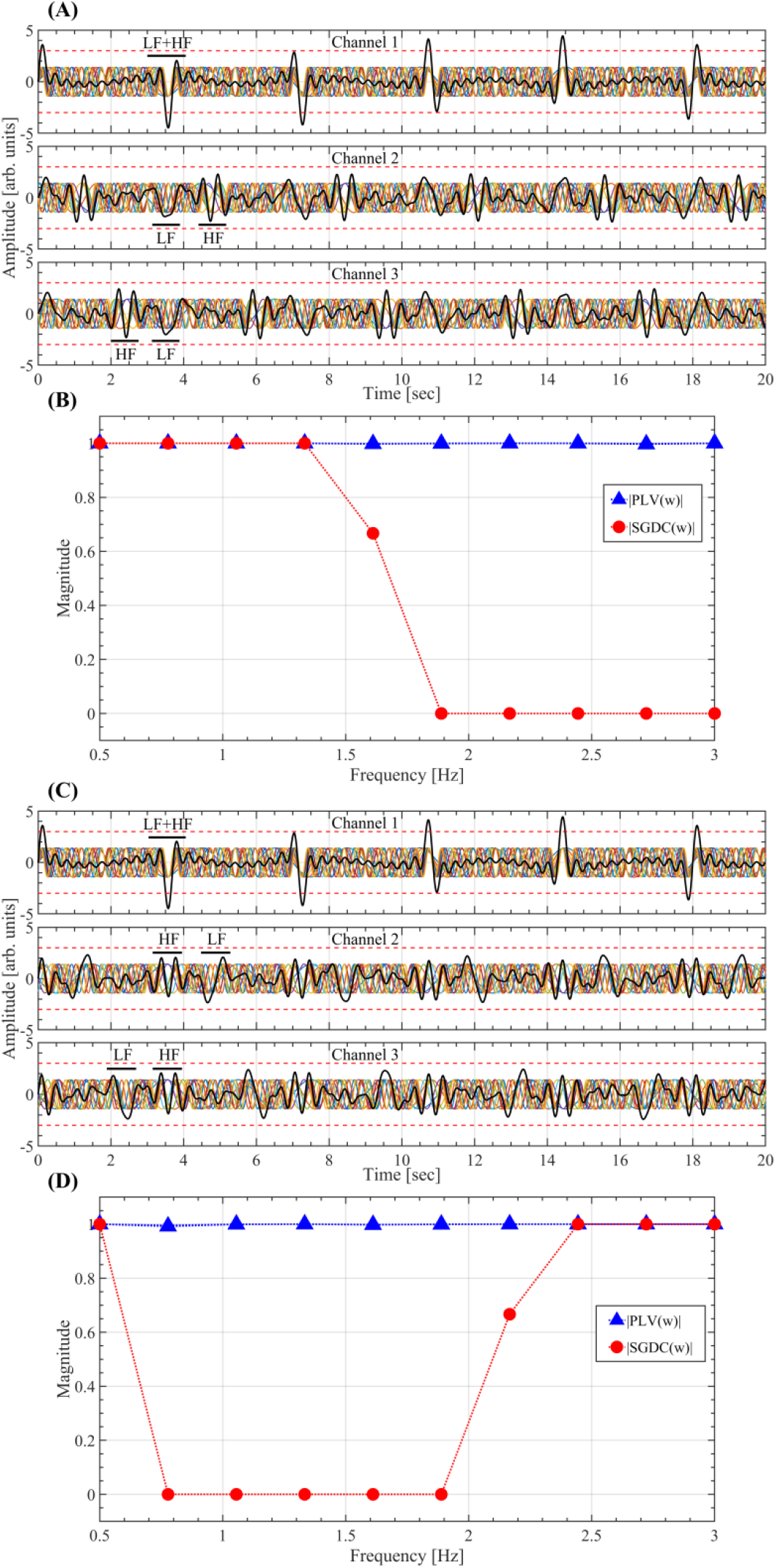
*SGDC*(*ω*) computed using the Eq. A.19 on a multi-channel configuration. (A) Three time series *x*_*r*_ (*t*) (black solid lines) synthesized using the Eq. A.11. In each channel, the signal *x*_*r*_ (*t*) was synthesized by the linear superposition of 10 sinusoidal tones (colored solid lines) with unitary amplitude and uniformly spaced frequencies (*f*_*s*_ Δ*ω/*(2*π*) = 0.278 Hz) within the range *f*_*s*_ *ω*(*k*)*/*(2*π*) = 0.5 + *k* 2*f*_*s*_2Δ*ω/*(2*π*) ∈ [0.5 *−* 3] Hz. In each channel, the phases of the oscillatory components were configured as stated in the set of Eqs. A.28. (B) *SGDC*(*ω*) and *PLV* measures computed using the Eqs. A.19 and A.27, respectively, for the multi-channel configuration shown in panel A. (C) Same as in A, but in this case the phase of the tones were configured as stated in the set of Eqs. A.29. (D) Same as in B for the multi-channel configuration shown in panel C.

### Appendix A.4. Spectral group delay consistency in the surrogate data

Here we analytically show that, on the one hand, A-surrogates significantly reduce the spectral group delay consistency (SGDC) across both frequency components (*SGDC*(*r*)) and brain regions (*SGDC*(*ω*)). On the other hand, B-surrogates significantly reduce the SGDC across frequency components (*SGDC*(*r*)), while preserving the SGDC across brain regions (*SGDC*(*ω*)).

We start by recalling the definition of *SGDC*(*r*) and *SGDC*(*ω*) for a multi-regional time series *x*_*r*_(*t*),

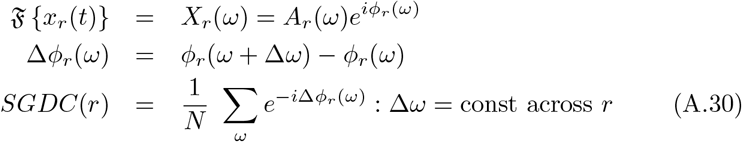

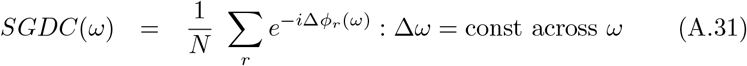

where *A*_*r*_(*ω*) and *ϕ*_*r*_(*ω*) are the amplitude and phase Fourier spectra, respectively. In Eqs. A.30 and A.31, *N* is the number of either frequency values or brain regions, respectively, and Δ*ϕ*_*r*_(*ω*) is the incremental phase computed across the spectral components of the DFT spectrum *X*_*r*_(*ω*) associated with the signals *x*_*r*_(*t*). In the case of the surrogate multi-regional time series 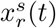, obtained by phase randomization of the original time series in the frequency-domain, we have,

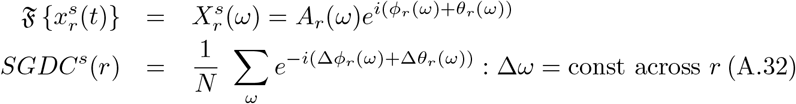

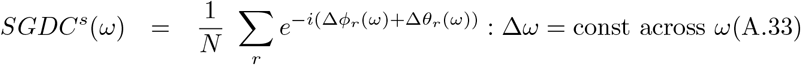

In the Eqs. A.32 and A.33, Δ*θ*_*r*_(*ω*) is the incremental phase associated with the random phase-shift *θ*_*r*_(*ω*) extracted from the surrogate DFT spectrum 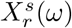 of each brain region *r*. Let us consider two extreme cases derived from the Eqs. A.30 and A.32 with Δ*θ*_*r*_(*ω*) varying randomly across *ω*,

1. For Δ*ϕ*_*r*_(*ω*) ≈ *const* ⇒ |*SGDC*(*r*)| ≈ 1 *>* |*SGDC*^*s*^(*r*)| ≈ 0.
2. For Δ*ϕ*_*r*_(*ω*) varying randomly across *ω* ⇒ |*SGDC*(*r*)| ≈ |*SGDC*^*s*^(*r*)| ≈ 0.

From these two extreme cases we infer that, for *θ*_*r*_(*ω*) varying randomly across *ω*, |*SGDC*(*r*)| is the upper bound of |*SGDC*^*s*^(*r*)|. As a consequence, for the A- and B-surrogates in general we obtain |*SGDC*^*s*^(*r*)| *<* |*SGDC*(*r*)|. Similarly, in the case of A-surrogates computed with *θ*_*r*_(*ω*) varying randomly across the brain regions *r*, Eqs. A.31 and A.33 in general produce |*SGDC*^*s*^(*ω*)| *<* |*SGDC*(*ω*) |. In the particular case of the B-surrogates, at each frequency *ω* we add the same phase-shift value *θ*_*r*_(*ω*) in all the brain regions *r*, producing Δ*θ*_*r*_(*ω*) = Δ*θ*(*ω*) ∀ 1 ≤ *r* ≤ *N* . As a consequence, by taking the modulus in both sides of the Eq. A.33 we obtain the equivalence between the true data and the B-surrogate in terms of |*SGDC*(*ω*)|,

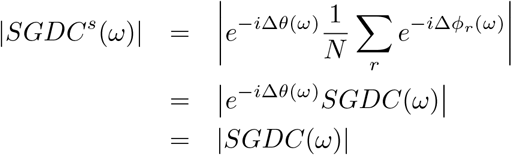

We confirmed this analytical results by computing the *SGDC*(*r*) and *SGDC*(*ω*) measures on the whole time series of our empirical MEG dataset and the corresponding A- and B-surrogates (see Section 2.8 in Methods). Fig. A.7A shows that the magnitude of the *SGDC*(*r*) measure is not preserved in both the A- and B-surrogates. Besides, Fig. A.7B shows that the magnitude of the *SGDC*(*ω*) measure is preserved in the B-surrogates, and not in the case of A-surrogates. Importantly, the reduction of the regional SGDC, as quantified by the *SGDC*(*r*) measure, offers an analytical rationale supporting the evidence showing that B-surrogates failed to reproduce the SEs observed in our MEG dataset (see Section 3.2) despite preserving both the regional PSDs and the cross-spectra (see Ap- pendix A.1). It is important to note that this equivalence between the true MEG data and the B-surrogates in terms of |*SGDC*(*ω*)| holds only when the *SGDC*(*ω*) measure is computed on the whole time series (i.e., non-time-resolved approach). On the other hand, if the *SGDC*(*r*) and *SGDC*(*ω*) measures are computed in a time-resolved manner on each salient event (see Fig. A.7C,D), the equivalence between the true MEG data and the B-surrogates in terms of |*SGDC*(*ω*)| does not longer hold. This is mainly due to the fact that true SEs and B-surrogate SEs are different in duration and size and, more crucially, they do not necessarily involve the same brain regions.

**Figure A.7:**
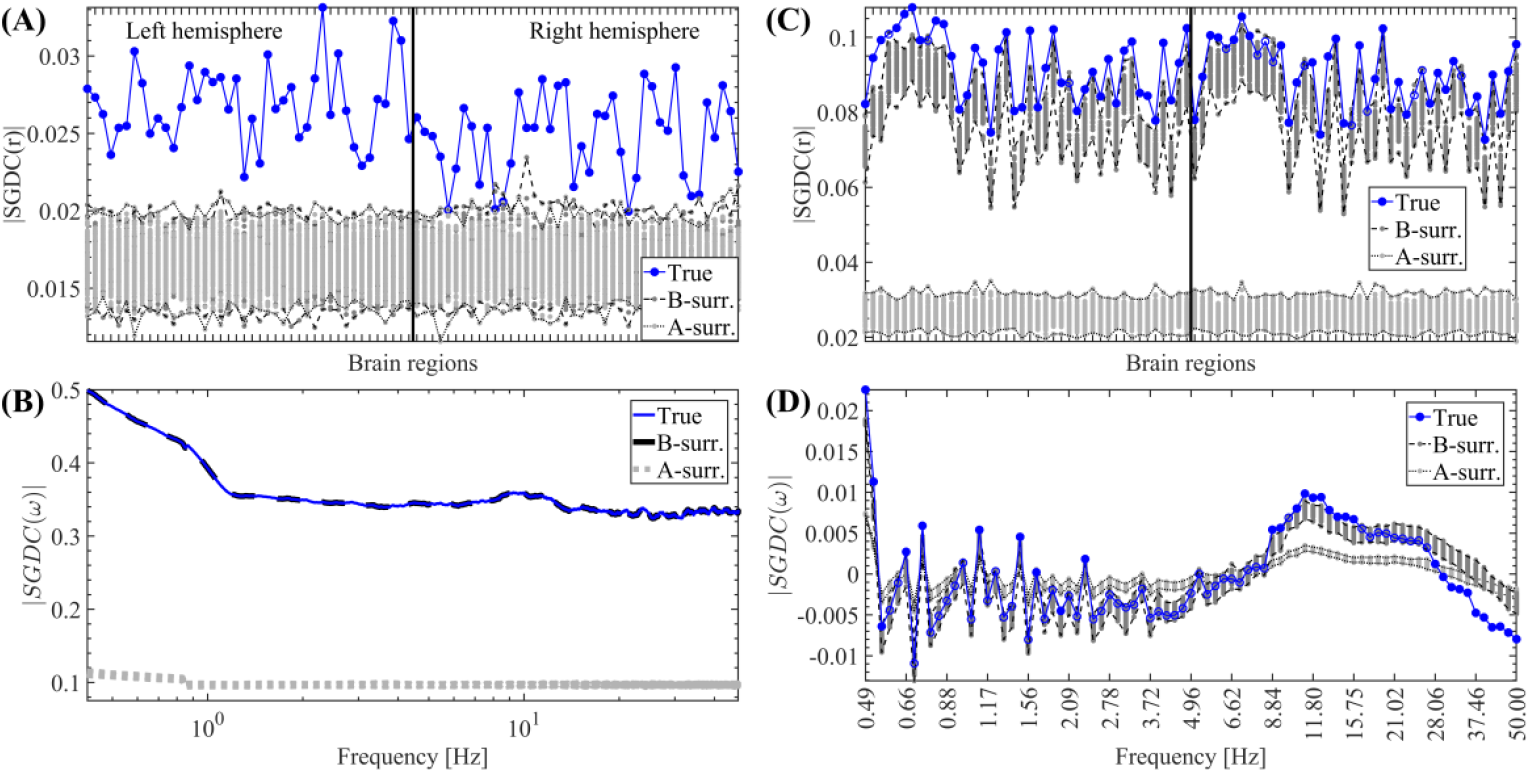
Spatial profiles associated with the SGDC measures. (A) *SGDC*(*r*) measure computed on the whole time series of each brain region (i.e., non-time-resolved approach). (B) *SGDC*(*ω*) measure computed on the whole time series of each brain region (i.e., non-time-resolved approach). Note that the pattern corresponding to the 100 B-surrogates (thick dashed black line) overlap with the spatial profile associated with the true MEG data (thin blue line). (C) *SGDC*(*r*) measure computed on each detected SE by considering the brain regions and time interval associated with each particular event (i.e., time-resolved approach). (D) *SGDC*(*ω*) measure computed on each detected SE by considering the brain regions and time interval associated with each particular event (i.e., time-resolved approach). The labels and ordering of the brain regions are the same as those shown in Fig. C.2. Symbols and abbreviations: SE, Salient Event.

## Appendix B. Supplementary numerical modeling results

**Figure B.1:**
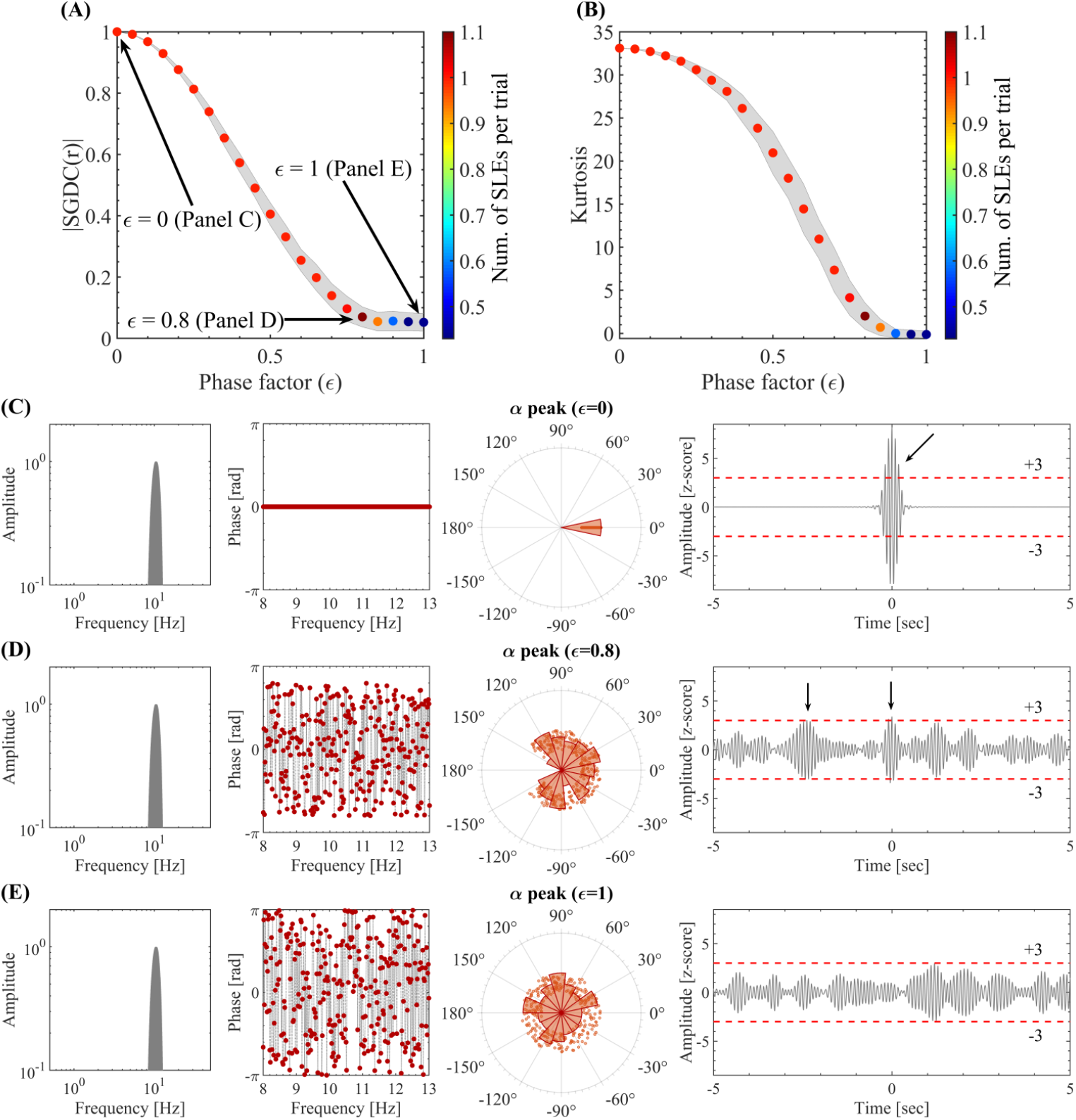
Spectral group delay consistency underlies the emergence of local above-threshold fluctuations from NOs. (A) Spectral group delay consistency, as quantified by the *SGDC*(*r*) measure, as a function of the phase factor values (*ϵ*). The colored markers indicate the mean |*SGDC*(*r*)| value across 100 synthetic time series of 10 sec in duration (trials). The shaded error bars in gray correspond to the standard deviation around the mean value. The pseudocolor scale represents the mean number of SLEs per trial. The *SGDC*(*r*) measure was obtained by computing the Eq. 1 on the synthetic phase values assigned to the spectral components in the alpha band. (B) Same as in A for the Kurtosis of the time series amplitude values, obtained by computing the Eq. A.25 on the signals in time-domain. (C) Amplitude spectrum (left), phase spectrum and distribution (middle), and resulting time series (right) corresponding to the signal model for a phase factor *ϵ* = 0. For the amplitude spectrum we used a Hann window with a null-to-null bandwidth = 8-13 Hz, frequency resolution *df* = 1*/*60sec ≈ 0.017 Hz. The phase values of the spectral components were constrained within the range [*−ϵπ, ϵπ*] and having a random dependence with the frequency. The black arrows in the right-most panel highlight the above-threshold fluctuations disclosed by the signal. (D) Same as in C for a phase factor *ϵ* = 0.8. (E) Same as in C for a phase factor *ϵ* = 1. Symbols and abbreviations: SLEs, Salient Local Events; SGDC, Spectral Group Delay Consistency.

**Figure B.2:**
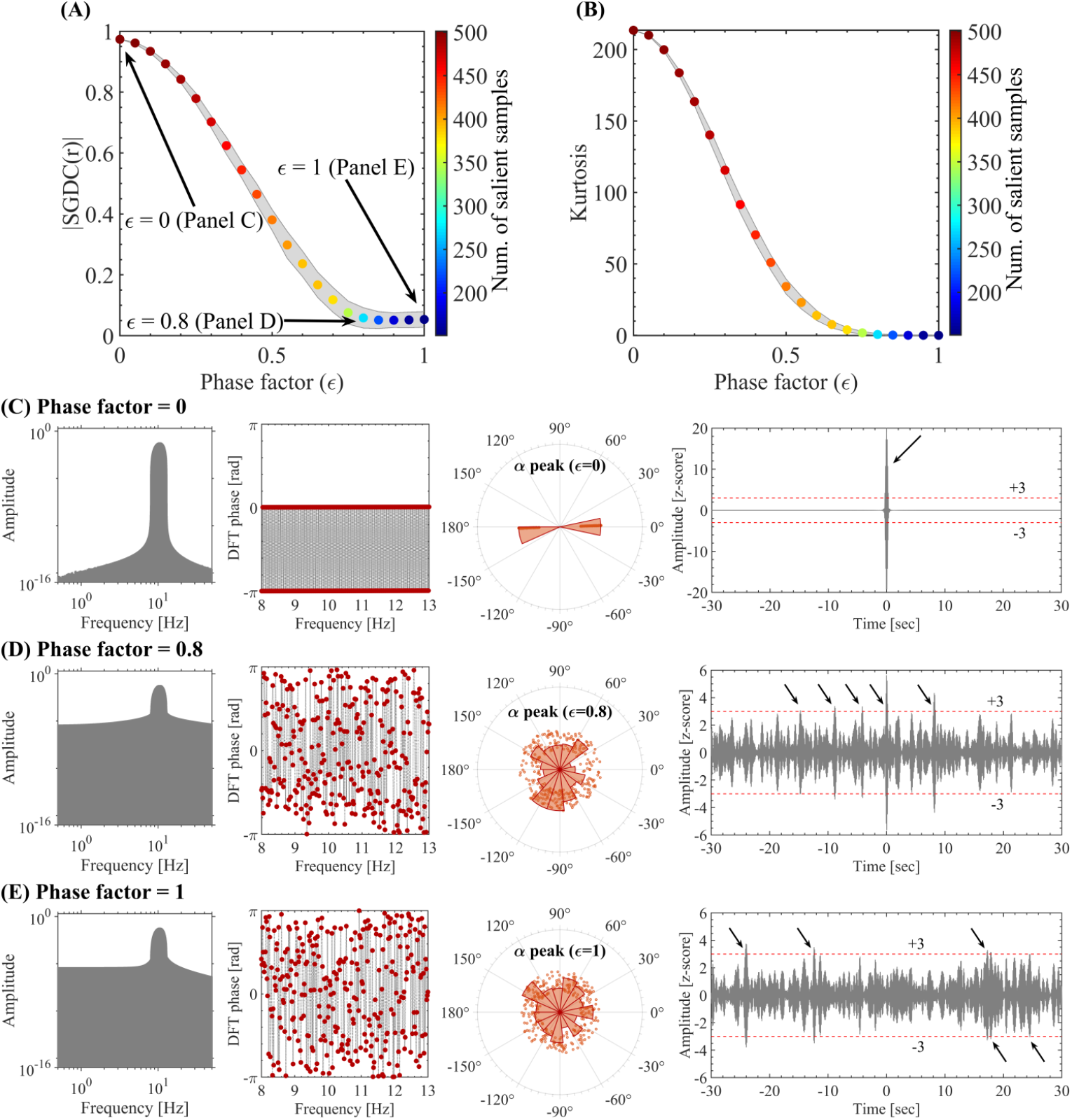
Spectral group delay consistency underlies the emergence of local above-threshold fluctuations from NOs. (A) Spectral group delay consistency, as quantified by the *SGDC*(*r*) measure, as a function of the phase factor values (*ϵ*). The colored markers indicate the mean |*SGDC*(*r*)| value across 100 synthetic time series of 60 sec in duration (trials). The shaded error bars in gray correspond to the standard deviation around the mean value. The pseudocolor scale represents the mean number of SLEs per trial. The *SGDC*(*r*) (Eq. 1) was computed using the alpha band phases obtained from the DFT applied to the time series resulting from the signal model (e.g., see the 60 sec in duration signals shown in panels C, D and E). This procedure inherently introduces spectral leakage due to the time-domain tapering (rectangular window), which affects the alpha band phase values involved in the computation of the *SGDC*(*r*) measure and is visible in the corresponding power spectra shown in panels C, D and E. (B) Same as in A for the Kurtosis of the time series amplitude values, obtained by computing the Eq. A.25 on the signals in time-domain. (C) Amplitude spectrum (left), phase spectrum and distribution (middle), and resulting time series (right) corresponding to the signal model for a phase factor *ϵ* = 0. The phase values of the spectral components were constrained within the range [*−ϵπ, ϵπ*] and having a random dependence with the frequency. The black arrows in the right-most panel highlight the above-threshold fluctuations disclosed by the signal. (D) Same as in C for a phase factor *ϵ* = 0.8. (E) Same as in C for a phase factor *ϵ* = 1. Symbols and abbreviations: SLEs, Salient Local Events; SGDC, Spectral Group Delay Consistency.

**Figure B.3:**
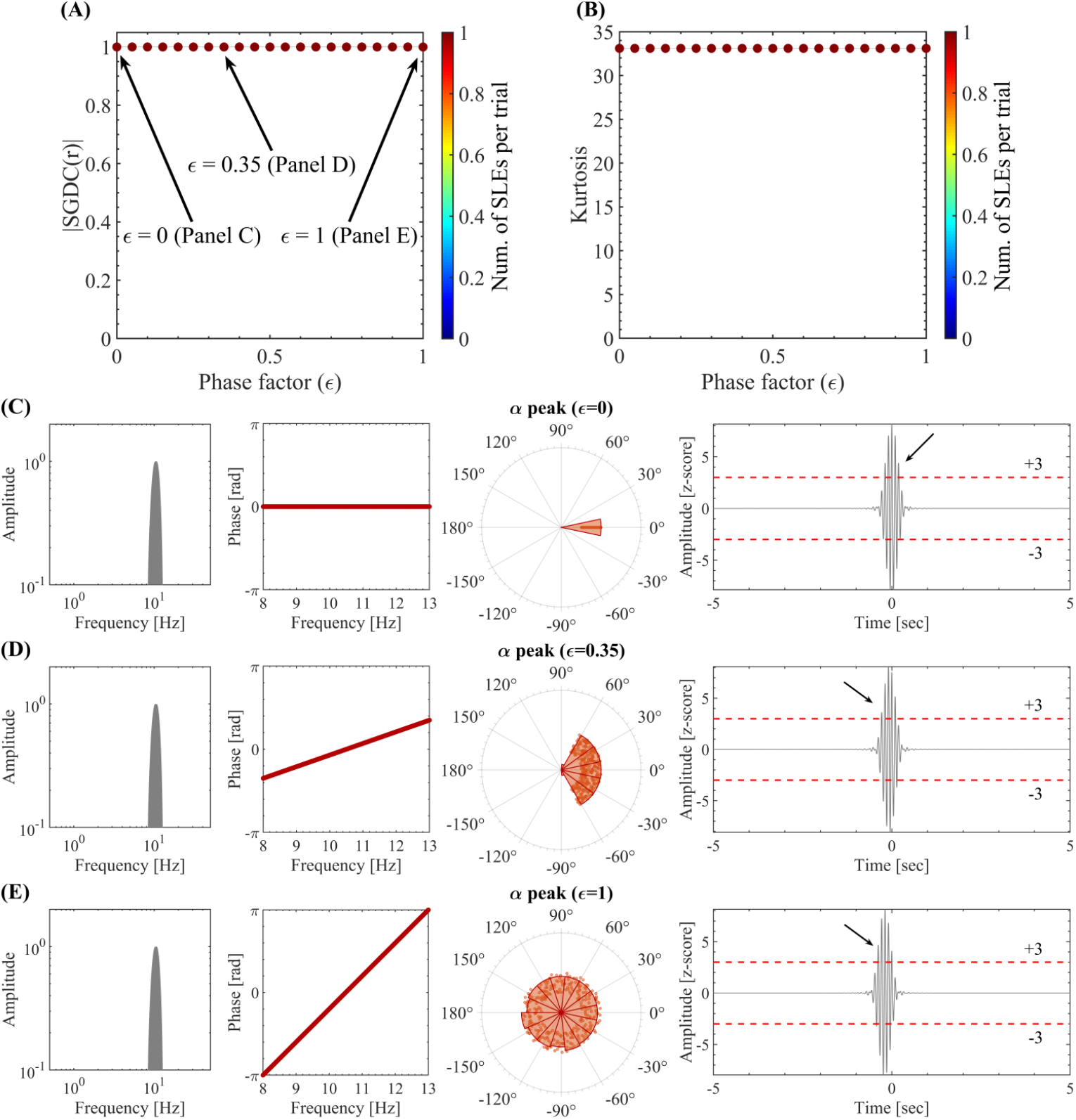
Spectral group delay consistency underlies the emergence of local above-threshold fluctuations from NOs. (A) Spectral group delay consistency, as quantified by the *SGDC*(*r*) measure, as a function of the phase factor values (*ϵ*). The colored markers indicate the mean |*SGDC*(*r*)| value across 100 synthetic time series of 10 sec in duration (trials). The shaded error bars in gray correspond to the standard deviation around the mean value. The pseudocolor scale represents the mean number of SLEs per trial. The *SGDC*(*r*) measure was obtained by computing the Eq. 1 on the synthetic phase values assigned to the spectral components in the alpha band. (B) Same as in A for the Kurtosis of the time series amplitude values, obtained by computing the Eq. A.25 on the signals in time-domain. (C) Amplitude spectrum (left), phase spectrum and distribution (middle), and resulting time series (right) corresponding to the signal model for a phase factor *ϵ* = 0. For the amplitude spectrum we used a Hann window with a null-to-null bandwidth = 8-13 Hz, frequency resolution *df* = 1*/*60sec ≈ 0.017 Hz. The phase values of the spectral components were constrained within the range [*−ϵπ, ϵπ*] and having a linear dependence with the frequency. The black arrows in the right-most panel highlight the above-threshold fluctuations disclosed by the signal. (D) Same as in C for a phase factor *ϵ* = 0.8. (E) Same as in C for a phase factor *ϵ* = 1. Symbols and abbreviations: SLEs, Salient Local Events; SGDC, Spectral Group Delay Consistency.

**Figure B.4:**
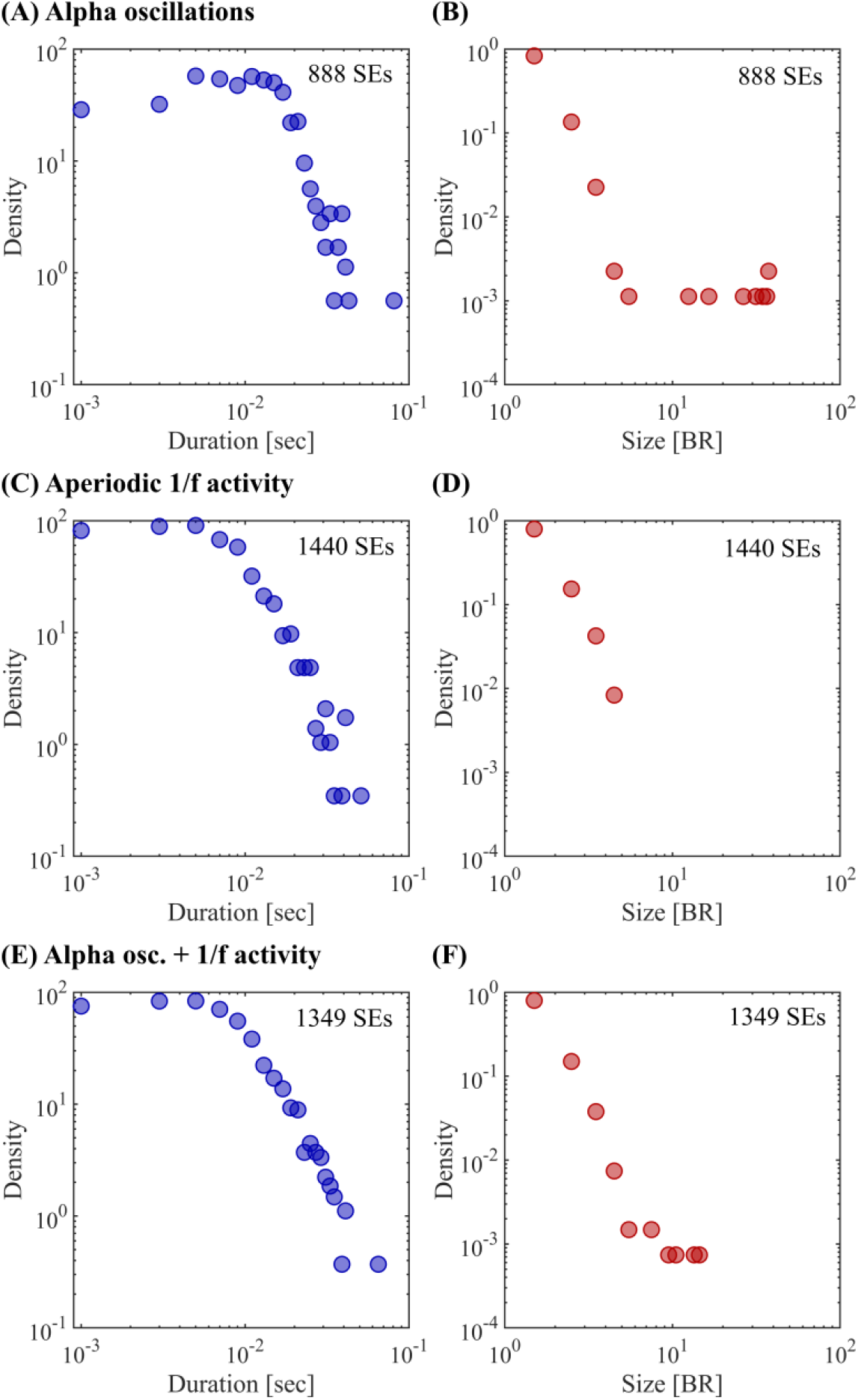
Distributions of size and duration corresponding to the SEs detected in the large-scale signal model. (A-B) Large-scale model for SEs including only alpha oscillations (random phase values in the alpha band constrained to the range [*− ϵπ, ϵπ*] with *ϵ* ∈ [0.75, 1]). Panels A and B show the distribution of SEs duration and size, respectively, computed on all the SEs detected in a simulated time series of 1-minute duration. See Figs. 6A,B. (C-D) Same as in A-B for the large-scale model including only broadband 1*/f* activity, and no oscillatory activity in the alpha band nor phase consistency values were present (*ϵ* = 1). See Figs. 6C,D. (E-F) Same as in A-B for the large-scale model including both broadband 1*/f* activity with non-constrained random phases (*ϵ* = 1) and alpha oscillations with random phases constrained proportionally to the observed alpha power in the range (*ϵ* ∈ [0.75, 1]). See Figs. 6E,F. Symbols and abbreviations: SEs, Salient Events.

## Appendix C. Supplementary empirical results including the deep sources

**Figure C.1:**
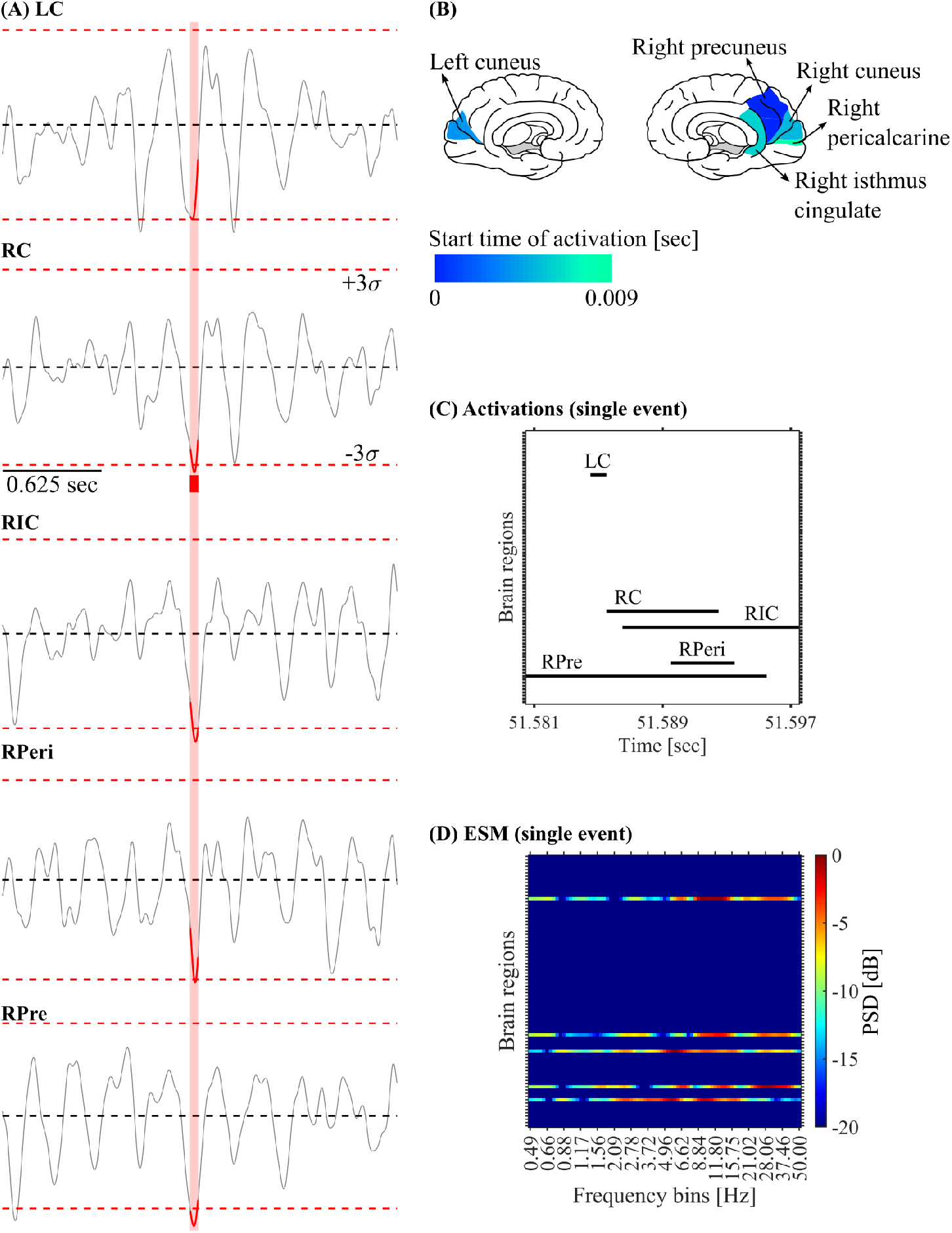
Salient Network Event (SNE). (A) Z-scored time series disclosing the above-threshold fluctuations associated with a SNE observed in the source-reconstructed MEG data. The time interval in which at least one brain region is active (i.e., duration of the SNE) is highlighted in red. (B) Brain plots showing the activation start time of the 5 brain regions recruited by the SNE shown in panel A. (C) Activation matrix of the SNE shown in panel A. The black segments correspond to the time intervals in which each brain region was active (i.e., absolute amplitude *>* 3*σ*). (D) ESM corresponding to the SNE shown in panel A. Symbols and abbreviations: ESM, Event Spectral Matrix; MEG, Magnetoencephalography; RPre, Right Precuneus; RC, Right Cuneus; RPeri, Right Pericalcarine; RIC, Right Isthmus Cingulate; LC, Left Cuneus.

**Figure C.2:**
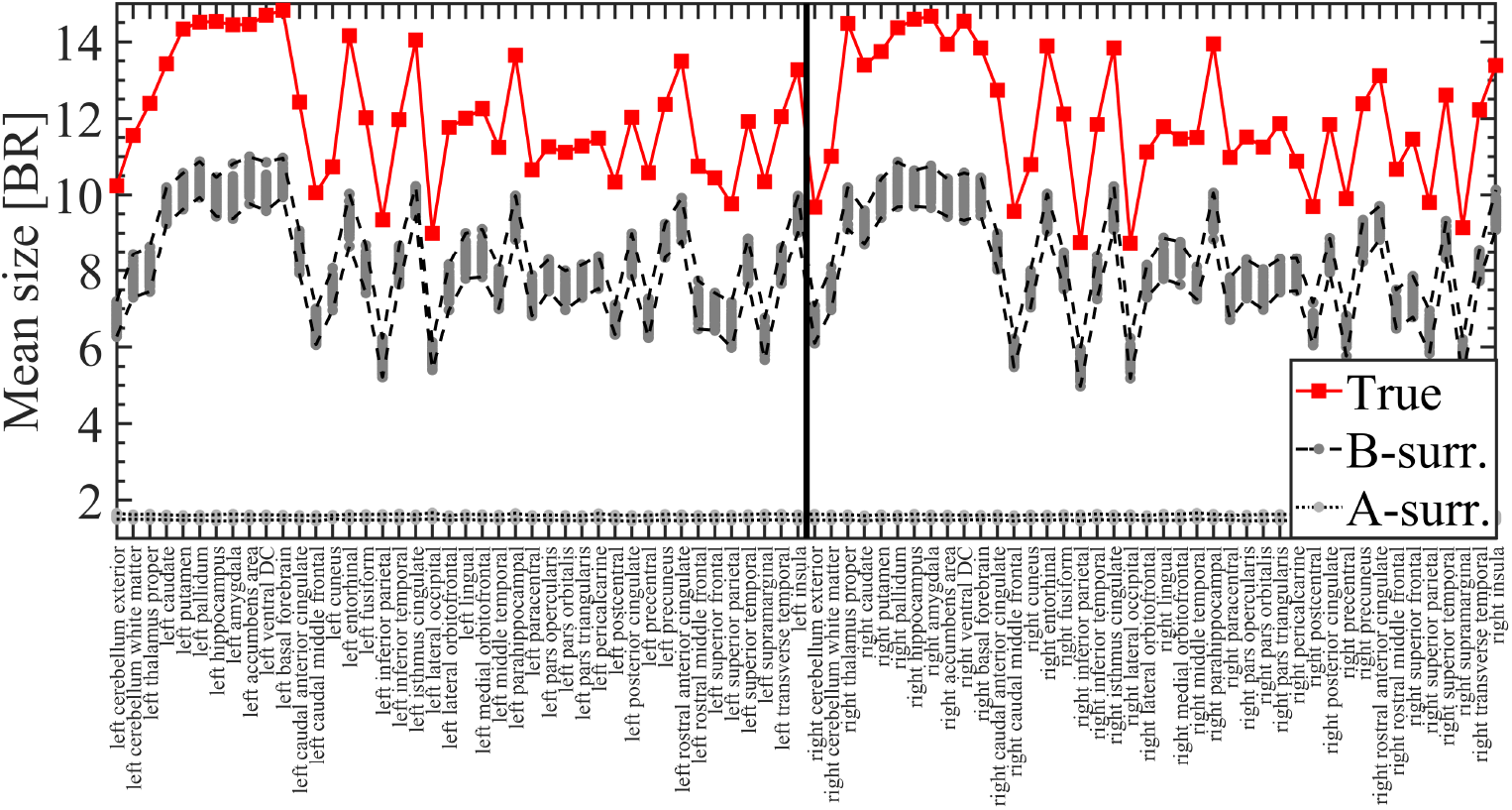
Labels and ordering of the brain regions used to compute all the spatial profiles shown in this work. Spatial profile showing the mean size of SEs propagating through each brain region (mean value across the 47 participants, see Section 2.4 in Methods). The mean event size is shown for the MEG data together with the 100 A- and B-surrogates (see Section 2.8 in Methods). Symbols and abbreviations: SEs, Salient Events; BR, Brain Regions.

**Figure C.3:**
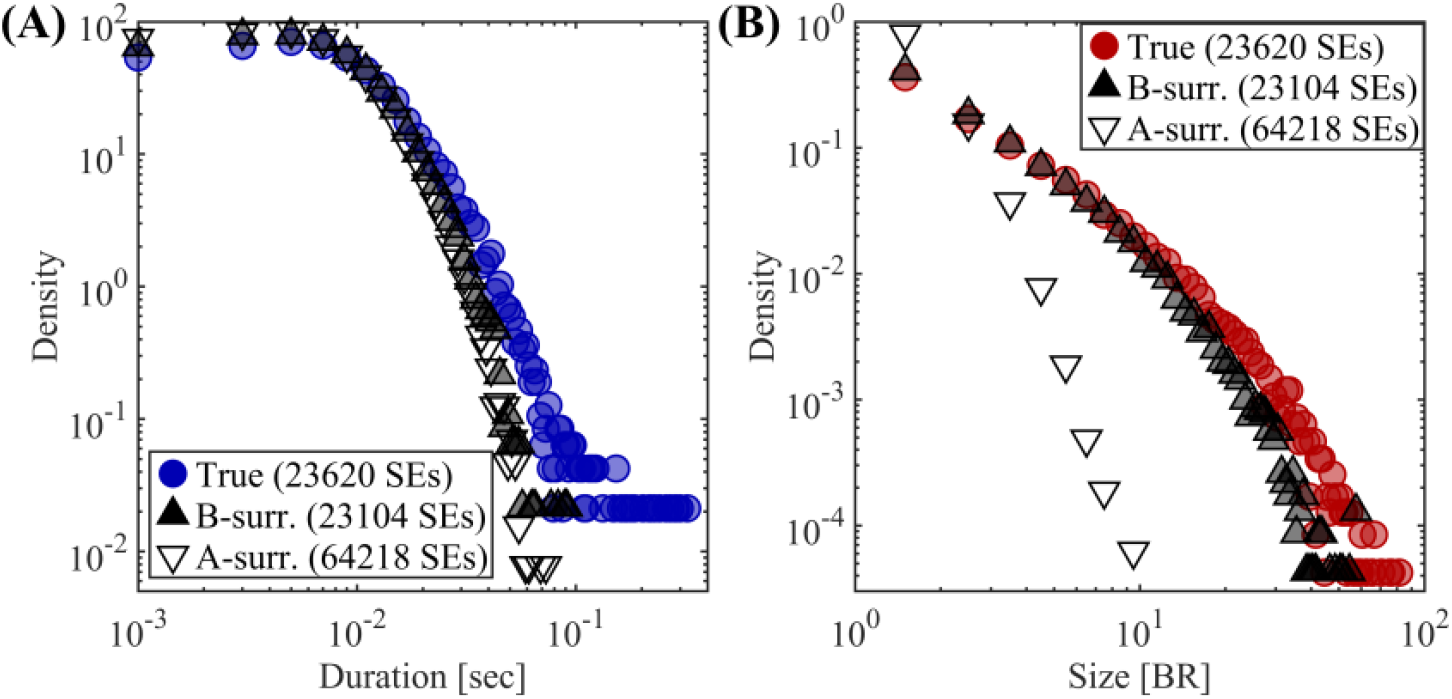
Statistical characterization of SEs. (A) Distribution of the duration of SEs observed in the true source-reconstructed MEG data (filled blue circles), the A-surrogate (empty down-pointing triangles) and the B-surrogate (filled up-pointing triangles) corresponding to a time binning of 1 time sample per time bin (time binning = 1 ms). In the three cases the SEs were computed on the 47 participants. (B) Same as in A for the size of SEs. To test the significance of the difference of the distribution means between the true MEG data and the surrogates (A and B), we computed a non-parametric permutation test (random sampling without replacement, 1 *×* 10^4^ permutations). The distributions of the duration and size of SEs observed in the true source-reconstructed MEG data, disclosed statistically significant differences with respect to both A- and B-surrogates (*P <* 0.001). Symbols and abbreviations: SEs, Salient Events; BR, Brain Regions.

**Figure C.4:**
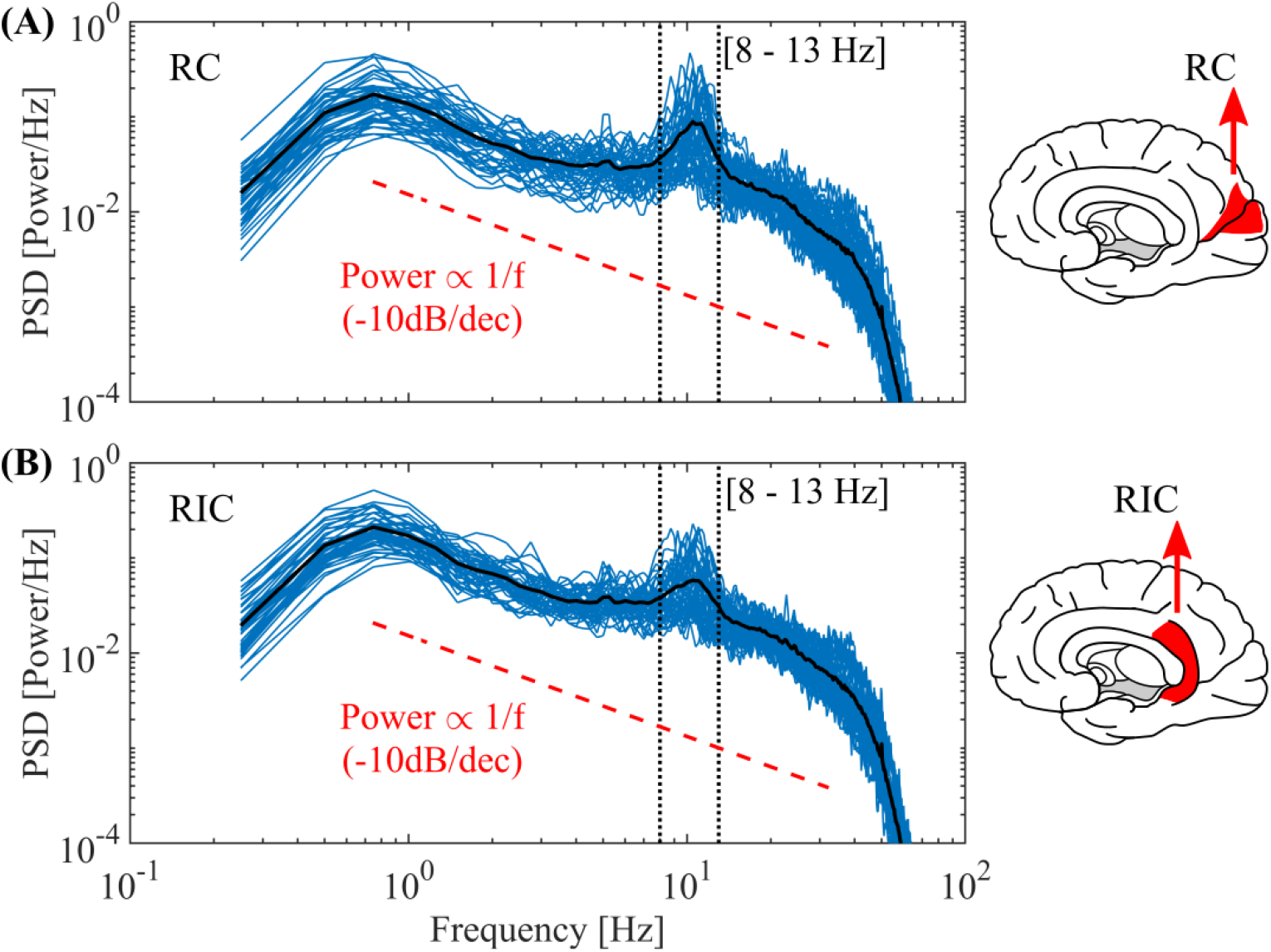
Power Spectral Density (PSD). Power spectra computed on the Right Cuneus (RC, panel A) and the Right Isthmus Cingulate (RIC, panel B) activities of each patient (blue lines) and the resulting average (black line). The PSDs were computed on 1 min duration source-reconstructed MEG data of 47 subjects. Note that the PSDs of the RC (panel A) disclose a prominent bump in the alpha band (8-13 Hz) characteristic of the occipital brain regions, however, a less prominent bump in the alpha band is also observed in regions away from the occipital cortex (see the PSDs of RIC shown in panel B). Symbols and abbreviations: PSD, Power Spectral Density; RC, Right Cuneus; RIC, Right Isthmus Cingulate.

**Figure C.5:**
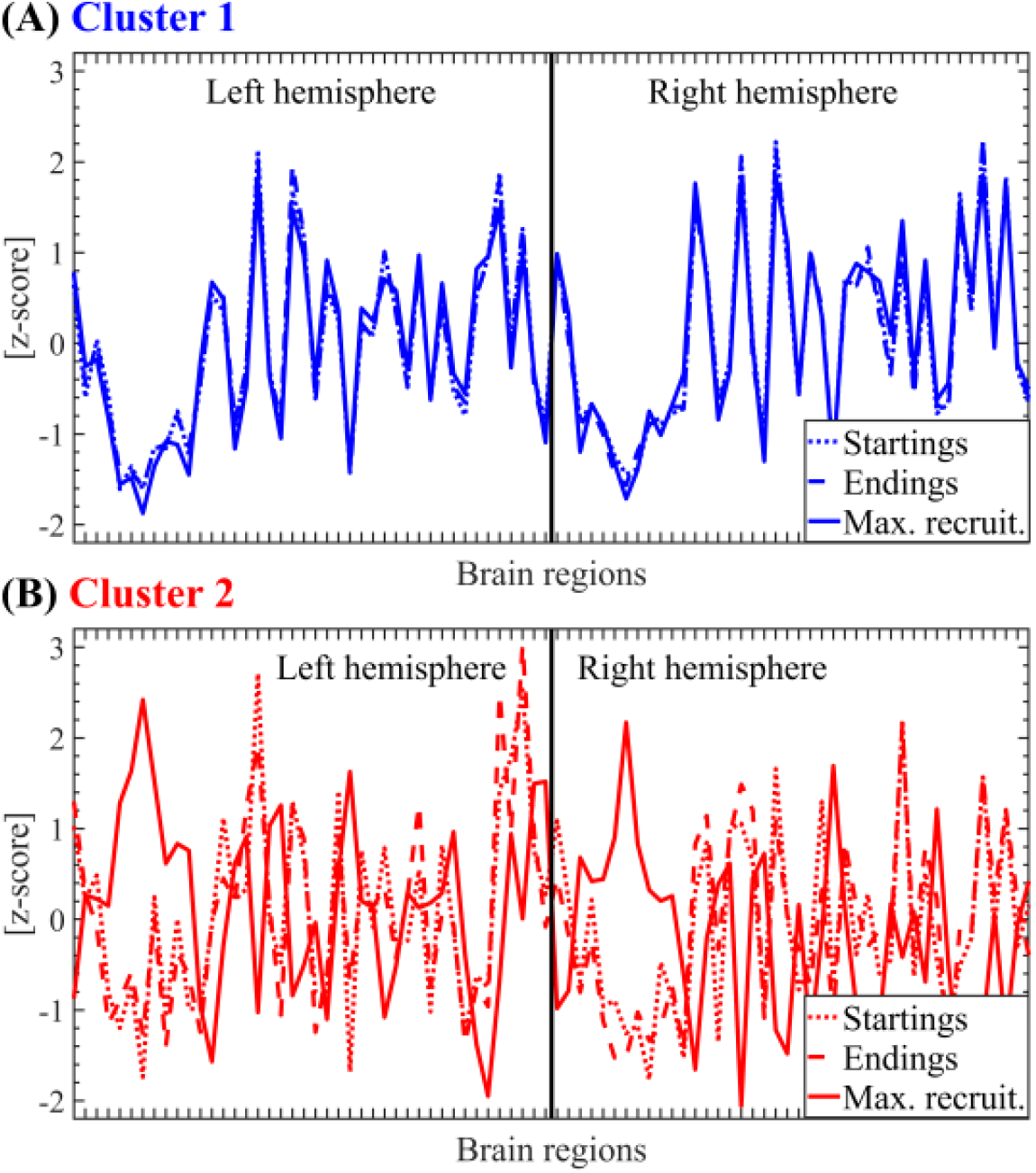
Salient events propagation modes segregated by SE clusters. (A) Spatial profile for the cluster 1 SEs starting, maximum recruitment and ending modes (see Section 2.7 in Methods) computed on 41 participants. Linear correlations between topographies: Startings vs Endings, *r* = 0.995, *P <* 0.001. Max. recruit. vs Startings, *r* = 0.978, *P <* 0.001. Max. recruit. vs Endings, *r* = 0.978, *P <* 0.001. (B) Same as in A for the cluster 2 SEs starting, maximum recruitment and ending modes. Linear correlations between topographies: Startings vs Endings, *r* = 0.895, *P <* 0.001. Max. recruit. vs Startings, *r* = *−*0.298, *P <* 0.01. Max. recruit. vs Endings, *r* = *−*0.280, *P <* 0.01. The SEs obtained from 41 subjects were clustered using the Louvain algorithm (resolution parameter *γ* = 1, see Section 2.9 in Methods). The reported P values for the statistical significance of the Pearson’s correlation were assessed using Student’s t distributions of the two-tailed hypothesis test under the null hypothesis that the correlation is zero. Symbols and abbreviations: SEs, Salient Events.

**Figure C.6:**
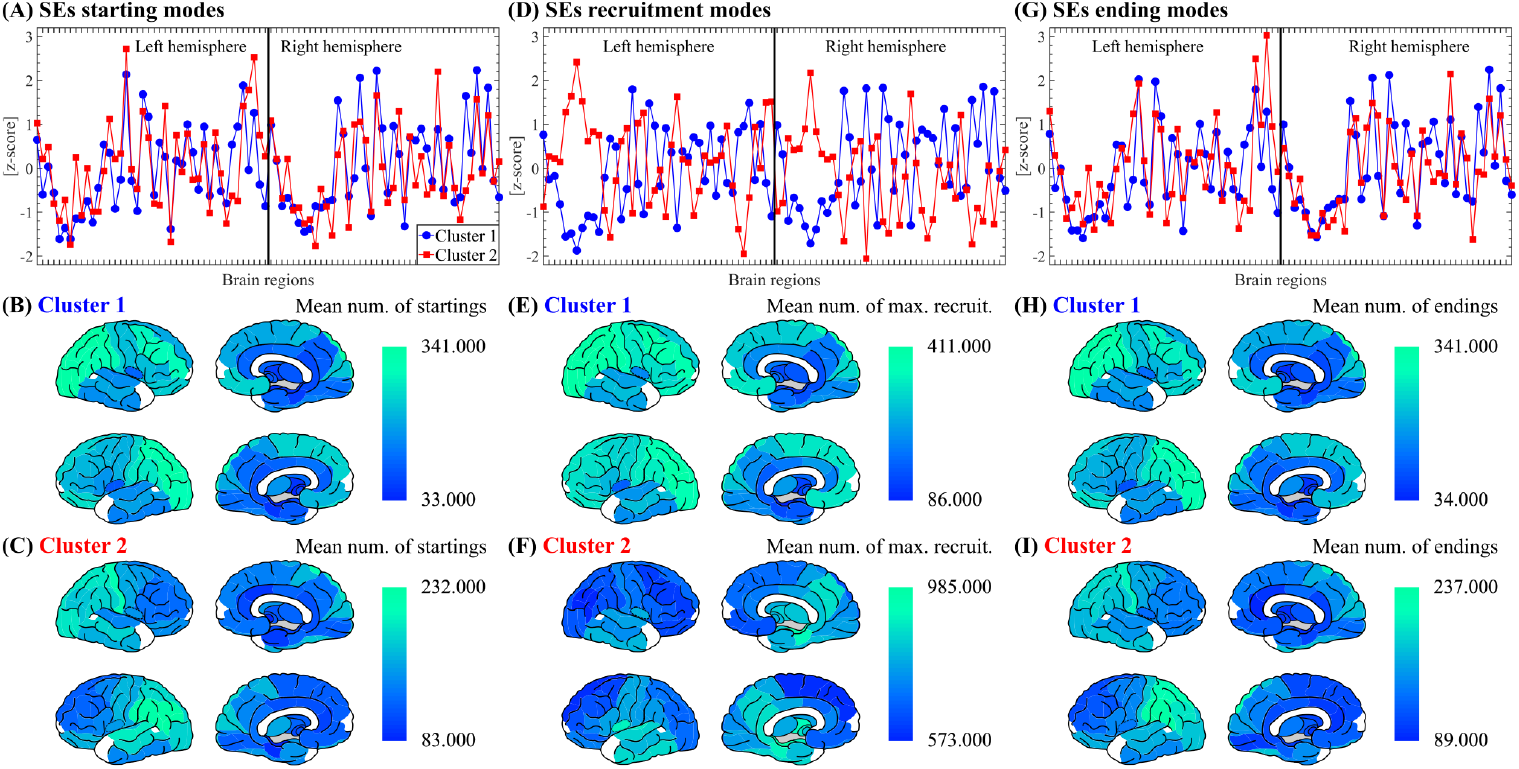
Salient events propagation modes. (A) Spatial profile for the SEs starting modes (see Section 2.7 in Methods) corresponding to the two SE clusters computed on 41 participants. The SEs obtained from 41 subjects were clustered using the Louvain algorithm (resolution parameter *γ* = 1, see Section 2.9 in Methods). The Pearson’s correlation between the spatial profiles of cluster 1 and cluster 2 SEs is *r* = 0.708, *P <* 0.001. (B) Brain topographies for the starting modes of cluster 1 SEs as shown in panel A. (C) Brain topographies for the starting modes of cluster 2 SEs as shown in panel A. (D-F) Same as A-C for SEs maximum recruitment modes (see Section 2.7 in Methods). In panel D, the Pearson’s correlation between the spatial profiles of cluster 1 and cluster 2 SEs is *r* =*−* 0.841, *P <* 0.001. (G-I) Same as A-C for SEs ending modes (see Section 2.7 in Methods). In panel G, the Pearson’s correlation between the spatial profiles of cluster 1 and cluster 2 SEs is *r* = 0.718, *P <* 0.001. The reported P values for the statistical significance of the Pearson’s correlation were assessed using Student’s t distributions of the two-tailed hypothesis test under the null hypothesis that the correlation is zero. Symbols and abbreviations: SEs, Salient Events.

### Appendix C.1. Amplitude threshold analysis

The validity and robustness of using a single amplitude threshold (|*z*| = 3) consistently across all 47 participants was investigated as follows. In each participant, the 1-minute source-reconstructed MEG time series of each brain region were first individually z-scored and then concatenated across all brain regions. Subsequently, we computed the histogram and estimated the empirical Probability Density Function (empirical PDF) corresponding to the amplitude values of the concatenated time series (see blue curves in Figs. C.7A and C.7B). Next, we compute the Gaussian distribution that best fit the empirical PDF within each of the 100 fitting intervals of amplitude values spanning the range [*Q*_1_(*z*) *−* 5 ∗ *IQR*(*z*), *Q*_3_(*z*) + 5 ∗ *IQR*(*z*)], where *Q*_1_, *Q*_3_, and *IQR* denote the first quartile, the third quartile and the interquartile range, respectively. This procedure yielded 100 Gaussian PDFs (see grey lines in Fig. C.7A). After that, we computed the RMS error between the empirical PDF and each of the 100 Gaussian PDFs. Where the RMS error was computed using a weighted difference to assign less importance to the difference in the tails of the distributions. As a result of this procedure, we obtained 100 RMS values (see Fig. C.7C). Finally, the optimal threshold for each participant was computed as half the fitting interval of amplitude values producing the minimum RMS error (see Fig. C.7B and the red arrow in Fig. C.7C). Note that the minimum RMS error is associated with the amplitude value (optimal threshold) beyond which the empirical PDF significantly departs from the (best fitted) Gaussian distribution. This procedure was applied separately to all the 47 participants included in the study (see Fig. C.8). The mean and standard deviation of the amplitude thresholds corresponding to the true MEG data shown in Fig. C.8 are 3.08 *±* 0.23. Importantly, the amplitude threshold used in this study (| *z*| = 3) lies approximately at the center of this range. The procedure described above for identifying the optimal amplitude threshold, based on minimizing the RMS error between the empirical PDF and the Gaussian PDFs, was also applied to one A-surrogate and one B-surrogate generated for each participant (see Fig. C.8). The mean and standard deviation of the *z* thresholds across participants were 5 *±* 0.23 for the A-surrogates and 4.7 *±* 0.58 for the B-surrogates, respectively. Of note, the |*z*| thresholds for the A- and B-surrogates were substantially higher than those for the true MEG data. This result is consistent with the fact that the phase randomization applied in the construction of A- and B-surrogates produces approximately Gaussian signals [51].

**Figure C.7:**
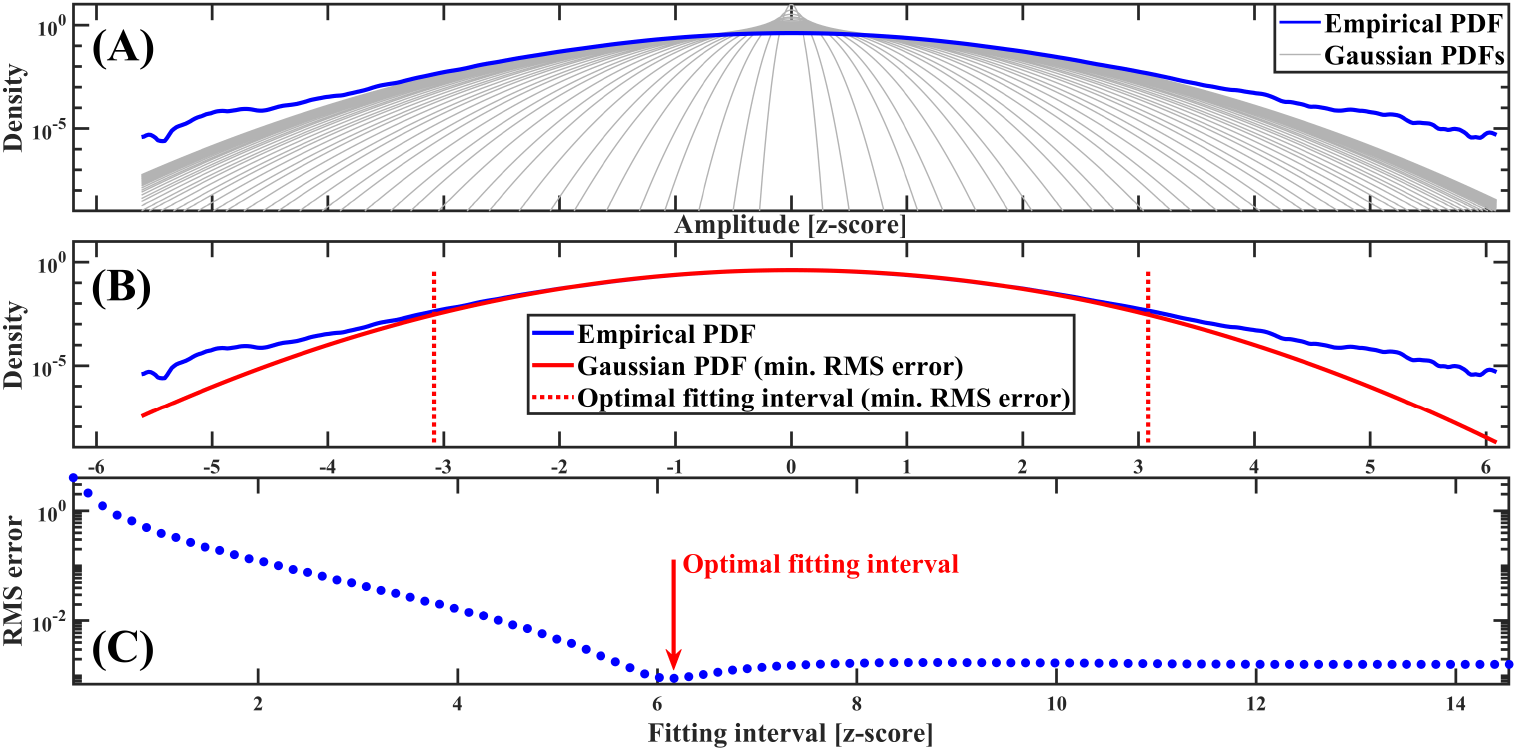
Procedure to find the optimal |*z*| threshold for Participant 47. (A) Empirical and the 100 Gaussian PDFs corresponding to the 100 fitting |*z*| intervals. (B) Empirical PDF together with the Gaussian PDF producing the minimum RMS error. (C) RMS error between the empirical PDF and each of the 100 Gaussian PDFs. Symbols and abbreviations: SNE, Salient Network Event; PDF, Probability Density Function; RMS, Root Mean Square.

**Figure C.8:**
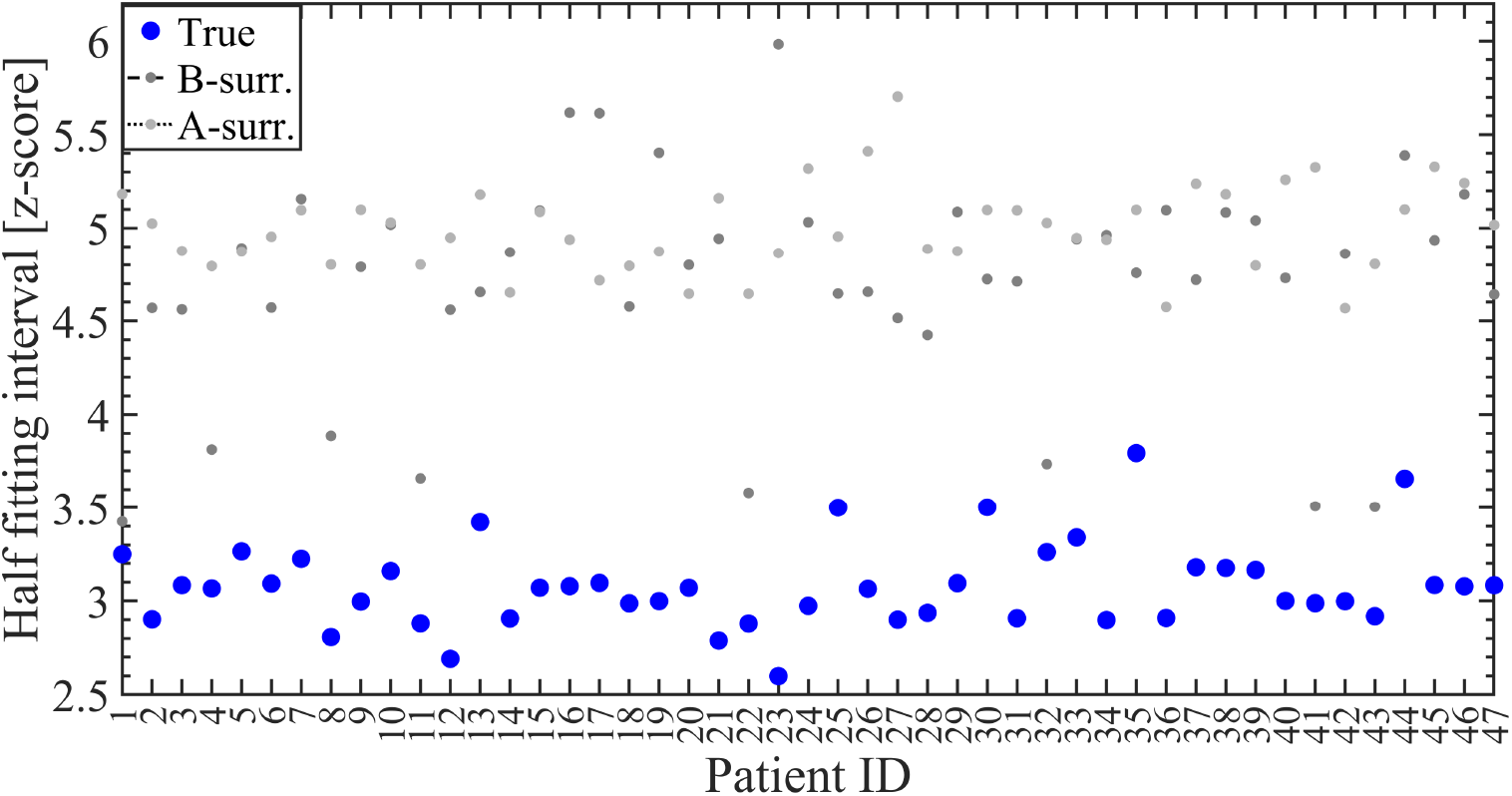
Optimal |*z*| thresholds for the 47 participants corresponding to the true MEG data including the deep sources, and one A-surrogate and one B-surrogate generated for each participant. The mean and standard deviation of the amplitude thresholds corresponding to the true MEG data (blue circles) are 3.08 *±*0.23. Symbols and abbreviations: MEG, Magnetoencephalography.

One of the main limitations of this study is related to the uncertain capability of our dataset to accurately identify deep brain sources along the cortical surface, mainly due to the ill-posed nature of the source-reconstructed MEG data. In order to address this issue, we re-computed the thresholding analysis presented above, but this time excluding the deep sources (see brain topographies in Figs. D.1F and D.3). The results are shown in Fig. C.9. It was found that the mean and standard deviation of the amplitude thresholds corresponding to the true MEG data excluding the deep sources are 3.08 *±* 0.24. Importantly, the amplitude threshold used in this study (| *z*| = 3) lies approximately at the center of this range. Besides, the mean and standard deviation of the |*z*| thresholds across participants were 5*±* 0.23 for the A-surrogates and 4.68*±* 0.59 for the B-surrogates, respectively. As a result, by comparing Figs. C.8 and C.9 we can conclude that the optimal *z* thresholds remain essentially unaltered across the 47 participants when the deep sources are excluded from the thresholding analysis in our MEG dataset.

**Figure C.9:**
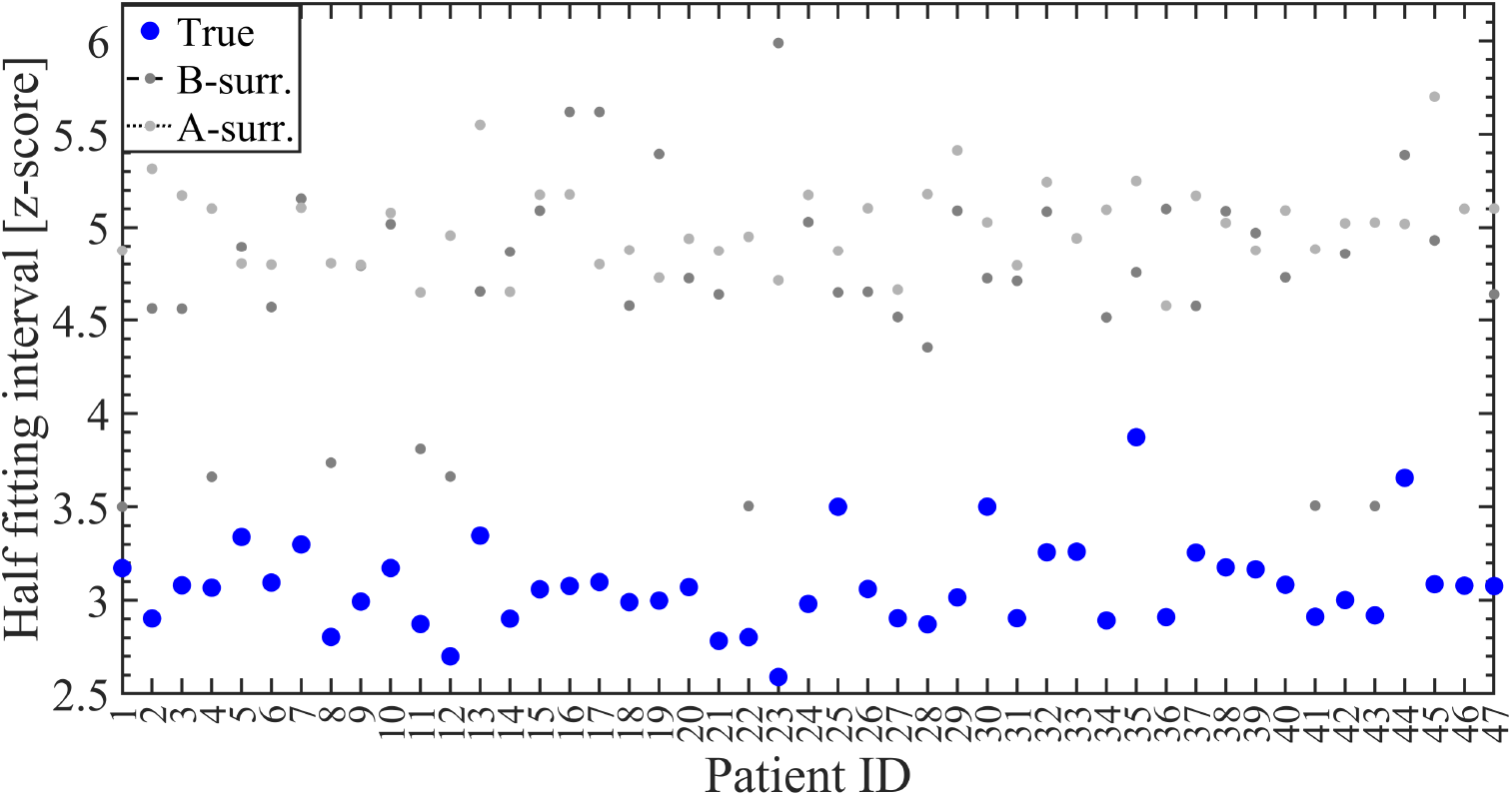
Optimal |*z*| thresholds for the 47 participants corresponding to the true MEG data excluding the deep sources, and one A-surrogate and one B-surrogate generated for each participant. The mean and standard deviation of the amplitude thresholds corresponding to the true MEG data (blue circles) are 3.08 *±*0.24. Symbols and abbreviations: MEG, Magnetoencephalography

### Appendix C.2. Spectral group delay consistency, transient cross-regional coherent NOs and BAA underlie SNEs

In Sections 3.4 and 3.5, we showed that the concurrent presence of BAA and NOs disclosing appropriate levels of SGDC, are two key ingredients sufficient to generate realistic above-threshold fluctuations in a single brain signal (i.e., SLEs). Importantly, we have analytically and computationally shown that only the consistency of the Fourier incremental phase values across frequencies (SGDC) provides a quantitative measure of the level of salience of the above-threshold fluctuations exhibited by the signal in the time-domain, and this relationship holds true regardless of the spectral leakage introduced by tapering in the time-domain (see Fig. B.2). In this section, we present empirical evidence supporting the theoretical findings described in Sections 3.4 and 3.5. Fig. C.10A shows the topography of the mean number of salient (above-threshold) samples assessed in each brain region. The Panels B and C of Fig. C.10 show, respectively, the time series and distributions of the amplitude values corresponding to the brain regions disclosing the maximum (Left supramarginal) and minimum (Left superior frontal) number of salient samples. Importantly, the scatter plots in Panels D and E of Fig. C.10 show a significant correlation between the topographies of the salient samples (Panel A) and, respectively, the *SGDC*(*r*) magnitude and kurtosis. This empirical evidence, together with the results shown in Figs. B.1, B.2 and B.3, further supports the interpretation of the *SGDC*(*r*) as a measure capturing the signal-level mechanism underlying the emergence of local above-threshold fluctuations.

Next, we present the rationale and results pointing out that SGDC is a key conceptualization also in connection with the emergence of realistic SNEs as collective phenomena involving multiple brain regions. Although the *SGDC*(*r*) measure assesses the emergence of local above-threshold fluctuations from the Fourier oscillatory constituents of the activity in a single brain region (i.e., SLEs), it does not account for cross-regional effects associated with SNEs. To quantitatively study the cross-regional effects of SGDC on our data we introduce the *SGDC*(*ω*) measure. The magnitude of *SGDC*(*ω*) is bounded in the range [0, 1] and quantifies how much the group delay at a given frequency *ω* varies across brain regions (Eq. 2). By using synthetic time series, in Appendix A.3 we show that the *SGDC*(*ω*) measure assesses the contribution of each frequency component in the co-activation (synchronization in time) of above-threshold fluctuations across brain regions (see Figs. A.5 and A.6). Of note, Figs. A.5 and A.6 show that the *SGDC*(*ω*) measure effectively resolves the cross-regional synchronization of SEs across frequency bands, whereas phase coherence measures (e.g., PLV: Phase Locking Value) are completely blind to this effect. Then, we used the *SGDC*(*ω*) measure to analyze the two SE clusters observed in our empirical MEG data. Figs. C.11A,B show the average ESMs of the two SE clusters identified by the Louvain algorithm (see Methods) computed on 10 subjects. As shown in Fig. C.11C, only cluster 2 SEs are associated |*SGDC*(*r*)| values higher than those disclosed by the C-surrogate SEs. Importantly, Fig. C.11D shows the increase of transient cross-regional coherence around the alpha band, as quantified by the *SGDC*(*ω*) measure, associated with the SEs disclosing the alpha spectral signature in the average ESM (i.e., cluster 2 SEs). These results are further evidence pointing out that the cluster 2 SEs observed in our MEG data co-occur with (or are coupled to) alpha bursts propagating across brain regions. Notably, Fig. C.11E shows that the transient cross-regional coherence around the alpha band associated with the cluster 2 SEs is also captured by the large-scale model presented in Section 3.5.

Next, we used the *SGDC*(*ω*) measure to analyze the surrogate data computed via phase randomization. Our empirical results show that despite preserving both the power spectrum (PSD) in each brain region and the cross-correlations (i.e., functional connectivity) B-surrogates fail to account for the SEs observed in our MEG dataset. Besides, A-surrogates, which only preserve the regional PSD, perform worst than B-surrogates in reproducing realistic SEs (see Figs. 1 and C.3). The analytical derivations presented in Appendix A.4 provide a unifying rationale for this evidence by pointing out that, on one hand, A-surrogates destroy both the burstiness of each brain region as assessed by the *SGDC*(*r*) measure and the synchronization of above-threshold fluctuations across brain regions as assessed by the *SGDC*(*ω*) measure (see Figs. A.7A,B). On the other hand, B-surrogates significantly reduce the SGDC across frequency components (*SGDC*(*r*), see Fig. A.7A), while preserving the SGDC across brain regions (*SGDC*(*ω*), see Fig. A.7B).

In summary, these results suggest that a) spectral group delay consistency in specific narrow frequency bands (as assessed by the *SGDC*(*r*) measure), b) transient cross-regional coherent NOs (intra-frequency coherence across brain regions assessed by the *SGDC*(*ω*) measure) and c) BAA, are all key ingredients for the emergence of realistic SEs.

**Figure C.10:**
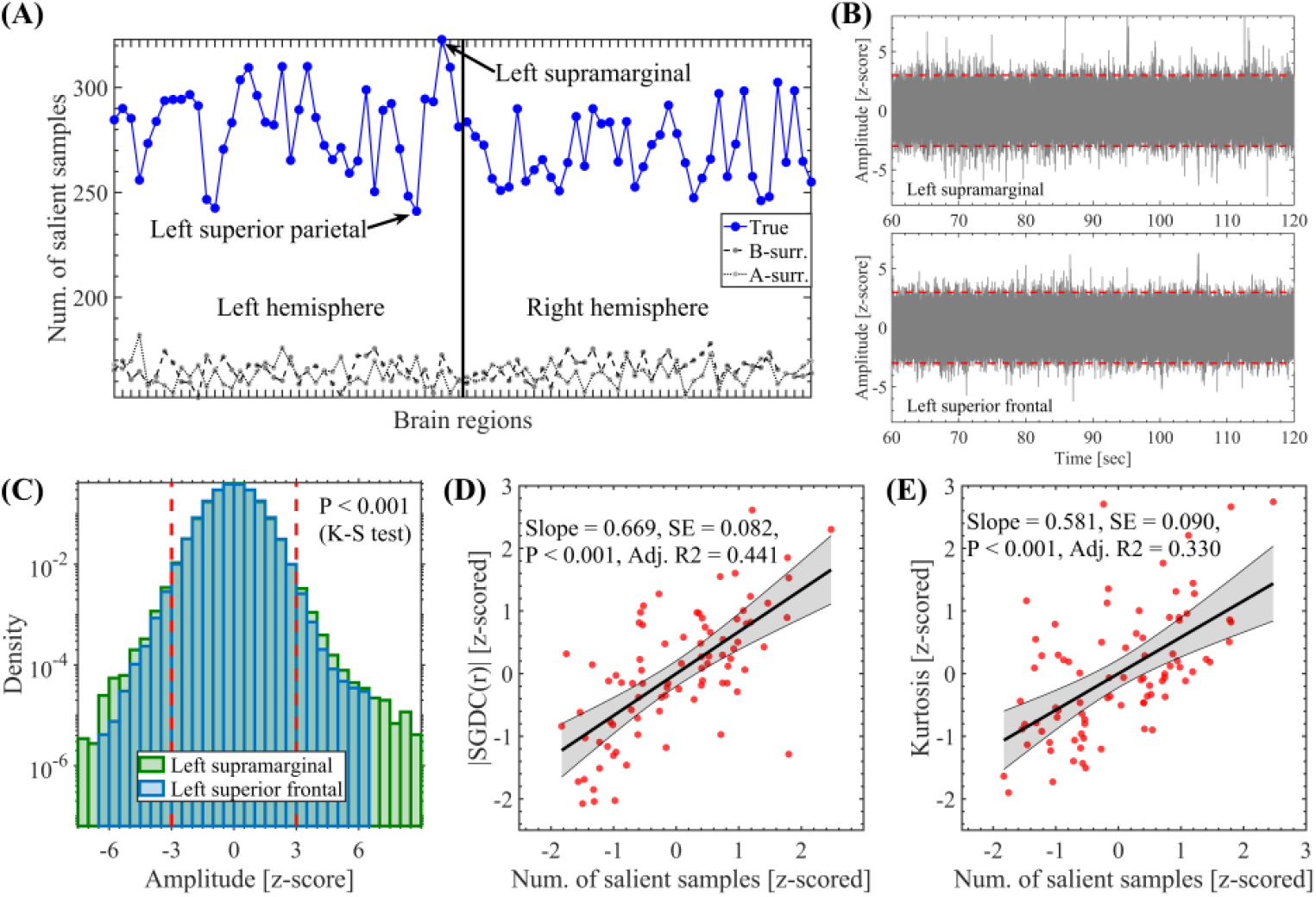
Measures capturing the salient samples topographies. (A) Topography showing the number of salient samples computed on the whole time series (1 min in duration) of each brain region (mean value across the 47 participants). (B) Time series corresponding to the brain regions disclosing the maximum (Left supramarginal) and the minimum (Left superior frontal) number of salient samples. Each plot shows the time series superimposed across the 47 participants. (C) Distributions of the amplitude values for the Left supramarginal and Left superior frontal time series concatenated the 47 participants. Two-sample Kolmogorov-Smirnov test: *P <* 0.001. (D) Scatter plot showing the correlation between the topographies associated with the salient samples and the magnitude of the *SGDC*(*r*) measure. Number of samples (red circles) = Number of brain regions = 84. (E) Same as in (D) for the kurtosis. Symbols and abbreviations: SGDC, Spectral Group Delay Consistency, K-S, Kolmogorov-Smirnov.

**Figure C.11:**
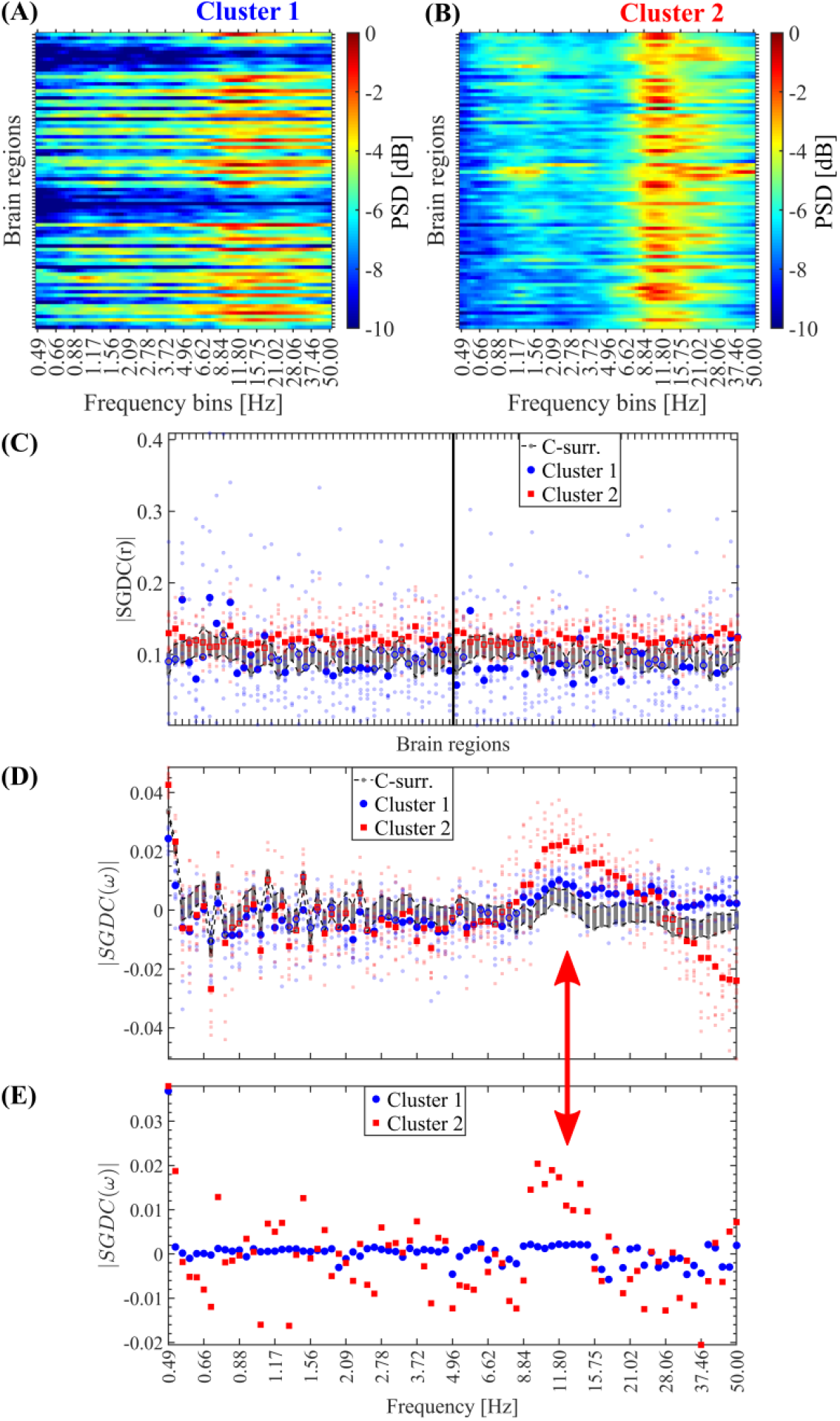
Transient cross-regional coherence around the alpha band is mainly associated with salient events. (A, B) Mean ESM of the two SE clusters identified by the Louvain algorithm computed on the SEs detected in the 10 participants. (C) Transient cross-frequency coherence quantified by the *SGDC*(*r*) measure (see Appendix A.3), associated with the two SE clusters shown in panels A and B. The *SGDC*(*r*) measure was computed in a time-resolved manner. That is, the *SGDC*(*r*) measure was computed on each detected SE by considering the brain regions and time interval associated with each particular event. Then, the *SGDC*(*r*) array was averaged selectively across the SEs segregated in the two clusters produced by the Louvain algorithm (see Section 2.9 in Methods). The small markers represent mean |*SGDC*(*r*)| values averaged across the SEs in each individual participant. The big markers represent mean |*SGDC*(*r*)| values averaged across the 10 participants. (D) Same as in C for the transient cross-regional coherence quantified by the *SGDC*(*ω*) measure (see Appendix A.3). (G) Same as in D for the synthetic data corresponding to the large-scale signal model (see Section 3.5). The red arrow highlight th4e2increase of the |*SGDC*(*ω*)| values around the alpha band. Symbols and abbreviations: SEs, Salient Events; ESM, Event Spectral Matrix.

## Appendix D. Supplementary empirical results excluding the deep sources

**Figure D.1:**
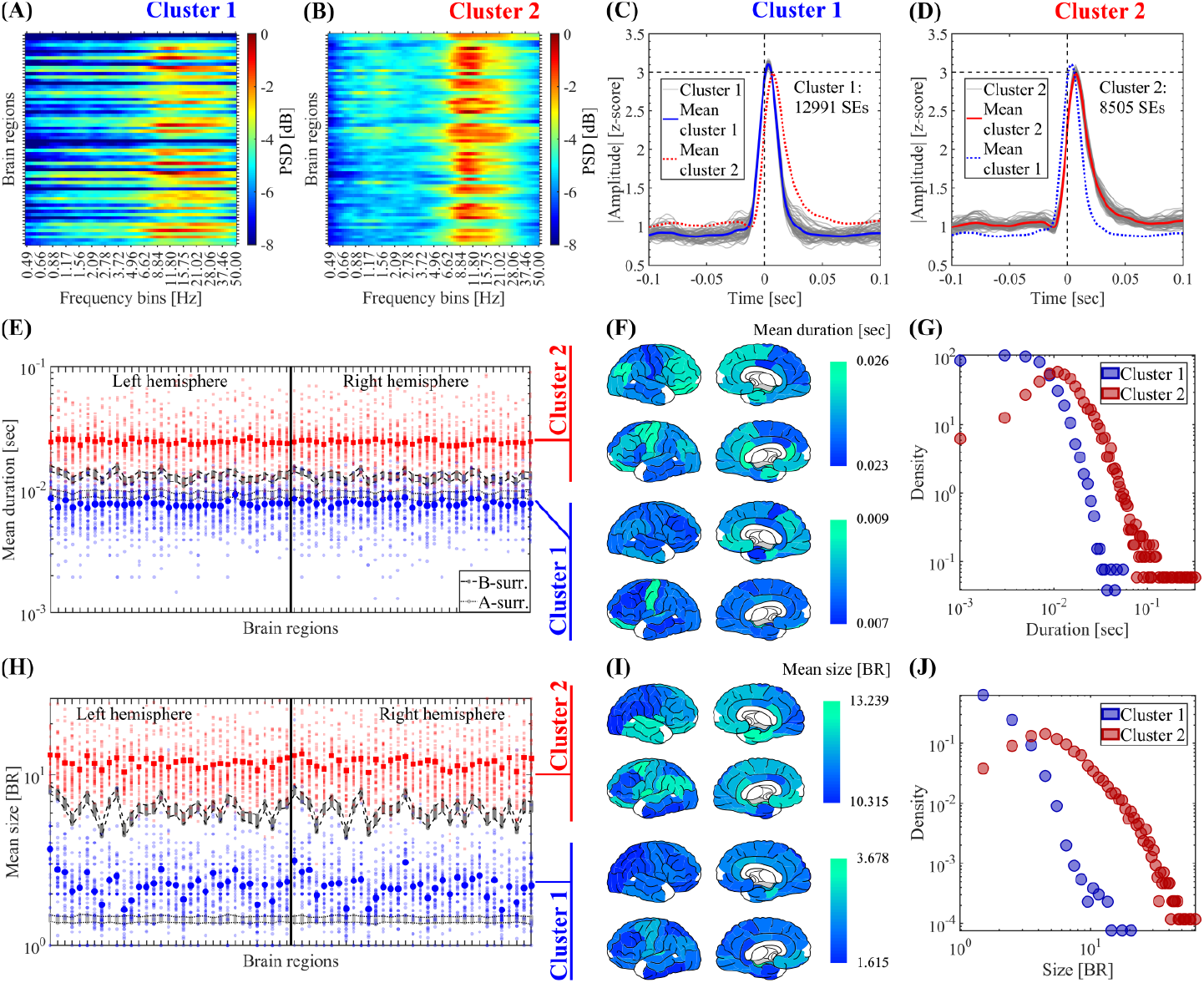
Clustering of SEs according to their spectral signature. The SEs obtained from 45 subjects were clustered using the Louvain algorithm (resolution parameter *γ* = 1, see Methods). (A, B) Mean ESM of the two SE clusters identified by the Louvain algorithm computed on the SEs detected in the 45 participants. (C, D) Waveform shapes of the SEs pertaining to the two SE clusters identified by the Louvain algorithm. Thin gray lines correspond to the average waveform shape in each brain region. Thick blue and red lines correspond to the resulting waveform shape averaged across the brain regions for cluster 1 and 2 SEs, respectively. (E) Spatial profile showing the mean duration of SEs pertaining to cluster 1 (in blue) and cluster 2 (in red). For the true data, the small and big markers correspond to the mean spatial profile in each patient and the average across the 45 participants, respectively (see Methods). The labels and ordering of the brain regions are the same as those shown in Fig. C.2. To test the significance of the difference of the mean SEs duration between cluster 1 and cluster 2, in each brain region we computed a non-parametric permutation test (random sampling without replacement, 1 *×* 10^4^ permutations). All the brain regions disclosed a statistically significant difference of the mean SEs duration between cluster 1 and 2 (the Bonferroni-adjusted two-tailed P values result *P <* 0.001 in all the brain regions). (F) Brain topographies for the mean duration of SEs averaged across the 45 participants as shown in panel E. (G) Distribution of the duration of SEs pertaining to the cluster 1 and cluster 2 observed in the 45 participants. (H) Same as in E for the size of SEs. To test the significance of the difference of the mean SEs size between cluster 1 and cluster 2, in each brain region we computed a non-parametric permutation test (random sampling without replacement, 1*×* 10^4^ permutations). All the brain regions disclosed a statistically significant difference of the mean SEs size between cluster 1 and 2 (the Bonferroni-adjusted two-tailed P values result *P <* 0.001 in all the brain regions). (I) Same as in F for the size of SEs. (J) Same as in G for the size of SEs. Symbols and abbreviations: SEs, Salient Events; ESM, Event Spectral Matrix; BR, Brain Regions.

**Figure D.2:**
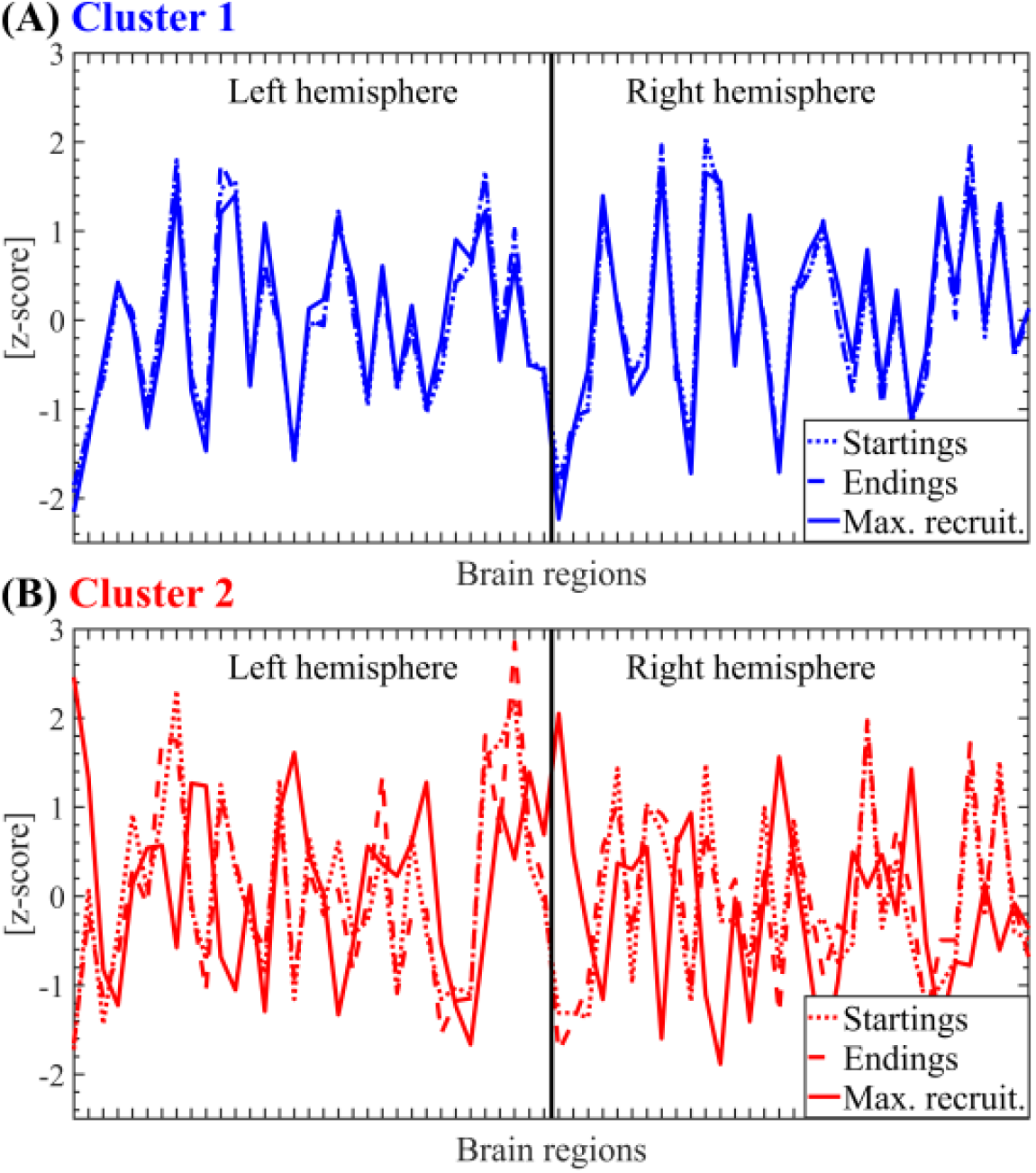
Salient events propagation modes segregated by SE clusters. (A) Spatial profile for the cluster 1 SEs starting, maximum recruitment and ending modes (see Section 2.7 in Methods) computed on 45 participants. Linear correlations between topographies: Startings vs Endings, *r* = 0.995, *P <* 0.001. Max. recruit. vs Startings, *r* = 0.972, *P <* 0.001. Max. recruit. vs Endings, *r* = 0.968, *P <* 0.001. (B) Same as in A for the cluster 2 SEs starting, maximum recruitment and ending modes. Linear correlations between topographies: Startings vs Endings, *r* = 0.917, *P <* 0.001. Max. recruit. vs Startings, *r* =− 0.052, *P* = 0.7. Max. recruit. vs Endings, *r* = −0.051, *P* = 0.7. The SEs obtained from 45 subjects were clustered using the Louvain algorithm (resolution parameter *γ* = 1, see Section 2.9 in Methods). The reported P values for the statistical significance of the Pearson’s correlation were assessed using Student’s t distributions of the two-tailed hypothesis test under the null hypothesis that the correlation is zero. Symbols and abbreviations: SEs, Salient Events.

**Figure D.3:**
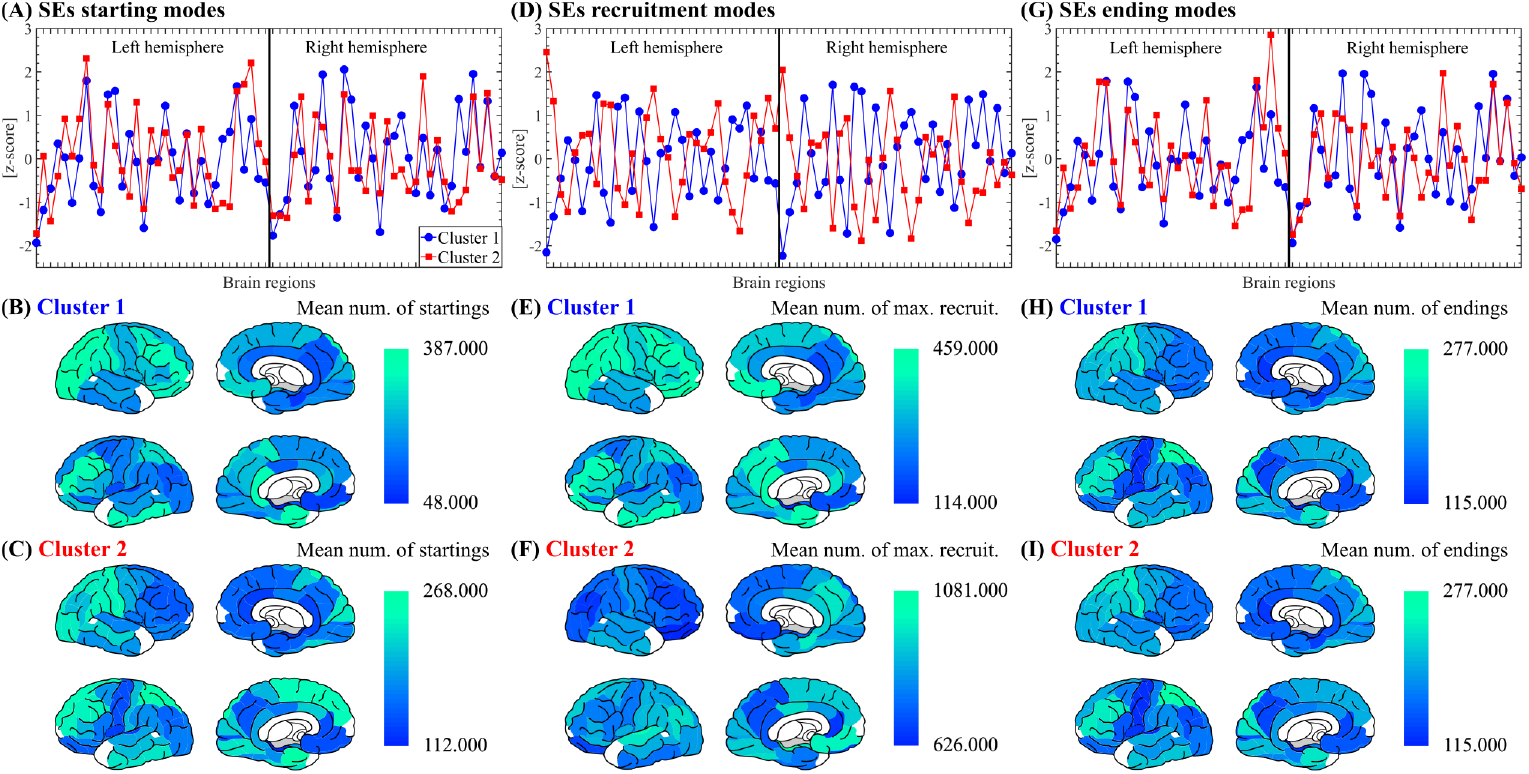
Salient events propagation modes. (A) Spatial profile for the SEs starting modes (see Section 2.7 in Methods) corresponding to the two SE clusters computed on 45 participants. The SEs obtained from 45 subjects were clustered using the Louvain algorithm (resolution parameter *γ* = 1, see Section 2.9 in Methods). The Pearson’s correlation between the spatial profiles of cluster 1 and cluster 2 SEs is *r* = 0.584, *P <* 0.001. (B) Brain topographies for the starting modes of cluster 1 SEs as shown in panel A. (C) Brain topographies for the starting modes of cluster 2 SEs as shown in panel A. (D-F) Same as A-C for SEs maximum recruitment modes (see Section 2.7 in Methods). In panel D, the Pearson’s correlation between the spatial profiles of cluster 1 and cluster 2 SEs is *r* = −0.842, *P <* 0.001. (G-I) Same as A-C for SEs ending modes (see Section 2.7 in Methods). In panel G, the Pearson’s correlation between the spatial profiles of cluster 1 and cluster 2 SEs is *r* = 0.571, *P* < 0.001. The reported P values for the statistical significance of the Pearson’s correlation were assessed using Student’s t distributions of the two-tailed hypothesis test under the null hypothesis that the correlation is zero. Symbols and abbreviations: SEs, Salient Events.

## Notes

### Competing Interest Statement

The authors have declared no competing interest.

### Summary of Updates

In this version we improved the description of the proposed SGDC framework.

